# Regulation of potassium homeostasis in *Caulobacter crescentus*

**DOI:** 10.1101/2023.07.05.547876

**Authors:** Alex Quintero-Yanes, Loïc Léger, Madeline Collignon, Julien Mignon, Aurélie Mayard, Catherine Michaux, Régis Hallez

## Abstract

Potassium (K^+^) is an essential physiological element determining membrane potential, intracellular pH, osmotic/turgor pressure, and protein synthesis in cells. Nevertheless, K^+^ homeostasis remains poorly studied in bacteria. Here we describe the regulation of potassium uptake systems in the oligotrophic α-proteobacterium *Caulobacter crescentus* known as a model for asymmetric cell division. We show that *C. crescentus* can grow in concentrations from the micromolar to the millimolar range by essentially using two K^+^ transporters to maintain potassium homeostasis, the low affinity Kup and the high affinity Kdp uptake systems. When K^+^ is not limiting, we found that the *kup* gene is essential while *kdp* inactivation does not impact the growth. In contrast, *kdp* becomes critical but not essential and *kup* dispensable for growth in K^+^-limited environments. However, in the absence of *kdp*, mutations in *kup* were selected to improve growth in K^+^-depleted conditions, likely by improving the affinity of Kup for K^+^. In addition, mutations in the KdpDE two-component system, which regulates *kdpABC* expression, suggest that the inner membrane sensor regulatory component KdpD works as a kinase in early stages of growth and as a phosphatase to regulate transition into stationary phase. Our data also show that KdpE is not only phosphorylated by KdpD but also by another non-cognate histidine kinase. On top of this, we determined the KdpE-dependent and independent K^+^ transcriptome and identified the direct targets of KdpE. Together, our work illustrates how an oligotrophic bacterium responds to fluctuation in K^+^ availability.

**Importance:** Potassium (K^+^) is a key metal ion involved in many essential cellular processes. Its transport and regulation have been mainly studied in the bacterial model species *Escherichia coli* and *Bacillus subtilis*. Here we show that the oligotroph *Caulobacter crescentus* can support growth at lower K^+^ concentrations by mainly using two K^+^ uptake systems, the low-affinity Kup and the high-affinity Kdp. Interestingly, in the absence of Kdp, point mutations in Kup was selected to increase affinity for K^+^, which improved growth in K^+^-depleted conditions. Using genome-wide approaches, we also determined the entire set of genes required for *C crescentus* to survive at low K^+^ concentration as well as the full K^+^-dependent regulon. Finally, we found that the transcriptional regulation mediated by the KdpDE two-component system is unconventional since unlike *E. coli*, the inner membrane sensor regulatory component KdpD works rather as a phosphatase on the phosphorylated response regulator KdpE∼P. To our knowledge, this is the first comprehensive study of K^+^ homeostasis in bacteria.

## Introduction

Potassium (K^+^) is the most abundant monovalent cation within living cells (1, 2) and it is essential to set the membrane potential and intracellular pH (2–5). As K^+^ also emerges as an important regulator of bacterial physiology in view of its role in diverse processes (6–17) including biofilm formation, chemotaxis, cell to cell communication or virulence, it is therefore expected that K^+^ transport is tightly regulated. In bacteria, the KdpDE two-component system (TCS) responds to potassium depletion by regulating the expression of the cognate high-affinity K^+^ transporter KdpFABC (18). This inner membrane complex consists of four subunits all encoded in the same operon, often together with the *kdpDE* genes (19, 20). In *Escherichia coli*, KdpD was shown to act as a dual sensor for K^+^, which independently regulates both enzymatic activities. Whereas the kinase activity was inhibited by extracellular K^+^, phosphatase activity was stimulated by intracellular K^+^ (21). This ultimately determines the phosphorylation status of the response regulator KdpE and thus provides a tight regulation of *kdpFABC* expression as well as a robust homeostasis in fluctuating environments (21).

Other K^+^ transporters have been described in detail in Gram-negative bacteria, such as the inner membrane low-affinity Trk and Kup systems for uptake, and the KefB efflux pump. The Trk potassium uptake system is highly conserved in bacteria and works in a multi-subunit complex comprising the TrkG/H, TrkA and TrkE proteins. TrkH (an extra copy known as TrkG is found in *E. coli*) is the trans-membrane transporter unit, which interacts with the NAD^+^- and NADH-binding peripheral protein TrkA and the ATP-binding protein TrkE in the cytoplasmic leaflet to control extracellular potassium uptake when high extracellular concentrations of K^+^ are available (22–24). On the other hand, Kup is a single protein, constitutively expressed and thought to function as K^+^/Na^+^ symporter (23, 25, 26).

Recently, it was reported that glutathione controls cell division via negative regulation of KefB in the Gram-negative oligotrophic α-proteobacterium *Caulobacter crescentus* (27). Indeed, mutants unable to synthesize glutathione led to cell filamentation and decreased intracellular K^+^ levels. Interestingly, these phenotypes were suppressed by mutations in the KefB efflux pump, suggesting that K^+^ homeostasis is crucial for bacterial cell cycle regulation. *C. crescentus* has been extensively used as a model for cell cycle studies since it divides asymmetrically to give birth to a larger sessile stalked cell and a smaller flagellated swarmer cell (28). DNA replication and cell division take place only in sessile stalked cells whereas the swarmer cells remain in a non-replicative but chemotactically active and motile phase able to explore new environments. Once swarmer cells find a suitable – resourceful – habitat for reproduction, they differentiate into stalked cells and concomitantly start DNA replication. In contrast, upon deprivation of environmental resources, the swarmer cells do not differentiate into replicative stalked cells (29).

In this study, we characterized the response of *Caulobacter* cells grown in environments containing what we defined as limiting, abundant and excessive K^+^ concentrations. *In silico* analyses revealed that *C. crescentus* does not have Trk system components but encodes a putative transporter with a cytoplasmic TrkA-like domain. Moreover, in addition to the KefB and KefC efflux pumps, we found genes predicted to code for a Kup and a KdpABC transporters as well as a KdpDE TCS. We generated mutants of these predicted K^+^ transporters and assessed their fitness in limiting, abundant and excessive K^+^ environments. We also characterized the KdpDE system and determined its regulon by ChIP-seq and RNA-seq.

## Results

### Growth of *C. crescentus* at different K^+^ concentrations

To primarily assess the impact of K^+^ on *C. crescentus* fitness, the wild-type (WT) strain was grown in K^+^-free minimal media (M2G-K) supplemented with K^+^ at different concentrations using either KCl or a combination of K_2_HPO_4_ and KH_2_PO_4_ (K_2_HPO_4_ + KH_2_PO_4_) like the one used in standard M2G minimal media (**Fig. 1**). First, when phosphate salts were used as K^+^ source, we observed that compared to M2G, which contains 7 mM K^+^, growth started to be significantly impaired at K^+^ concentrations below or equal to 0.025 mM and above or equal to 7 mM (**Fig. 1AB**). Second, by using KCl, we found different K^+^ concentrations critical for growth since the WT already failed to grow at 0.1 mM but supports growth up to 25 mM (**Fig. 1CD** and **Fig. S1A**). Notwithstanding these differences likely due to the PO4^3-^ and Cl^-^ counter-anions, the growth of the WT at K^+^ concentrations ranging from 0.5 mM to 5 mM was indistinguishable from the one in M2G, that is 7 mM (**Fig. 1**). Therefore, based on these data, we will use in this study potassium phosphate salts at 0.025 mM and 0.5 mM respectively as limiting and abundant concentrations while KCl will be used at 50 mM as excessive concentration.

**Figure 1.**
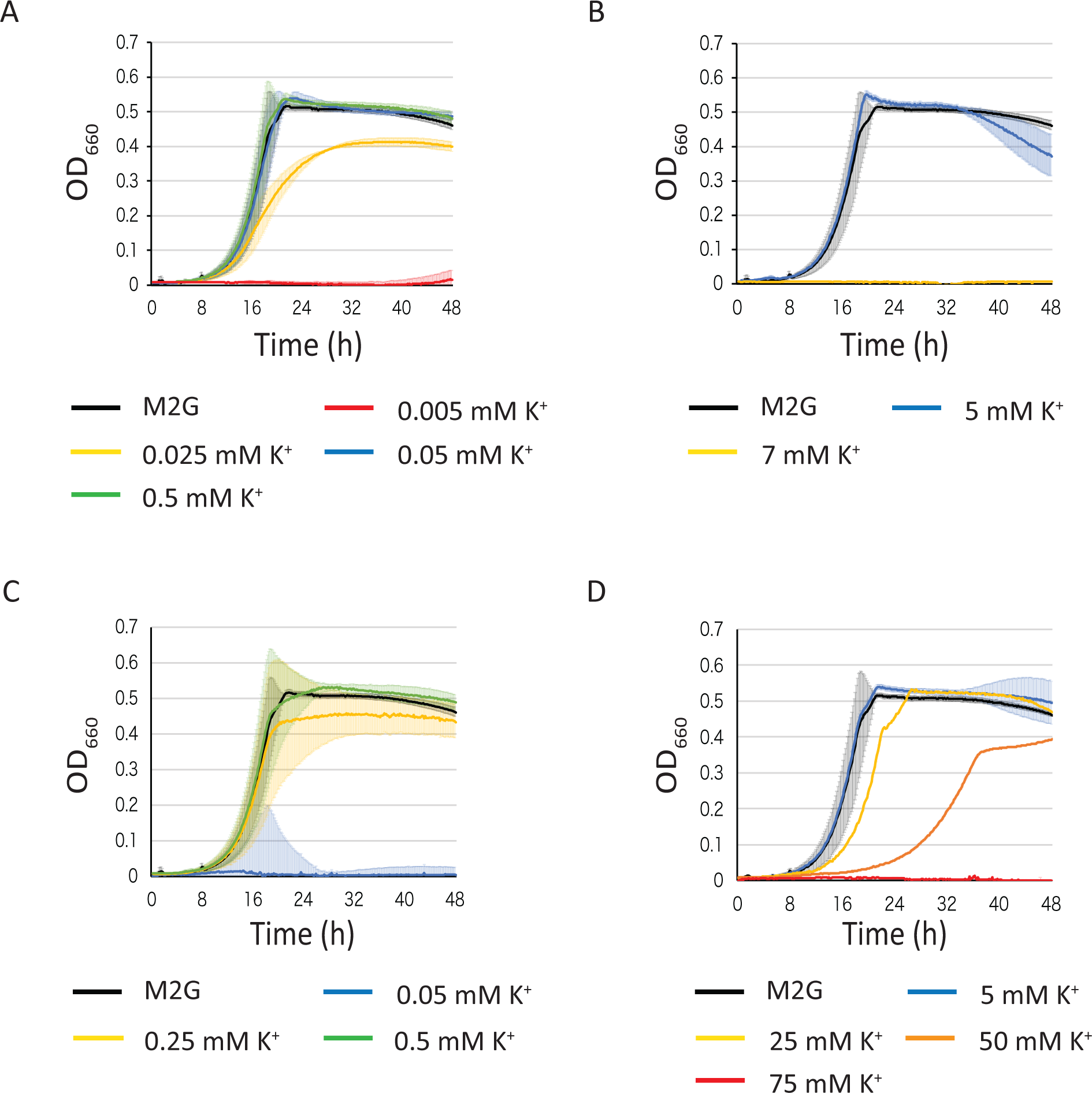
Impact of K^+^ on *C. crescentus* growth. Growth of WT in M2G-K supplemented with different K^+^ concentrations using K_2_HPO_4_ + KH_2_PO_4_ (A-B) and KCl (C-D) as K^+^ source. Data represent average, n=3, and error bars= ±SD. Other concentrations tested are presented in Fig. S1. Flow cytometry analysis was used to determine DNA content throughout the cell cycle of a synchronized population of WT strain grown in M2G-K, 0.025 mM K^+^ and 0.5 mM K^+^ (D). Samples were withdrawn every 20 minutes for 140 minutes (D).

### K^+^ transport and regulation systems in *C. crescentus*

By using *in silico* analyses, we found in the genome of *C. crescentus* NA1000 (NC_011916.1) several genes encoding potential K^+^ uptake (*kup* and *kdpABCDE*) or efflux (*kefB*, *kefC* and *kefG*) systems (**Fig. 2**). Although Trk orthologs were not found, a putative transporter (CCNA_01688) containing a cytoplasmic TrkA-like domain was identified (**Fig. 2**). Based on the function of the Kdp, Kef, Kup and Trk systems in other bacteria (2, 12, 30–32), we predicted that K^+^ transport in *C. crescentus* occurs via CCNA_01688, KefBG, KefC and Kup when potassium is abundant in the environment (**Fig. 2A**) whereas the Kdp system might be inactive (**Fig. 2B**) (33, 34). In contrast, K^+^-depleted conditions should activate the KdpDE TCS and trigger expression of the high affinity KdpABC transporter (**Fig. 2C)** (35).

**Figure 2.**
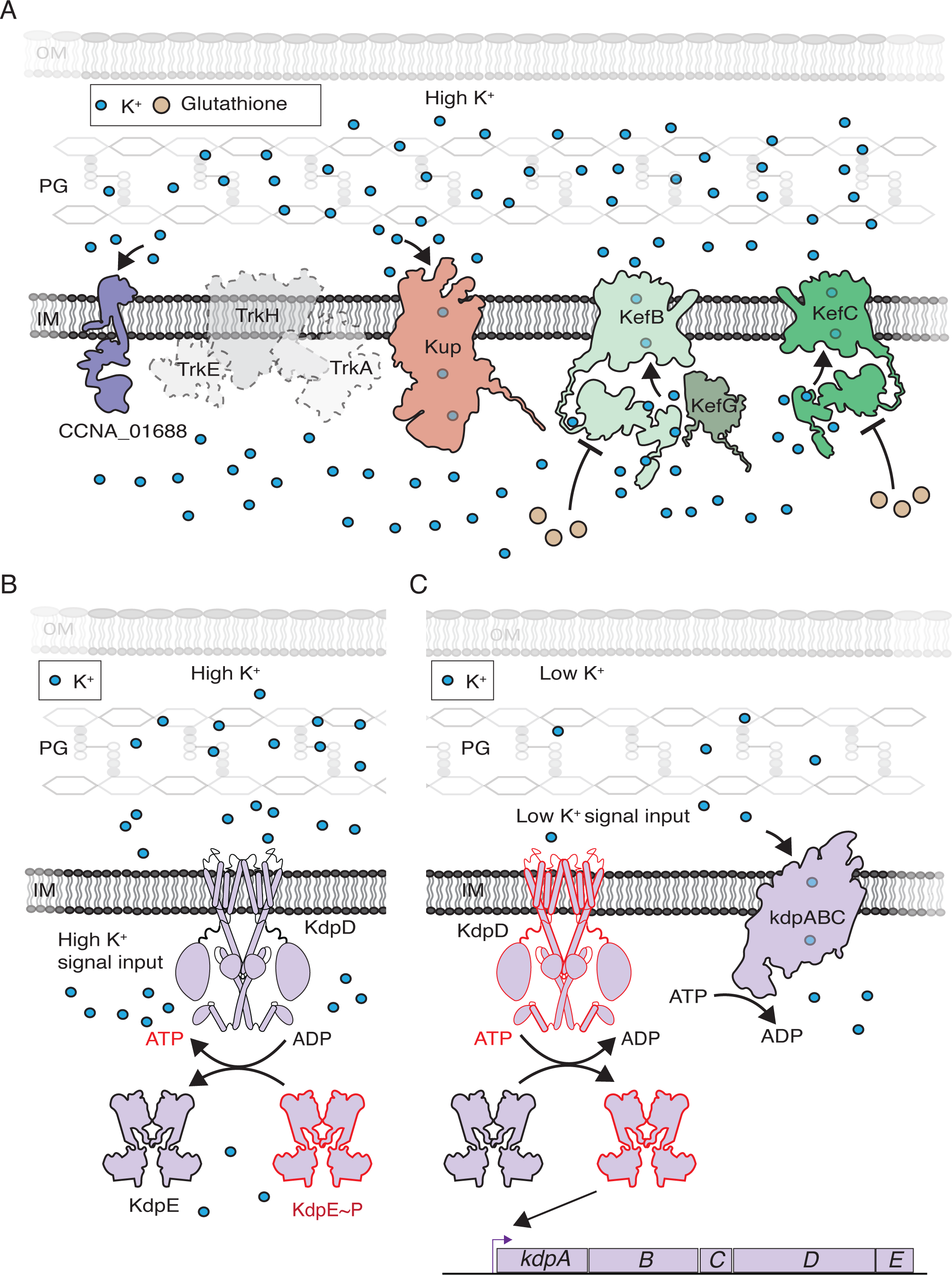
K^+^ transport and response systems in Gram-negative bacteria. Schematic diagram of K^+^-related proteins predicted to work at high (A-B) or low (C) K^+^ concentrations. Coloured proteins correspond to the ones encoded in *C. crescentus* genome: CCNA_01688, Kup (CCNA_00130), KefB (CCNA_00204), KefG (CCNA_00205), KefC (CCNA_03611), KdpDE (CCNA_01666-CCNA_01667) KdpFABC (CCNA_01662-CCNA_01665). The genes were identified using Biocyc collection of microbial genomes (85) and structures of the proteins obtained using predictions with alphafold from Uniprot (86). Outer Membrane (OM), peptidoglycan (PG), inner membrane (IM).

To test these predictions, we first constructed knock-out (KO) mutants for all the K^+^ transport and regulatory systems described above. We successfully inactivated all these genes except *kup*, suggesting it might be essential at least at the K^+^ concentration found in the complex media PYE used to construct the KO strains. Note that a Δ*kefB* and Δ*kdpABCDE* (hereafter referred to as Δ*kdp*) were respectively used as KefBG and Kdp inactive mutants (**Fig. 2C**). Then, we measured the growth for the mutants in M2G-K supplemented with limiting (0.025 mM), abundant (0.5 mM) or excessive (50 mM) K^+^ concentrations (**Fig. 3**). In limiting K^+^ conditions, only Δ*kdp* had a strong growth delay (**Fig. 3A**) whereas all the KO mutants grew as good as the WT, both in minimal and complex media supplemented with abundant K^+^ concentrations (**Fig. 3B** and **Fig. S1B**). However, and as far as we know in contrast to what is described in other bacteria, a *kdp* inactive mutant could still grow in such limiting conditions, suggesting that another transporter is used in K^+^-depletion conditions, at least when *kdp* is inactive. At excessive K^+^ concentration, we found that (i) Δ*CCNA_01688* had a slight growth delay; (ii) Δ*kdp* had a lower plateau and (iii) Δ*kefC* barely grew in comparison to WT (**Fig. 3C**). Interestingly, the growth delay of Δ*CCNA_01688* and the lower plateau of Δ*kdp* were not observed in complex PYE media supplemented with an excess of K^+^ whereas Δ*kefC*, like in minimal medium, did not grow (**Fig. S1C**).

**Figure 3.**
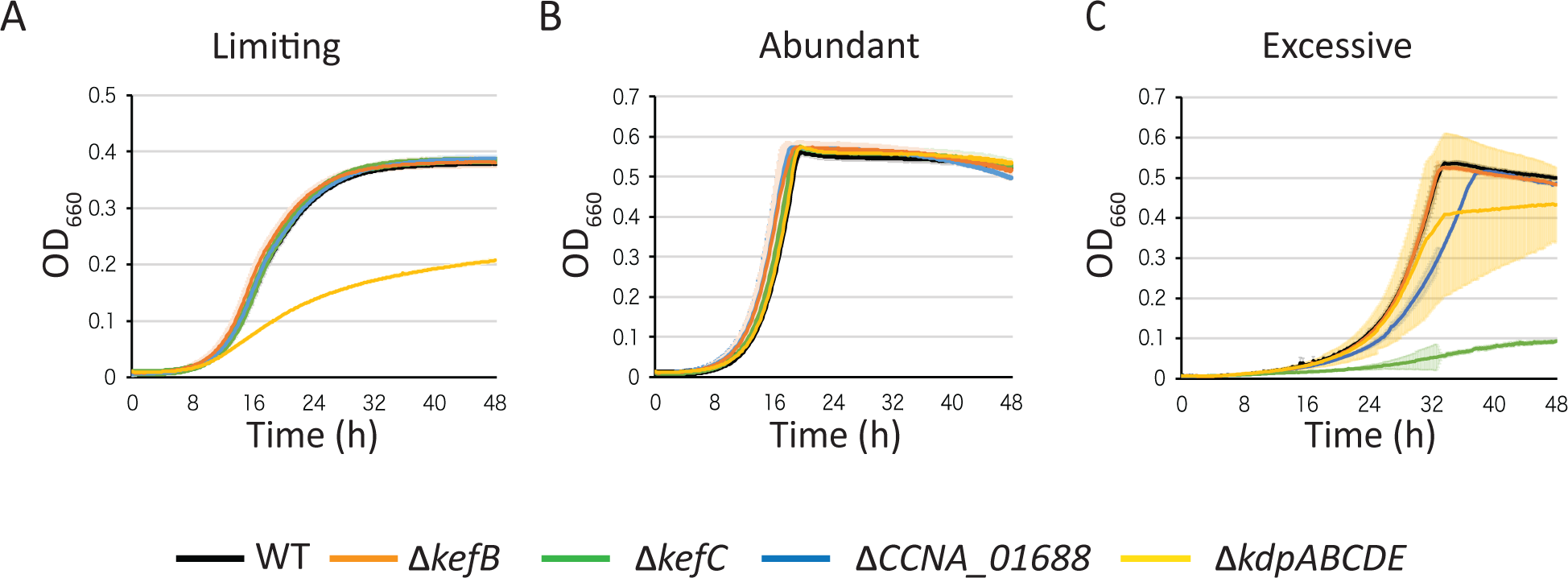
Deletion of K^+^-related genes impact *C. crescentus* growth. Growth of WT, Δ*kefB*, Δ*kefC*, ΔCCNA_01688 and Δ*kdpABCDE* mutants in minimal media M2G-K supplemented with (A) limiting (0.025 mM), (B) abundant (0.5 mM) and (C) excessive (50 mM) K^+^ concentrations. Data represent average, n=3, and error bars= ±SD.

### K^+^-dependent essential genome

Transposon mutagenesis coupled to next-generation sequencing (Tn-seq) were used to determine a core set of genes important for adaptation and response to abundant or limiting K^+^ condition. Isolated colonies harbouring transposon (Tn) insertions were grown on and collected from solid agar plates (**Fig. 4A**). We used M2G-K with 1 % agar concentration to lower as much as possible K^+^ traces from the agar in our experiment. Indeed, even at 1% agar, the K^+^ traces were sufficient to allow WT cells to grow without exogenous source of K^+^ (on M2G-K, **Fig. 4B**). In contrast, at least 0.025 mM K^+^ had to be supplemented to M2G-K to allow growth of Δ*kdp* while at 0.5 mM K^+^, both the mutant and the WT grew similarly to the growth observed on regular M2G which contains 7 mM K^+^ (**Fig. 4B**). Therefore, we decided to use M2G-K without exogenous K^+^ and M2G plates respectively as the limiting and abundant K^+^ conditions for the Tn-seq (**Fig. 4B**).

**Figure 4.**
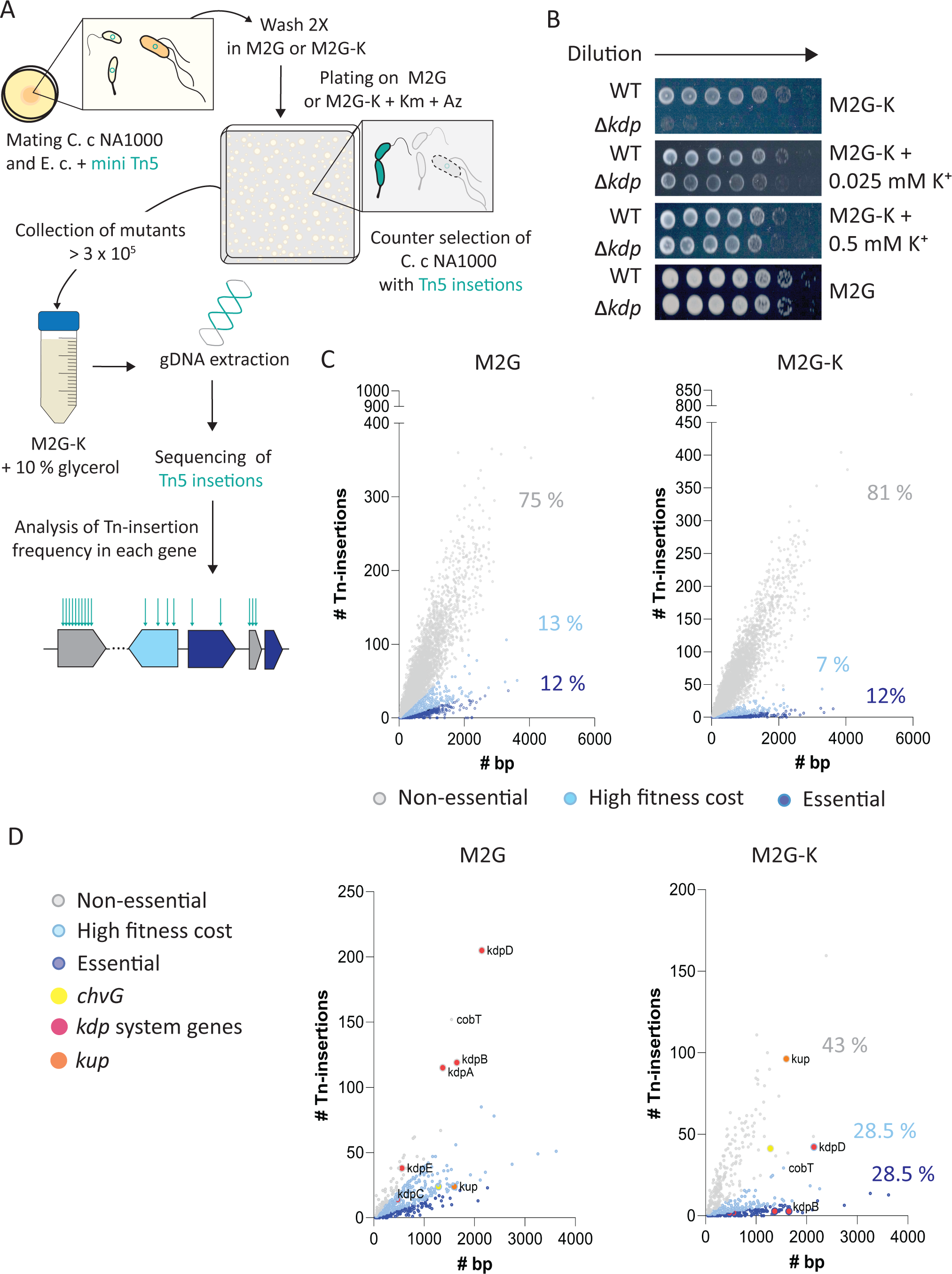
Tn-seq and RNA-seq profile of *C. crescentus* cells in limiting K^+^ conditions. (A) Diagram of the Tn-seq experiment. As explained in detail in Material and Methods section, *E. coli* (E. c) cells were used to deliver a mini Tn5 vector into *C*. *crescentus* (C. c) by conjugation. Mating spots were washed twice in either M2G or M2G-K. Cells were plated in either M2G or M2G-K square plates supplemented with kanamycin (Km) and aztreonam (Az) to counter select *C. crescentus* cells with Tn5 insertions (green cells) over *C. crescentus* without insertions and *E. coli* cells. Cells were collected to complete > 3 x 10^5^ clones. Thereafter gDNA extraction and Tn5 directed sequencing was performed to determine the frequency of insertions for each gene in *C. crescentus* genome and determine genes that are essential, high-fitness cost or non-essential in M2G and M2G-K plates. (B) Growth of WT and Δ*kdp* cells in M2G-K plates without or supplemented with K^+^. (C) Number of Transposon insertions (#Tn insertions) against the length (#bp) of 4186 genes in M2G and M2G-K media. Non-essential genes are highlighted in grey, high-fitness cost genes in light blue, and essential in dark blue. (D) Representation of genes that changed in their fitness-cost categories according to analysis done in M2G and M2G-K (C). (C-D) The percentage of genes in each category is indicated following the colour code.

We did calculations on the ratio of Tn insertions per base pair (# Tn insertions / bp) for internal 80% of each ORF in each condition (M2G and M2G-K) (**Table S1**). As in studies performed in other bacteria, we analysed the frequency of Tn insertions and observed a bimodal distribution that allowed us visualizing two clusters of genes in each condition, one with low density of Tn insertions (considered as essential genes) and the other one with a higher density (considered as non-essential genes) (**Figure S2A)**. Based on this frequency of Tn insertions, we used Ward’s clustering analysis (36) and defined three fitness cost categories, that are (i) essential, (ii) high fitness cost (HFC) and (iii) non-essential (**Fig. 4A**). For the M2G library, genes with values of Tn insertions / bp ≤ 0.01333333 were considered as essential, 0.01333779 ≤ and ≤ 0.04063018 as HFC, and ≥ 0.04081632 as non-essential. For the M2G-K library, genes values of ≤ 0.0047112 were considered as essential, 0.0047909 ≤ and ≤ 0.0211178 as HFC, and ≥ 0.0215792 as non-essential (**Figure 4C** and **Table S1**). Among the 4186 annotated genes in *C. crescentus*, the percentage of essential genes in both media (M2G and M2G-K) was similar (∼12 %), whilst the percentage of HFC genes was lower (∼7 %) in M2G-K than in M2G (∼13 %). Hence, the percentage of non-essential genes was higher in M2G-K (∼ 81 %) than in M2G-K (∼75 %). The shift from the HFC to the non-essential category in *C. crescentus* cells grown in M2G-K could be explained by the reduction of osmotic stress in this medium. Indeed, we showed recently that M2G generates a cell envelope stress for *C. crescentus* due to the high content of monovalent cations (37).

Genes with known or predicted biological functions were grouped according to the classification of cluster of orthologues groups (COG) (**Fig S2B**). This allowed to highlight biological functions that are enriched in abundant and limited K^+^ conditions. Overall, the COG enrichment in the three fitness categories (Essential, High Fitness Cost – HFC and Non-Essential) was substantially similar in both M2G and M2G-K media. The C group “energy production and conversion” genes was the most abundant for all the fitness categories in M2G-K, as well as for essential and non-essential genes in M2G, while for HFC genes it was the second most abundant, after the J group “translation, ribosomal structure and biogenesis”.

By comparing the gene pool in each category for each media, we identified a set of 537 genes that changed their fitness cost from M2G to M2G-K and vice versa (**Fig. 4D** and **Table S2**), including 153 genes which became exclusively essential (28.5 %), 153 HFC (28.5 %) and 231 non-essential (43 %) in M2G-K. First, we observed that the TCS genes *chvGI*, known to be HFC in M2G due to its response to hyperosmotic stress (37), became non-essential in M2G-K, thereby supporting that M2G-K is a hypo-osmotic environment compared to M2G. Second, all the *kdp* genes categorised as non-essential in M2G became essential in M2G-K, except *kdpD* which became HFC. In contrast, *kup* moved from the HFC category in M2G to non-essential one in M2G-K. This confirms the importance of the Kdp and Kup systems respectively in limiting and abundant K^+^ conditions. Considering the conservation of the *kdp* system in bacteria, it is intriguing that *kdpD* – encoding the HK component – did not have the same fitness cost in M2G-K than the genes coding for the transporter (*kdpABC*) and the response regulator (*kdpE*) components. While this supports that KdpE likely regulates, directly, the expression of *kdpABC* genes, it suggests that KdpD and KdpE have different regulatory effects on growth in K^+^-depleted environments. Finally, we did not observe changes in fitness for the other predicted potassium transport systems since all of them were categorised as non-essential genes in both M2G and M2G-K, further supporting that CCNA_01688, KefBG and KefC are not important for growth in abundant and limiting K^+^ conditions.

### Kup becomes dispensable in limiting K^+^ conditions

Tn-seq data suggest that inactivation of *kup* gene could be facilitated at limiting K^+^ concentrations (**Fig. 4C** and **D**, **Tables S1** and **S2**). Kup has been described in different bacteria as the major K^+^ transporter, essential in hyperosmotic conditions using sucrose as osmotic stressor (38, 39). Moreover, in a previous study, we found that the TCS ChvGI in *C. crescentus* positively controls the expression of *kup* in hyperosmotic conditions (37). In agreement with this, we could not obtain a Δ*kup* strain by using a markerless, SacB-dependent recombination approach on complex media (PYE) plates supplemented with 3% sucrose for the counterselection. Since addition of sucrose considerably increases the osmolality of the media (40), we aimed to construct a Δ*kup* mutant by lowering osmolytes in the media, that is on M2G-Na-K (lacking both Na_2_HPO_4_ and KH_2_PO_4_ salts) plates supplemented with 0.5 % sucrose. In these conditions, we successfully generated Δ*kup* mutant candidates that were then tested for growth at different K^+^ regimes. Interestingly, we observed that the Δ*kup* mutant grew in limiting K^+^ concentration (M2G-K supplemented with 0.025 mM K^+^), although at a slower growth rate but with a higher plateau compared to the WT (**Fig. 5A**). As expected from the Tn-seq data, we found that Δ*kup* failed to grow when exposed to abundant and excessive K^+^ concentrations (**Fig. 5A**). We also confirmed that *kup* is essential in hypertonic conditions since the mutant did not grow in limiting K^+^ conditions when 3% sucrose were added. (**Fig. 5A**). Furthermore, we also found that overexpression of *kup* in WT is detrimental for growth in limiting K^+^ conditions, at least when the Kdp transporter is functional (**Fig. 5B**). Altogether, these data indicate that Kup allows potassium uptake in abundant and excessive K^+^ conditions but can be detrimental for unknown reasons at limiting K^+^ concentrations.

**Figure 5.**
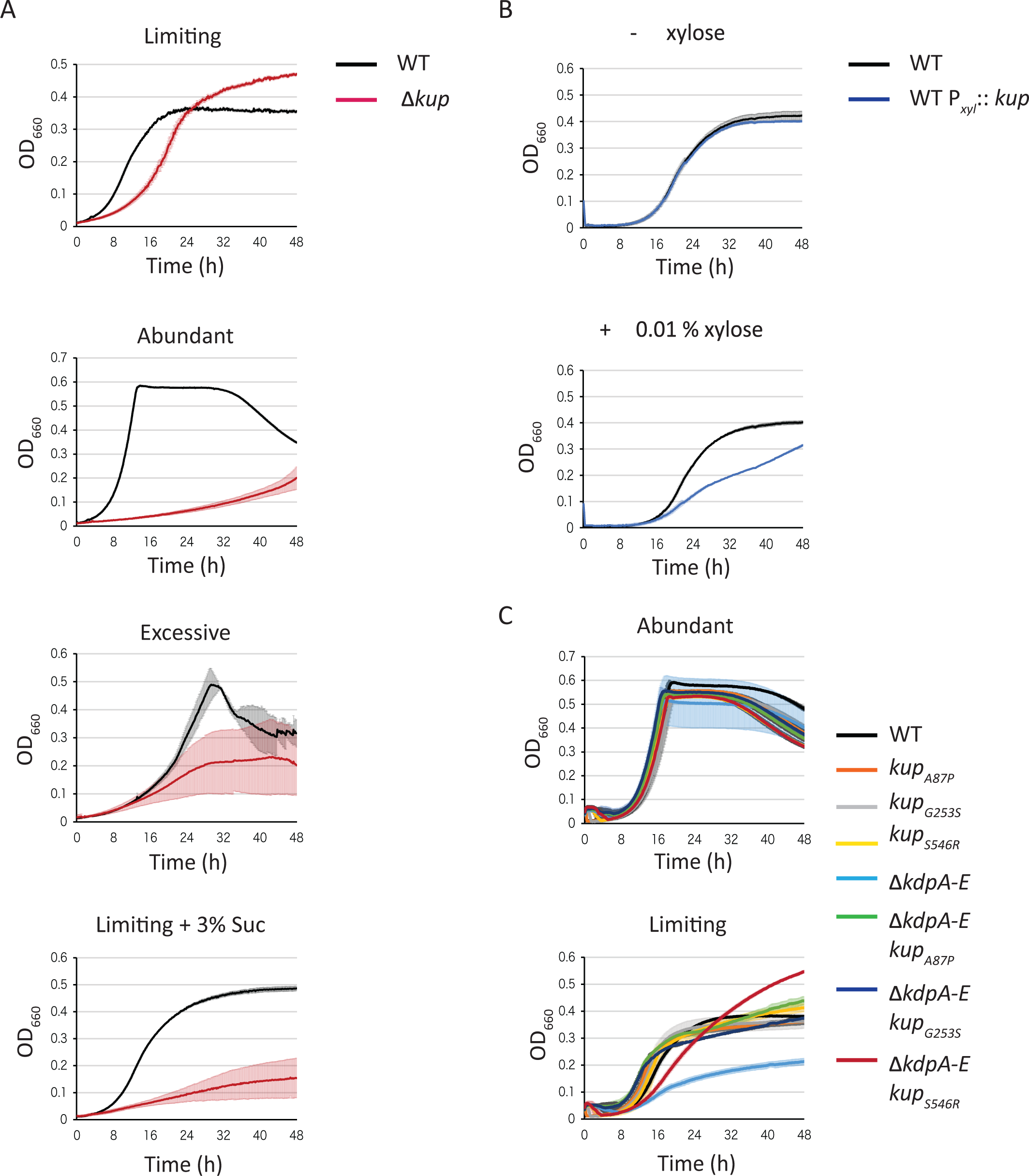
Impact of deletion and overexpression of the *kup* gene. (A) Growth of WT and Δ*kup* mutant in minimal media M2G-K supplemented with limiting (0.025 mM), abundant (0.5 mM), excessive (50 mM) K^+^ concentrations or limiting (0.025 mM) K^+^ concentrations and 3% sucrose. (B) Growth of WT expressing *kup* under xylose induction. C. Growth of *kup_A87P_*, *kup_G253S_* and *kup_S456R_* single and double mutants (with Δ*kdpA-E*) in abundant and limiting K^+^ conditions. Data represent average, n=3, and error bars= ±SD.

While characterizing the growth of Δ*kdp* cells on M2G-K plates (**Fig. 4A**), we inadvertently selected growth suppressors. These suppressors improved the growth of Δ*kdp* in limiting K^+^ conditions (**Fig. S2D**). Whole genome sequencing analyses of several of these suppressors allowed the identification of three mutations in *kup*: *kup_A87P_*, *kup_G253S_* and *kup_S456R_*. Using homologous recombination, we constructed these mutations in WT and Δ*kdp* backgrounds. First, when cells were grown in minimal media supplemented with abundant K^+^ concentrations, these mutations did not have any significant fitness cost whatever *kdp* was functional or not (**Fig. 5C**). Second, the single point mutants in an otherwise WT background grew similarly to the WT strain at limiting K^+^ concentrations (**Fig. 5C**). Third, each *kup* mutation suppressed the defect in of Δ*kdp* when grown in limiting K^+^ condition (**Fig. 5C**).

To get molecular insight into the mutation-mediated improved K^+^ transport by Kup, molecular dynamics (MD) simulations were carried out on the modeled WT Kup protein and each of the 3 mutants (A87P, G253S, and S456R). To highlight structural changes that could promote K^+^ binding/transport, 1 µs MD simulations were done in an environment mimicking the lipid composition of *Caulobacter* inner membrane (41, 42) (**Fig. S3A**) and Kup coordinates were aligned to one protomer of *Bacillus subtilis* KimA (PDB entry: 6S3K), the only available (Cryo-EM) structure of a KUP family protein (43) (**Fig. 6A**). The 8 residues in KimA described to be involved in the coordination and transport of two K^+^ cations are conserved in Kup with the following correspondences (KimA/Kup): D36/D54, Y43/Y61, D117/D146, T121/T150, S125/S154, T230/T260, E233/E263 and Y377/Y410.

**Figure 6.**
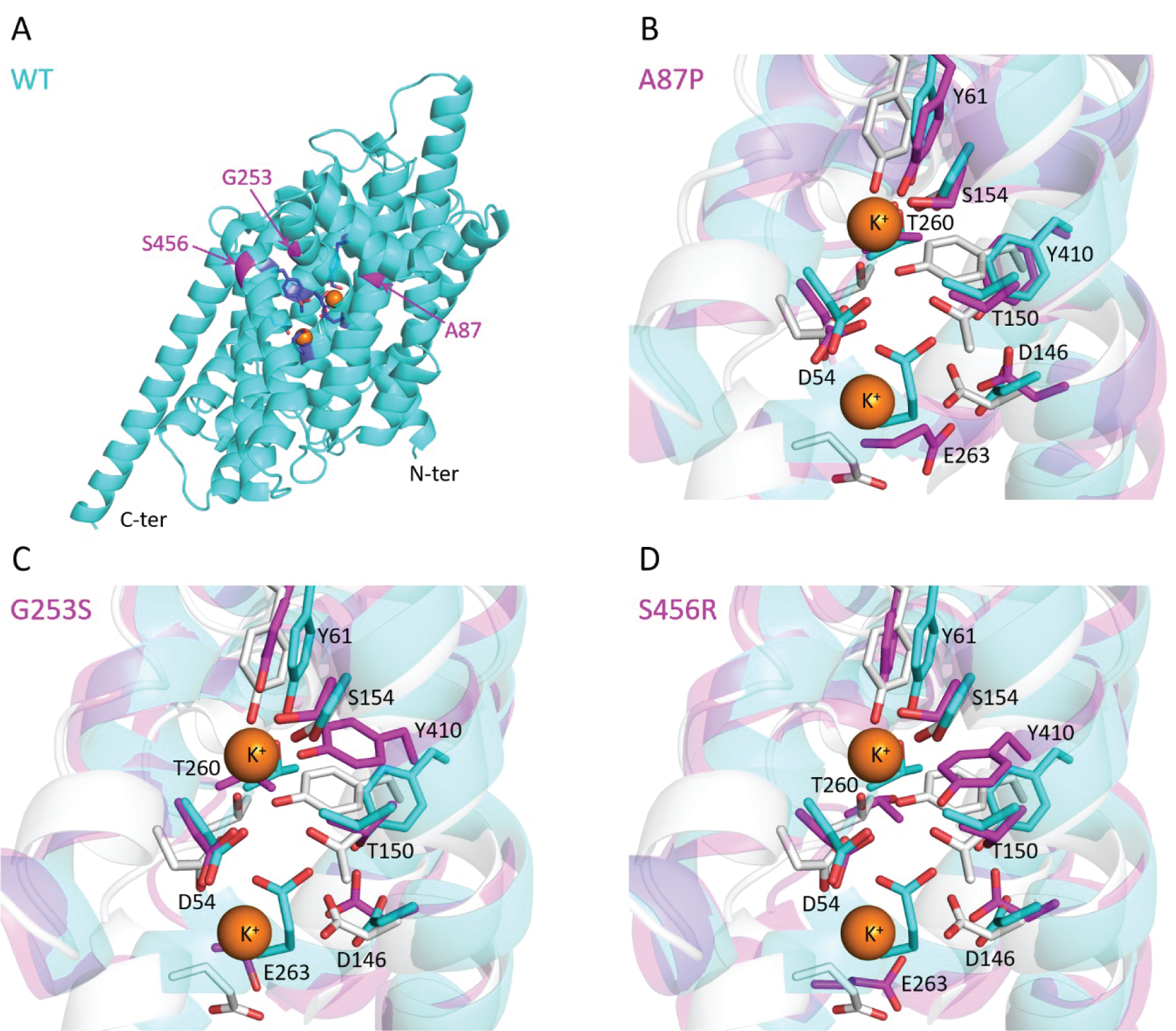
Molecular dynamics-highlighted structural changes of the putative K^+^ binding site of Kup mutants. (A) Snapshot of Kup WT after 1 µs of simulation. The protein is represented in cyan as cartoon, the considered mutation sites are colored in magenta and indicated by arrows, and the residues composing the KimA-based K^+^ binding cavity are highlighted as sticks in dark blue. (B) Snapshot of the KimA-based K^+^ binding site of Kup A87P, (C) G253S, and (D) S456R after 1 µs of simulation. On each of these panels, Kup WT (cyan) and mutated Kup (magenta) aligned to KimA (PDB entry: 6S3K; light grey) are shown as cartoon, conserved residues between Kup and KimA constitutive of the K^+^ binding pocket are represented as sticks in the corresponding color, and K^+^ cations as orange spheres. The displayed amino acid numbering only corresponds to Kup sequence.

The A87P mutation leads to only a few change, the WT conformation being mainly retained (**Fig. 6B**). However, a flipping of E263 towards the orientation described for the corresponding glutamate in KimA (E233) is observed. The same observation was made with the G253S and S456R mutations (**Fig. 6C and 6D**), suggesting that E263 reversal is structurally relevant for the increase of K^+^ transport by the Kup mutants. Furthermore, although to a lesser extent, the position of another residue is also changed in the 3 mutants. Indeed, D146 points more towards the K^+^ binding cavity, making the carboxylate group more available for coordination. Finally, G253S and S456R mutants are characterized by a reorientation of both Y61 and Y410 to a position close to that of Y43 and Y377, respectively, in KimA, with the tyrosyl moiety directed towards the putative K^+^ location. Thus, our MD simulations suggest that despite their different locations on Kup, the 3 mutations lead to subtle structural changes in the orientation of key residues described to directly participate in the binding and/or transport of K^+^.

Altogether, these data indicate that (i) Kup allows potassium uptake in abundant and excessive K^+^ conditions, and that (ii) mutations in Kup improve K^+^ transport at limiting concentrations, likely by increasing affinity for K^+^.

### K^+^- and KdpE-dependent regulons

To further characterise the response of *C. crescentus* to K^+^-depleted conditions, we performed RNA-seq on RNA samples extracted from cells grown in minimal media in K^+^ limiting or abundant conditions. In our experiment, we identified a total of 594 genes – 385 upregulated and 209 downregulated – whose expression was impacted upon K^+^ limitation with at least a 2-fold change (FC ≥ 2) and a false-discovery rate (FDR) lower than 5% (P*adj* ≤ 0.05) (**Fig. 7A**). As expected, this experiment confirmed that the expression of *kdp* genes was strongly induced in the K^+^-depleted medium. Among the other upregulated genes, we found genes coding for the phosphate starvation protein PhoH (CCNA_02727), a xylose isomerase family protein (CCNA_01701), a TonB-dependent receptor (TBDR, CCNA_01859), an AraC family transcription regulator (CCNA_01858) and hypothetical proteins (CCNA_03313 and CCNA_03970) (**Fig. 7A** and **Table S3A**). Among the top downregulated candidates, we found genes expressing a small RNA (CCNA_R0158), a hemin receptor (CCNA_02277), a putative transporter (CCNA_03022), a protein with a calcium binding EF Hand binding domain (CCNA_02274) and the chaperons GroES (CCNA_00722) and GroEL (CCNA_00721) (**Fig. 7A** and **Table S3B**).

**Figure 7.**
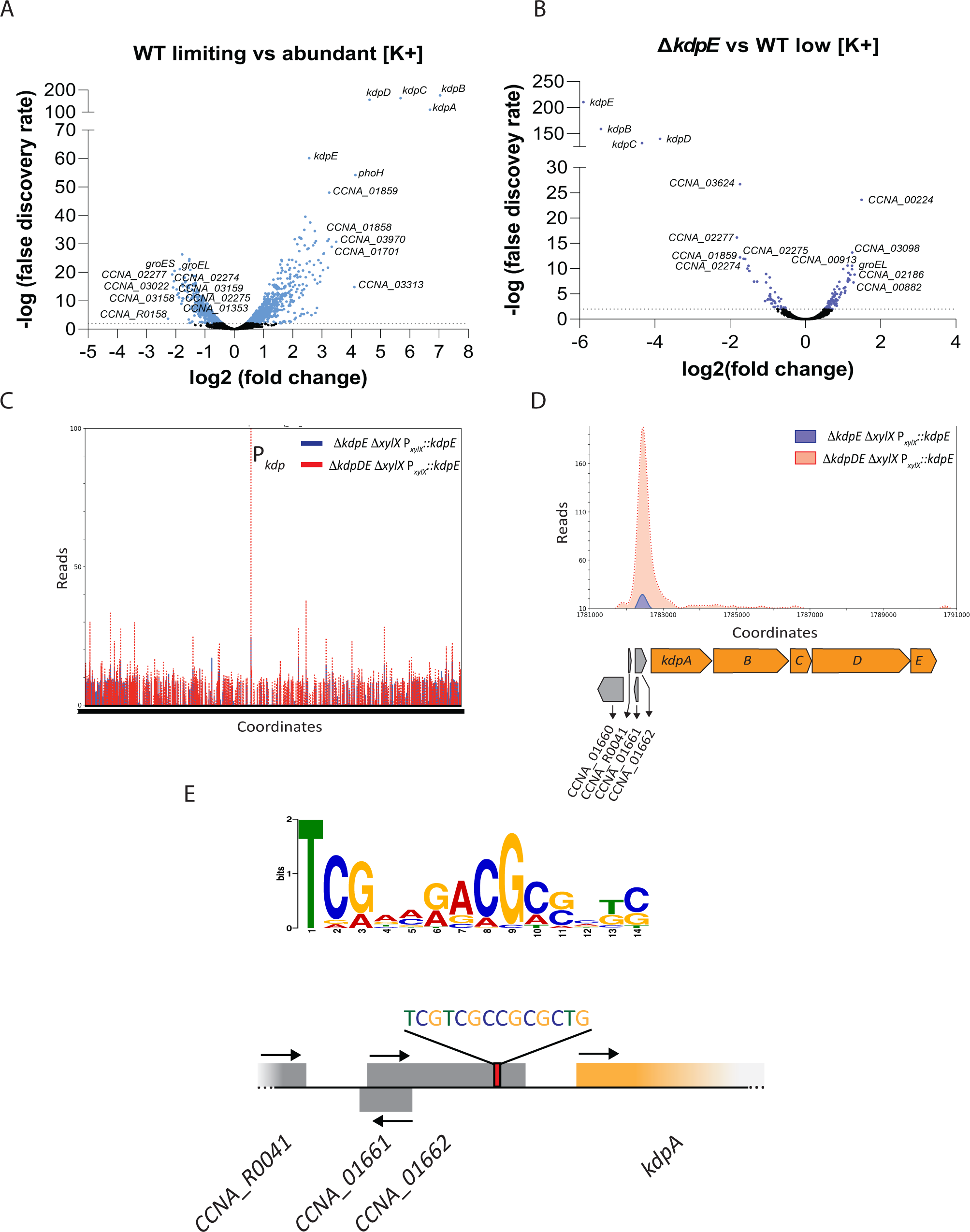
K^+^ and KdpE-dependent regulon. (A-B) Volcano plots representing the relation between the log_2_ fold change (FC) and -log False Discovery Rate (FDR) on gene expression between (A) WT in limiting *vs* abundant K^+^ condition, and (B) Δ*kdpE* vs WT in limiting K^+^ condition. Genes identified are presented as dots. Significant down- and up-regulated genes are presented as blue (A) and purple (B) dots, while genes with no significant alterations are presented as black dots. Top 10 down and upregulated genes are highlighted for each analysis. (C) Genome-wide occupancy of KdpE on the chromosome of *C. crescentus* determined by ChIP-seq on Δ*kdpE* P*_xylX_*::*kdpE* Δ*xylX* and Δ*kdpDE* P*_xylX_*::*kdpE* Δ*xylX*. The x-axis represents the coordinates on the genome (Mb), the y-axis shows the normalized ChIP-Seq read abundance in reads. The promoter region of *kdp* is highlighted. (D) Occupancy of KdpE on the *kdp* promoter Δ*kdpE* P*_xylX_*::*kdpE* Δ*xylX* and Δ*kdpDE* P*_xylX_*::*kdpE* Δ*xylX*. (E) WebLogo of predicted KdpE consensus sequence obtained with MEME and identification (with a *p-value* 7.71 x 10^-5^) of the DNA binding sequence upstream of *kdpA.* Arrows indicate gene orientation for expression for each gene.

As our results suggest that the Kdp system is important for growth upon K^+^ depletion, we also performed RNA-seq experiments with a Δ*kdpE* mutant grown in M2G-K supplemented with limiting K^+^ concentrations. Then, we compared the RNA enrichment in Δ*kdpE vs* WT to determine which genes are affected by KdpE in a K^+^-depleted condition using FC ≥ 1.5 and *P*_adj_ ≤ 0.05 as parameters. Compared to WT, we found 83 upregulated and 50 downregulated genes in Δ*kdpE* grown in the K^+^-depleted media (**Fig. 7B** and **Tables S4AB**). Among the 50 downregulated genes in Δ*kdpE* (**Table S4B**), 72% (36/50) were also found in the regulon of the WT starved for K^+^, 19 upregulated (including *kdp* genes; **Table S3A**) and 17 downregulated (**Table S3B**). This suggests that KdpE is strictly required to activate the expression of the 19 genes in a K^+^-depleted environment. On the contrary, the 17 other genes that are repressed in WT cells upon K^+^ depletion are even further downregulated in the absence of KdpE, suggesting that KdpE limits the repression of these genes. The expression of the last 14 genes was insensitive to K^+^ concentration while downregulated in Δ*kdpE* (**Tables S4AB**), suggesting that they are positively regulated by KdpE but independently of the K^+^ concentration. Among the 83 upregulated genes in Δ*kdpE* (**Table S4A**), only ∼36% (30/83) were also found in the regulon of the WT starved for K^+^, 12 downregulated (including *groEL* and *groES* genes; **Table S3B**) and 18 upregulated (**Table S3A**). This indicates KdpE is required to repress the first 12 genes upon K^+^ depletion while it limits the activation of the other 18 genes in a K^+^-depleted environment. The remaining 53 upregulated genes in Δ*kdpE* (**Tables S4AB**) were insensitive to K^+^ availability, suggesting that they are likely repressed by KdpE but in a K^+^-independent way.

Surprisingly, the vast majority (528 genes; 88.9 %) of the 594 genes whose expression changed in the WT grown in K^+^ limiting conditions seems to be KdpE-independent, suggesting that other regulators, yet to be discovered, are activated upon K^+^ depletion. An alternative explanation might be that Δ*kdpE* cells exposed to limiting K^+^ concentration suffer from a general stress response, which ultimately leads to global transcriptional changes. This could also explain why half of the 133 genes (67/133) found in the Δ*kdpE* regulon were not identified in the WT regulon.

In order to unveil genes whose expression is directly regulated by KdpE, we identified the DNA regions bound by KdpE using ChIP-seq. For that, we first constructed strains in which *kdpE* was expressed under the xylose inducible promoter (P*_xylX_*::*kdpE*) in a Δ*kdpE* Δ*xylX* background. Inactivating *xylX* avoids xylose consumption by *C. crescentus* as a carbon source thereby allowing constant expression of *kdpE* upon induction. Expression of *kdpE* from P*_xylX_* efficiently complemented the growth of both strains in the presence of xylose since Δ*kdpE* Δ*xylX* P*_xylX_*::*kdpE* cells grew similarly to the WT or Δ*xylX* cells whereas Δ*kdpDE* Δ*xylX* P*_xylX_*::*kdpE* cells grew at a higher plateau (**Fig. S4A**) like Δ*kdpD* cells (see below). Moreover, using polyclonal anti-KdpE antibodies, KdpE was easily detected by western blot in both strains although KdpE protein levels were lower in Δ*kdpDE* Δ*xylX* than in Δ*kdpE* Δ*xylX* (**Fig. S4B**), suggesting that KdpD has a global positive impact on KdpE levels. It is noteworthy that KdpE was undetectable in WT strain, at least at high K^+^ concentrations, indicating that P*_xylX_*::*kdpE* leads to KdpE overexpression upon xylose induction.

A total of 110 and 146 DNA binding sites were respectively detected in the Δ*kdpE* Δ*xylX* P*_xylX_*::*kdpE* and Δ*kdpDE* Δ*xylX* P*_xylX_*::*kdpE* strains grown in K^+^ limiting conditions (**Fig. 7C**; **Tables S5** and **S6**). About a third of the DNA regions bound by KdpE identified in both strains – 40 out of 110 (36.4%) found in Δ*kdpE* Δ*xylX* P*_xylX_*::*kdpE* and 46 out of 146 (31.5%) found Δ*kdpDE* Δ*xylX* P*_xylX_*::*kdpE* – are located in the close vicinity of genes whose expression showed a FC ≥ 1.5 in RNA-seq experiments (highlighted in green in **Tables S5** and **S6**), supporting that these regions are indeed KdpE-binding sites.

The highest numbers of reads were found in the *kdp* promoter region, not only in Δ*kdpE* Δ*xylX* P*_xylX_*::*kdpE* cells but more surprisingly also in Δ*kdpDE* Δ*xylX* P*_xylX_*::*kdpE* cells. Even more surprising is the fact that the peak in the Δ*kdpDE* background was higher than in the Δ*kdpE* one (**Fig. 7C**). In fact, this is true for most if not all regulated genes, but the difference was particularly important for the *kdp* promoter (**Fig. 7D**), suggesting that KdpE could be hyperphosphorylated in the absence of KdpD and therefore that KdpD mainly acts as a negative regulator of KdpE∼P. Moreover, it indicates that, although P*_kdp_* is the main target of KdpE, multiple other genes are potential direct targets of KdpE. Analysis of the top 50 sequences identified in the ChIP-seq with the Δ*kdpE* Δ*xylX* P*_xylX_*::*kdpE* strain using MEME allowed the identification of the conserved DNA motif TCGAMRACGCSMKC likely bound by KdpE (**Fig. 6E**). For the peak in the vicinity of the *kdpABCDE* operon (top hit of the analysis), the KdpE motif was located 113 bp upstream of the *kdpA* start codon, within the CCNA_01662 gene (**Fig. 7E**).

### *C. crescentus* KdpD regulates KdpE both positively and negatively

To further assess the role of the *C. crescentus* Kdp system in K^+^-limiting conditions, we first constructed KO mutants of the transport complex (Δ*kdpABC*) and the TCS (Δ*kdpDE*) genes. In comparison to the WT, both Δ*kdpABC* and Δ*kdpDE* mutants had a growth delay at limiting but not at abundant K^+^ concentration (**Fig. 8A** and **Fig. S5A**). Moreover, both mutants grew similarly, suggesting that the *kdpABC* is not expressed in the absence of the TCS, as described in other bacteria.

**Figure 8.**
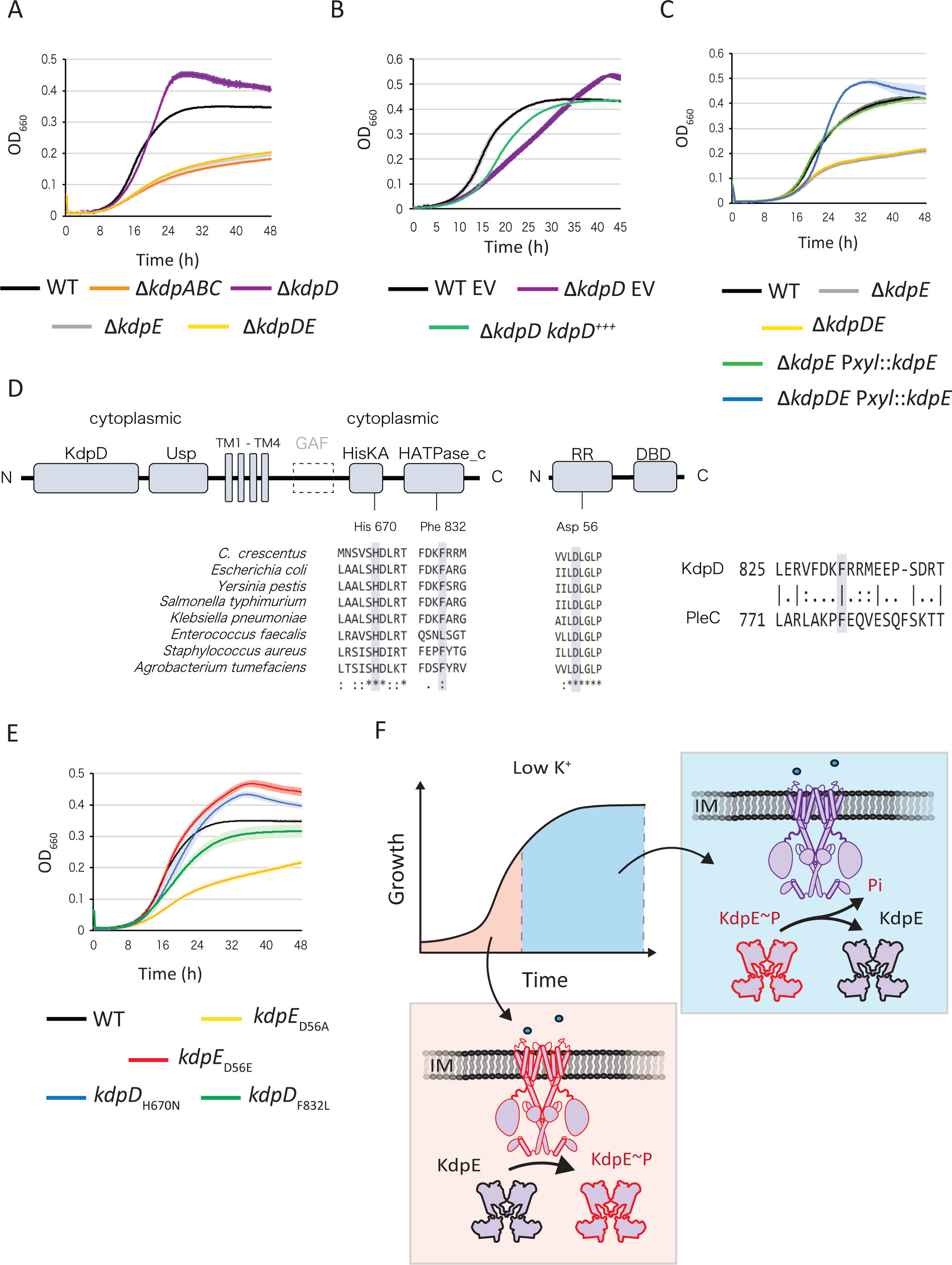
KdpD downregulates KdpE to control growth in low K^+^. (A) Growth of WT, Δ*kdpABC,* Δ*kdpD*, Δ*kdpE* and Δ*kdpDE*. (B) Growth of WT and Δ*kdpD* carrying either pMR10 empty vectors (EV) or pMR10 P*_kdp_::kdpD*. (C) Growth of WT, Δ*kdpE*, Δ*kdpDE* expressing *kdpE* from the *xylX* locus (P*_xylX_::kdpE*). (D) Domain architectures and relevant conserved residues in *C. crescentus* KdpD and KdpE from different bacteria, and between KdpD and PLeC from C. crescentus. Protein domains were predicted by searching the Pfam database (cut-off E value = 1.0). Accession numbers are as follows: KdpD (PF02702), Usp (PF00582), Dhp (PF00512), CA (PF02518), RD (PF00072) and DBD (PF00486). Multiple sequences alignments were performed with Clustal Omega. Shown KdpD protein sequences were as follows: *C. crescentus*, Uniprot: A0A0H3C8H5; *E. coli*, UniProt: P21865; *Y. pestis*, UniProt: Q7CJR5; *S. typhimurium*, UniProt: Q8ZQW4; *K. pneumoniae*, UniProt: B5XZF1; *E. faecalis*, UniProt: Q838F4; *S. aureus*, UniProt: A0A0H2XIE2; and *A. tumefaciens*, Uniprot: A0A2L2LHM9. Shown KdpE proteins sequences were as follows: *C. crescentus*, Uniprot: A0A0H3C8M3; *E. coli*, UniProt: P21866; *Y. pestis*, UniProt: Q0WDK2; *S. typhimurium*, UniProt: Q8ZQW5; *K. pneumoniae*, UniProt: A0A0W8ASI2; *E. faecalis*, UniProt:Q8KU70; and *S. aureus*, UniProt: Q2FWH6; and *A. tumefaciens*, Uniprot: A0A1B9TGM5. PleC protein sequence: P37894 (E) Growth of *kdpD* and *kdpE* catalytic point mutants. Growth was done in minimal media M2G-K supplemented with limiting (0.025 mM) K^+^ concentrations (A-C and E), media was additionally supplemented tetracycline for plasmid selection (B) or with 0.01 % xylose for the induction of *kdpE* expression (C). Data represent average, n=3, and error bars= ±SD.

To study the HK *kdpD* and RR *kdpE* genes independently of each other, we constructed single KO mutants, Δ*kdpD* and Δ*kdpE*. Deletion of *kdpE* resulted in a similar growth delay than the ones displayed by Δ*kdpABC*, Δ*kdpDE* and Δ*kdpABCDE* (**Fig. 8A** and **Fig. S5B**). On the contrary, Δ*kdpD* behave differently since compared to WT, it had a slight but reproducible growth delay in exponential phase while it later reached a higher plateau in stationary phase (**Fig. 8A**). This peculiar growth behaviour was complemented by expressing *kdpD* from the native *kdp* promoter (P*_kdp_*::*kdpD*) on a replicative and low-copy number plasmid in a Δ*kdpD* background (**Fig. 8B**). Complementation of the Δ*kdpE* mutant was achieved by expressing *kdpE* from the xylose inducible promoter (P*_xylX_*::*kdpE*) either in a Δ*kdpE* or a Δ*kdpDE* background. While the growth defect of Δ*kdpE* was fully complemented, even in the absence of the xylose inducer, expressing back *kdpE* in Δ*kdpDE* cells led to the same growth behaviour than the single Δ*kdpD* mutant, that is a slight growth delay in exponential phase and a higher plateau in stationary phase (**Fig. 8C** and **Fig. S5C**). In addition, the fact that Δ*kdpE* is epistatic on Δ*kdpD*, since Δ*kdpDE* grew as poor as Δ*kdpE* or Δ*kdpABC* (**Fig. 8A**), shows KdpE is entirely responsible for the Δ*kdpD* phenotypes. Together, these data support that both components of the TCS are important for the growth in K^+^-depleted environments and suggest that KdpD and KdpE have antagonistic roles for growth in such conditions. Kdp regulatory proteins KdpD and KdpE were further analysed by identifying domains and their organisation. Comparison of Pfam features highlighted that the KdpD sequence of *C. crescentus* lacks the cytoplasmic GAF domain known to be linked to the DHp domain in *E. coli* (44) (**Fig. 8D**). Also, multiple sequence alignments enabled to locate the conserved catalytic – phosphorylatable – residues in both the HK (KdpD) and the RR (KdpE), that are respectively the histidine 670 (His670) and the aspartate 56 (Asp56) (**Fig. 8D**). To better understand the KdpDE regulatory mechanism, we constructed *kdpD and kdpE* catalytic point mutants and assessed their impact on growth in abundant and limiting K^+^ conditions. In agreement with what we observed with the KO mutants, all the catalytic mutants grew as good as the WT in abundant K^+^ conditions (**Fig. S5D**). At limiting K^+^ concentrations however, the phospho-ablative mutant of KdpE (*kdpE_D56A_*) had a strong growth delay, similar to the one displayed by Δ*kdpE* (**Fig. 8E** and **Fig. S5E**). In contrast, the phospho-mimetic mutant (*kdpE_D56E_*) grew similarly to the WT up to the late exponential phase, to finally reach a higher final OD comparable to the one observed with Δ*kdpD* grown in same conditions (**Fig. 8E** and **Fig. S5E**). As expected, a KdpD catalytically dead mutant (*kdpD_H670N_*) phenocopied Δ*kdpD* in limited K^+^ (**Fig. 8E** and **Fig. S5E**). Since *kdpE_D56A_* cells had a much stronger growth delay than Δ*kdpD* or *kdpD_H670N_*, it indicates that KdpE is still phosphorylated despite the absence of KdpD kinase activity. Thus, KdpE is very likely phosphorylated by another – non-cognate – HK. On the other hand, the fact that *kdpE_D56E_* cells grew slightly but reproducibly better than Δ*kdpD* and *kdpD_H670N_* suggests that KdpD also phosphorylates KdpE, at least during the early exponential phase of growth. Thus, KdpD might (i) phosphorylate KdpE in early exponential phase, with at least another HK, and (ii) mainly act as a phosphatase on KdpE∼P in late exponential phase, likely to downregulate the expression of *kdp* and restrict growth in K^+^-limiting conditions (**Fig. 8F**).

To test this hypothesis, we constructed a KdpD kinase-deficient but phosphatase active mutant (K^-^/P^+^) by replacing a conserved phenylalanine residue by a leucine residue (*kdpD_F832L_*) in a way reminiscent to what was described for another HK encoding gene, *pleC_F778L_* (45) (**Fig. 8D**). Interestingly, we found that unlike the *kdpD_H670N_*, *kdpD_F832L_* cells failed to reach a higher plateau in stationary phase (**Fig. 8E** and **Fig. S5E**), strongly supporting our hypothesis that KdpD primarily works as a phosphatase in late exponential phase. Moreover, the K^-^/P^+^ mutant *kdpD_F832L_* grew better than *kdpE_D56A_* and Δ*kdpE* mutants in low K^+^ conditions (**Fig. 8E** and **Fig. S5E**), supporting that there is indeed at least another HK that phosphorylates KdpE. Finally, the fact that the *kdpD_F832L_* mutant had a growth delay compared to WT suggests that KdpD also acts as a kinase for KdpE in early exponential phase (**Fig. 8E** and **Fig. S5E**). However, we cannot rule out the possibility that the phosphatase activity of KdpD_F832L_ is impacted by the mutation. For instance, it might be stronger than the WT thereby interfering with the phosphorylation levels of KdpE in early exponential phase. Nevertheless, our results altogether indicate that (i) KdpD works both as a kinase and a phosphatase to control KdpE phosphorylation (KdpE∼P) levels depending on the growth phase and K^+^ availability and that (ii) KdpE can be phosphorylated by (an)other non-cognate HK(s) for regulating K^+^ transport in K^+^-depleted environments.

### KdpD-dependent activation of the *kdp* promoter in low K^+^ conditions

In order to study the Kdp system at the transcriptional level, a plasmid carrying a P*_kdp_*::*lacZ* reporter fusion was introduced in the different mutant strains, cells were grown in limiting and abundant K^+^ concentrations and β-galactosidase activity was measured. Consistent with the idea that the Kdp system is a high-affinity potassium transporter, P*_kdp_* basal activity in WT cells was barely detectable (∼300 Miller Units) at abundant K^+^ concentration while it was strongly induced in WT (∼9.000 Miller Units) or in Δ*kdpABC* cells (∼10.500 Miller Units) upon K^+^ depletion (**Fig. 9A**). In contrast, P*_kdp_* activity was completely annihilated in Δ*kdpE* and *kdpE_D56A_* cells (**Fig. 9A**), indicating that phosphorylated KdpE (KdpE∼P) is strictly required for *kdp* expression both in limiting and abundant K^+^ conditions. In support of that, the basal activity of P*_kdp_* at abundant K^+^ concentration was 10 times higher (∼3.000 Miller Units) in *kdpE_D56E_* cells than in WT cells (**Fig. 9A**). Interestingly, P*_kdp_* was still active at low K^+^ concentration in a Δ*kdpD, kdpD_H670N_,* and *kdpD_F832L_* background (∼4.600, ∼4.200, ∼4.700 Miller Units, respectively), but not to the same extent than in WT cells (∼9.000 Miller Units) (**Fig. 9A**). In addition, the basal P*_kdp_* activity at abundant K^+^ concentration was completely abolished in the strains expressing the K^-^/P^+^ *kdpD_F832L_* variant, suggesting that in the WT, the KdpD phosphatase activity is counterbalanced by its kinase activity on KdpE in K^+^-replenished conditions. In support of that, compared to WT cells grown in abundant K^+^ condition, the basal P*_kdp_* activity was higher when both the hydrolase and kinase activities were inactivated (K^-^/P^-^), that is in Δ*kdpD* cells and to a lesser extent in *kdpD_F832L_* cells.

**Figure 9.**
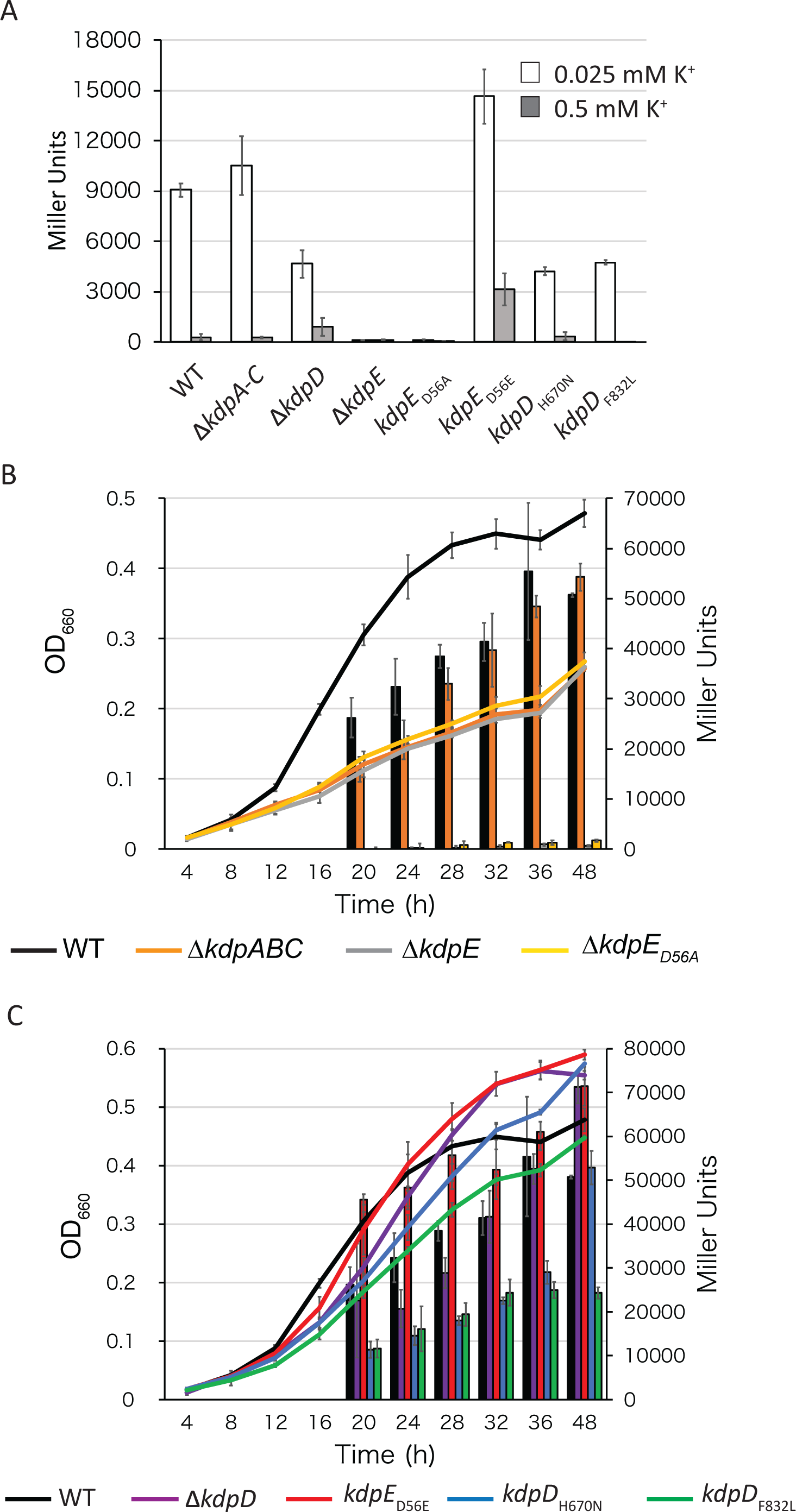
Regulation of *kdp* expression. (A) β-Galactosidase activities of WT and *kdp* mutants carrying a transcriptional P*_kdp_*::*lacZ* fusion grown in M2G-K supplemented with limiting (0.025 mM) or abundant (0.5 mM) K^+^ concentrations. β-Galactosidase activities of WT and *kdp* mutants carrying a transcriptional P*_kdp_*::*lacZ* fusion along the growth in M2G-K supplemented with limiting (0.025 mM).. (B-C) β-Galactosidase activities of WT and *kdp* mutants carrying a transcriptional P*_kdp_*::*lacZ* over time of growth in M2G-K supplemented with limiting (0.025 mM) K^+^ concentration. Statistical analyses were carried out via single factor ANOVA. Data represent average, n=3, and error bars= ±SD.

Then, we followed the expression of the P*_kdp_*::*lacZ* fusion in WT and *kdp* mutant cells along the growth at limiting K^+^ concentration. Despite Δ*kdpABC*, Δ*kdpE* and *kdpE_D56A_* all poorly grew in limiting K^+^ conditions, P*_kdp_* was active in Δ*kdpABC* cells to a level comparable to the WT one while P*_kdp_* activity was almost undetectable in both *kdpE* mutants (**Fig. 9B**). These data suggest that (i) the Kdp transporter (KdpABC) is not required for *kdp* expression while (ii) the response regulator is, in contrast, indispensable. In support of this, we found the phospho-mimetic mutant of *kdpE* (*kdpE_D56E_*) displayed a relatively constant but high activity of P*_kdp_* alongside the growth curve (**Fig. 9B**).

The *kdpD* mutants had a different behaviour. First, compared to WT, the activity of P*_kdp_* in Δ*kdpD* cells was lower during early exponential phase but higher in stationary phase (**Fig. 9C**). Similarly, *kdp* expression was significantly lower both in *kdpD_H670N_* (K^-^/P^-^) and in *kdpD_F832L_* (K^-^/P^+^) during early exponential phase. However, P*_kdp_* activity strongly increased when reaching the late exponential only in *kdpD_F832L_* cells. Altogether, these results support our hypothesis that (i) KdpD plays dual antagonistic activities on KdpE, as a kinase in early exponential phase and a phosphatase in late exponential phase of growth, and that (ii) KdpE can be phosphorylated by another non-cognate HK all along the growth.

## Discussion

As the most abundant monovalent cation used by living cells to drive many cellular processes, regulating potassium (K^+^) homeostasis is critical for survival. The oligotrophic α-proteobacterium *Caulobacter crescentus* is cyclically subjected to nutrients deprivation in its natural environments. In contrast to carbon, nitrogen or phosphate, the response of *C. crescentus* to K^+^ depletion has never been characterised. Here, we first determined the range of K^+^ concentrations at which *C. crescentus* supports growth. In addition, we identified all the genes predicted to code for proteins involved in transport, sensing and regulation of K^+^ homeostasis and tested their growth behaviour at low (limiting), optimal (abundant) and high (excessive) K^+^ concentrations. Then, we defined the set of genes that (i) become essential and/or (ii) are induced upon K^+^-environmental depletion. This allowed us to identify the low affinity and the high affinity K^+^ transporters, respectively Kup and KdpABC, as critical components that maintain K^+^ homeostasis in *C. crescentus* (**Fig. 10**). Finally, we deeply characterized the role played by the two-component (TCS) regulatory system KdpDE in the response to K^+^ depletion.

**Figure 10.**
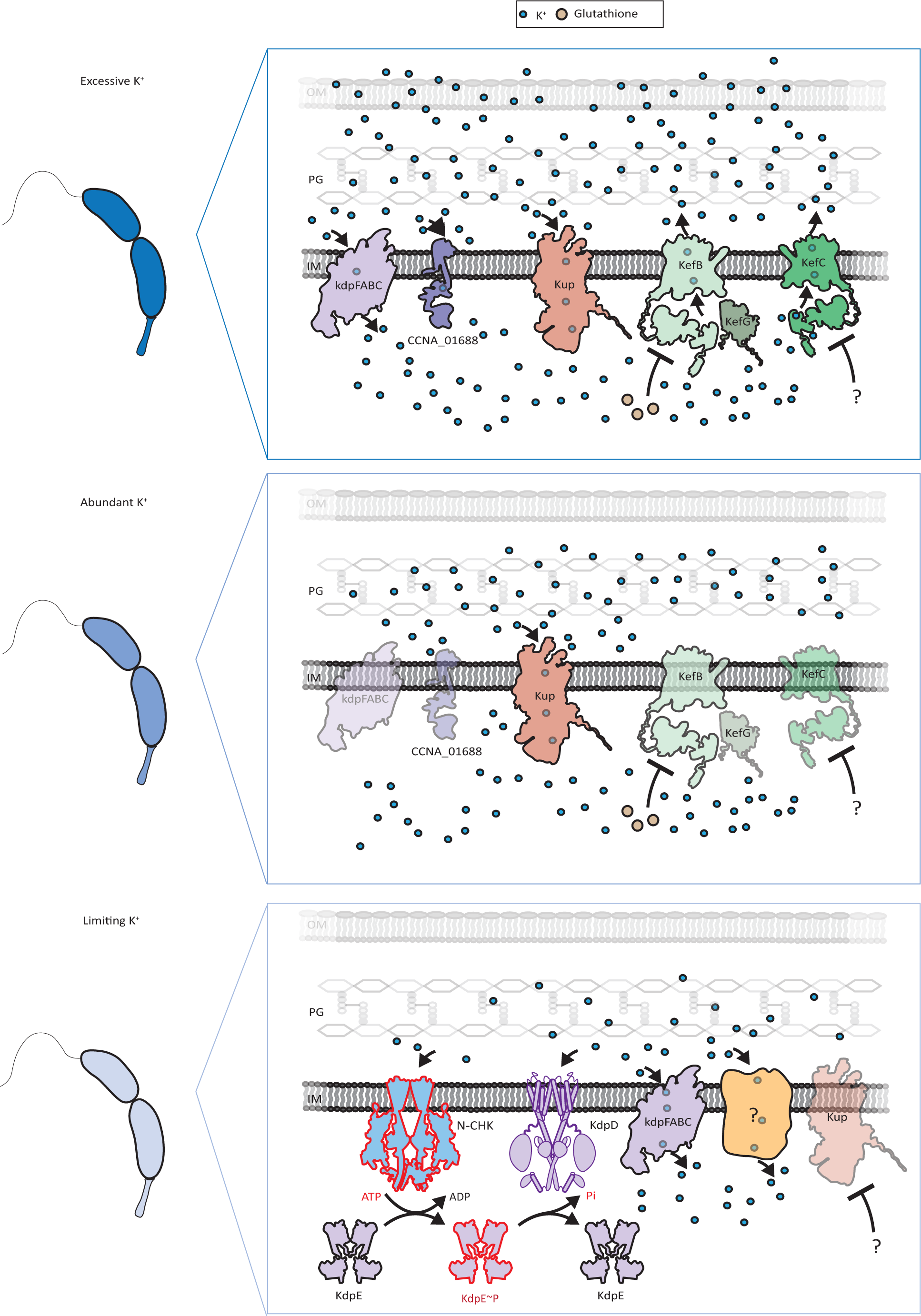
Model of K^+^ transport in *C. crescentus.* Our data indicate that in excess of K^+^, the uptake transporters Kup, CCNA_01688 and Kdp, as well as the efflux pump KefC are important actors for K^+^ homeostasis, while KefB plays a minor role probably under glutathione regulation (see discussion in text). At a K^+^ abundant concentration, Kup is the main transporter. In contrast, in limiting conditions, our data suggest that Kdp and (an)other transporter(s) (Kup or unknown protein in yellow) are the main transporters. KdpD acts as a bifunctional kinase/phosphatase and KdpE is phosphorylated by KdpD and a non-cognate histidine kinase (N-CHK) to control *kdp* expression, K^+^ uptake and growth. In contrast, in such conditions, *kup* expression can be detrimental, and is potentially downregulated through a yet unknown mechanism.

By analysing the Tn-seq data, we found that the COG categories – biological functions – for which the highest number of genes became essential or high fitness cost (HFC) upon K^+^ limitation (M2G *vs* M2G-K) were (i) C “energy production and conversion” (∼1% increase of essential and ∼6 % increase of HFC) and (ii) J “Translation, ribosomal structure and biogenesis” (∼2% increase of essential and ∼3 % increase of HFC). In fact, it is not so surprising to highlight these two categories as sensitive to K^+^ availability. Indeed, ATP concentration and respiration state are two well-known parameters that drive K^+^ transport across the membrane. For instance, the electron transport chain determines the membrane potential, which in turn will influence the distribution of other ions like K^+^ that accumulate on the cytoplasmic face of the membrane. This might therefore explain why Tn insertions in genes coding for subunits of the respiratory chain or the ATP synthase are counter-selected when K^+^ become limiting. On top of this, active transport of K^+^ across the membrane by the high-affinity transporter consumes a lot of energy provided by ATP, so that decreasing ATP concentration likely has a negative impact on K^+^ import. On the other hand, K^+^ cations are critical for maintaining the structure and sustaining the function of ribosomes. Not only K^+^ is required to stabilize the transfer RNAs (tRNAs), the ribosomal RNAs (rRNAs) and the ribosomal proteins within the ribosomes, but also to increase proofreading by accommodating the right aminoacyl-tRNA in the ribosomal A-site (46, 47). Additionally, many GTPases involved in translation display a K^+^-dependent activation of GTP hydrolysis (1). Thus, the impact K^+^ has on translation could explain the lower Tn insertion density observed in rRNAs, tRNAs and genes coding for ribosomal proteins in cells grown in limiting K^+^ conditions.

Several physiological processes have been described to be K^+^-dependent. Besides being required by numerous enzymes related to basic metabolic functions (1, 48), such as for example the pyruvate kinase (2), K^+^ modulates the activity of enzymes involved in bacterial cell cycle. For instance, in *E. coli*, K^+^ has been shown to stimulate the polymerisation of the cell division protein FtsZ (49), the activity of the DNA polymerase (50). Moreover, the DNA supercoiling activity of the DNA gyrase was described to be triggered by K^+^ in both *B. subtilis* (51) and *Micrococcus luteus* (52). Recently, it was reported that glutathione controls cell division via negative regulation of the potassium (K^+^) efflux pump KefB (27). Indeed, cell filamentation and decreased intracellular K^+^ levels were observed in mutants unable to synthesize glutathione (Δ*gshB*), suggesting that K^+^ homeostasis is crucial for bacterial cell cycle regulation. Interestingly, mutations that suppressed Δ*gshB* phenotypes were found in *kefB* and *kefG*, but not in *kefC*. This indicated that, glutathione regulation in *C. crescentus* occurs mainly through *kefB*. Therefore, it is likely that the glutathione-dependent negative regulation on KefB still occurred in Δ*kefC* cells, which would explain why the single *kefC* knock-out mutant failed to grow when K^+^ was in excess whereas the single Δ*kefC* grew. K^+^ was also described as critical for the chaperonin activity of GroEL/GroES (46) or DnaK (53). Interestingly, we observed that *groEL* and *groES* became essential upon K^+^ depletion, thereby suggesting that some K^+^-dependent mechanisms are conserved in bacteria.

KdpDE is the major, if not the only bacterial system described in the literature to respond to limiting K^+^ availability. Although KdpE was shown in *E. coli* to regulate expression of the anti-sigma factor Rsd and the ribosome modulation factor Rmf at low K^+^ concentrations, thereby allowing KdpE to globally control transcription and translation in limiting K^+^, other regulatory systems independent of KdpDE have not been yet described in bacteria. The K^+^-dependent regulon of *C. crescentus* allowed us to identify almost 600 genes whose expression changes upon K^+^ limitation. Surprisingly, only a small proportion of this regulon depends directly or indirectly on the response regulator KdpE, suggesting that there are unknown transcriptional regulators sensitive to K^+^ availability.

The analysis of the *C. crescentus* genome allowed us to identify several genes and systems predicted to transport K^+^. By systematically characterizing the growth profile of the corresponding mutants in different K^+^ regimes, we found that the high affinity Kdp system primarily serves to import K^+^ at limiting K^+^ concentrations, like in many other bacterial species. Nonetheless, Kdp is unlikely the sole K^+^ transport system involved in this context. Despite its growth defect, the Δ*kdpABC* mutant was still able to grow at low K^+^ concentration. Since complete depletion of K^+^ led to a growth arrest in *C. crescentus*, this implies that K^+^ is imported by additional system(s), at least in limiting K^+^ conditions. Interestingly, we found that the low affinity Kup transporter became dispensable in such limiting K^+^ conditions. However, mutations in Kup have been identified to improve the growth the Δ*kdp* mutant on M2G-K plates, strongly suggesting that Kup participates to K^+^ import when it is poorly available, for instance when Kdp activity is impaired. To test whether Kup is at work at low K^+^ conditions, it might be interesting to try to combine Δ*kdp* and Δ*kup* mutations together. Nonetheless, K^+^ could be transported in such limiting conditions by other unknown transporters. Several putative transporters were identified as essential and high fitness cost in our Tn-seq analysis (i.e CCNA_01159 and CCNA_00494) and up-regulated in the RNA-seq analysis (i.e CCNA_00339, CCNA_01852, CCNA_01587, CCNA_01588 CCNA_02570, CCNA_02571, …). It is possible that one or several of them assist the activity of KdpABC in limiting K^+^.

As in *E. coli*, the expression of *kdp* operon in *C. crescentus* is activated upon K^+^ starvation by the KdpDE TCS (54). However, we found surprising differences since in the absence of the histidine kinase (HK) KdpD, (i) *kdp* is still expressed, although to a lesser extent than in WT cells and (ii) the *kdp* promoter (P*_kdp_*) is more active in stationary phase. Our genetic dissection of the catalytic activity of KdpDE with point mutants in *kdpE* (phospho-mimetic *kdpE_D56E_* and phospho-ablative *kdpE_D56A_*) and KdpD (K^-^/P^-^ *kdpD_H670N_* and K^-^/P^+^ *kdpD_F832L_*) strongly suggest that (i) KdpE can be phosphorylated by another – non-cognate – HK, at least in the absence of KdpD and that (ii) KdpD switches mainly to a phosphatase in late exponential phase of growth.

In *E. coli*, deletion of the *kdpD* results in a prominent growth arrest, of more than 20 hours, before restarting growth thanks to KdpE phosphorylation by PhoR. However, it never surpassed the WT plateau as seen with *C. crescentus* Δ*kdpD*, *kdpD_H670N_* and *kdpE_D56E_* mutants. Thus, the higher plateau is likely due to the loss of KdpD phosphatase activity. The phosphatase activity of KdpD has be shown in *E. coli* to be stimulated by intracellular K^+^ intracellular levels (21). Four amino acids located in the second periplasmic loop were identified as an extracellular K^+^-specific recognition site. Hence, substitution of those four amino acids by alanine resulted in a strain unable to sense extracellular K^+^. In contrast, the K^+^-recognition site for intracellular sensing remains to be identified. However, although specific residues could not be highlighted, the C-terminal cytoplasmic domain of KdpD was described to display a K^+^-dependent phosphatase activity (20, 55). Although this has yet to be experimentally investigated, authors suggested that the cytoplasmic GAF domain of KdpD may be responsible for intracellular K^+^ sensing. Most GAF domains are involved in regulatory functions through binding to cyclic nucleotides or other small ligands including ions (55, 56). Interestingly, we found that KdpD from *C. crescentus* lacks a GAF domain. Therefore, it is possible that such a connection between intracellular sensing and phosphatase activity does not exist in *C. crescentus*, which would explain why KdpD mainly acts as a phosphatase. Recently, it was reported that *E. coli* KdpD also works as serine kinase in high K^+^ concentrations to control KdpB ATPase activity and ultimately stop Kdp-dependent K^+^ transport (57). The phosphorylated serine residue in KdpB as well as the two Walker (A and B) domains in tandem driving the serine kinase activity of KdpD are highly conserved in *C. crescentus* (**Fig. S6**), indicating that this post-transcriptional control of K^+^ influx might also work in *C. crescentus*.

As mentioned above, in *E. coli*, the HK PhoR can crosstalk with the RR KdpE, which allows a Δ*kdpD* mutant to grow in a low K^+^ environment after a long lag phase (58). Since PhoR is conserved in *C. crescentus*, it constitutes a good candidate for the non-cognate HK phosphorylating KdpE. To test this hypothesis, we could measure growth behaviour as well as P*_kdp_* activity in a strain in which both *kdpD* and *phoR* were inactivated. PhoR is a sensor kinase regulating the phosphorylation levels and activity of the RR PhoB, which drives the Pho regulon, that is the entire set of genes whose expression relies on phosphate availability. Interestingly, a crosstalk between KdpD and PhoB was also described, which means that in the absence of PhoR, PhoB can be phosphorylated by KdpD in low phosphate conditions (58). The double crosstalk suggests there is tight connection between K^+^ and phosphate homeostasis. Since both TCS – KdpDE and PhoBR – are conserved in *C. crescentus*, it could be worth testing if the crosstalk exists in both directions as well.

Interestingly, we observed that Kup activity varies in excessive concentrations depending on the media. This indicates that Kup activity is regulated in toxic K^+^ concentrations depending on environmental resources and/or metabolic states. Also, suppressor mutations (A87P, G253S, and S456R) that bypass the growth defect of Δ*kdp* cells in limiting K^+^ conditions have been mapped in *kup* without impacting the growth when K^+^ was available. A plausible explanation would be that mutations found in Kup increase its affinity for K^+^, thereby increasing K^+^ transport to compensate the loss of Kdp, and that the efflux systems (KefB and KefC) limits the intracellular K^+^ concentration in abundant K^+^ conditions. In support of that, MD simulations have revealed mutation-driven structurally relevant changes in the putative K^+^ binding cavity of Kup. First, all mutants led to the flipping of E263 to a position closer to the one described for the corresponding E233 in KimA, which likely makes the second binding pocket accessible and would allow the coordination of another K^+^ cation. Indeed, the position observed for E263 in the WT Kup configuration appears to interfere with the second K^+^ binding site. Second, the carboxylate group of D146 was made more available for K^+^ coordination within this second binding site in a way similar to the one described for the corresponding D117 in KimA. Third, compared to the WT Kup, positions of Y61 and Y410 in the G253S and S456R mutants were also found to be shifted towards the positions observed for the corresponding Y43 and Y377 in KimA, thereby exposing to a greater extent their tyrosyl moiety towards the inside of the cavity where K^+^ ion is located. Such structural modifications would be induced by local reorganization of H bonds at the mutation sites, resulting in a cascade of subtle conformational changes towards the binding cavity. While the introduction of P87 abrogates two potential H bonds between A87 and the backbone of V83 and V84, S253 can form new H bonds with an adjacent α-helix thanks to its polar side chain (**Fig. S3BC**). Conversely, mutation of S456 into R456 decreases the number of intra-helix H bonds through the loss of the hydroxyl moiety (**Fig. S3D**).The available Cryo-EM structure of KimA likely represents an inward-occluded state in which two K^+^ cations are trapped in the intracellular cavity by the ideally positioned Y43, D117, E233 and Y377 residues (43). Thus, our *in silico* results are consistent with a higher K^+^ affinity observed experimentally in Kup mutants, likely due to the reorientation of the corresponding Y61, D146, E263 and Y410 residues into a KimA-like configuration, which consequently enhances K^+^ coordination.

Studies in other bacteria indicate Kup is a single membrane protein showing moderate affinity for K^+^ (59, 60). In *Lactococcus lactis,* the second messenger c-di-AMP binds to Kup homologues, in addition to the Trk homologue, to inhibit K^+^ transport (61). It is therefore possible that Kup activity is regulated by intracellular signalling molecules to coordinate K^+^ uptake with environmental conditions. KdpD homologues have also been reported to bind to c-di-AMP in other Gram-positive bacteria (62, 63). High levels of c-di-AMP inhibit the KdpD-dependent upregulation of the *kdp* operon in *Staphylococcus aureus* (63). On top of this, genes encoding K^+^ transporters are also regulated by a c-di-AMP-dependent riboswitch. The 5’ leader region (LR) of the *B. subtilis kimA* transcript can adopt two mutually exclusive conformations, one favouring transcription readthrough while the other promoting transcription termination (64). Upon direct binding of c-di-AMP, the *kimA* 5’ LR preferentially forms a structure exposing a predicted intrinsic transcription terminator, thereby inhibiting transcription elongation of *kimA*. Interestingly, a study showed that transcription levels of *kimA* were higher in low K^+^ conditions (0.1 mM KCl) than at higher K^+^ concentrations (5 mM KCl) (65). Indeed, the high-affinity K^+^ transporter KimA was exclusively detected in low K^+^ conditions. This K^+^-dependent control of *kimA* expression is suggested to be achieved through a modulation of c-di-AMP intracellular levels. Supporting this hypothesis, concentrations of c-di-AMP were increased by two-fold in high K^+^ conditions. More than just increasing in concentration, c-di-AMP becomes indispensable in high K^+^ conditions since a strain lacking c-di-AMP synthesizing genes is viable at low but not at high K^+^ concentrations. *C. crescentus* produces c-di-GMP, rather than c-di-AMP, to control cell development and division. If we speculate that c-di-GMP levels fluctuate reminiscently to that of c-di-AMP, K^+^-depletion would result in low c-di-GMP levels thereby preventing cell differentiation. Although it remains unclear how K^+^ limitation leads to a decrease in c-di-AMP levels, it has been shown that the enzymes producing c-di-AMP in *B. subtilis* were less abundant in low K^+^ conditions (65). Evaluating the growth behaviour of cells unable to synthesize c-di-GMP (66) in excessive or limiting K^+^ conditions could be an initial step to assess if such regulation is conserved.

The nitrogen-related phosphotransferase component EIIA^Ntr^ has been shown in *E. coli* to regulate not only K^+^ transport, by modulating KdpD kinase activity (67–69) but also phosphate starvation response, by modulating PhoR activity (70). For instance, in limiting K^+^ conditions (< 5 mM), the unphosphorylated form of EIIA^Ntr^ was shown to directly interact with KdpD and stimulate its kinase activity, thereby promoting phosphorylation of KdpE and the subsequent increase of *kdp* genes expression (71). This EIIA^Ntr^-based control of KdpD has also been reported in the plant symbionts *Rhizobium leguminosarum* (72) and *Sinorhizobium fredii* (8), two α-proteobacteria closely related to *C. crescentus*. It was later discovered that EIIA^Ntr^ regulates sigma factor selectivity by modulating K^+^ levels. Indeed, the competition between σ^70^ and σ^S^ for interaction with the RNA polymerase (RNAP) is influenced by K^+^ intracellular levels (73). Depending on the phosphorylation state of EIIA^Ntr^, the resulting K^+^ intracellular levels preferentially favour the binding of either σ^70^ or σ^S^ to the core RNAP (73). Interestingly, PTS^Ntr^components regulate the levels of another signalling molecule – (p)ppGpp – that connects cell division with environmental and metabolic cues in *C. crescentus* (74–76). Knowing that (p)ppGpp also influences gene expression by regulating sigma factor competition (77), it is therefore tempting to speculate that EIIA^Ntr^ and/or (p)ppGpp participate to the K^+^ homeostasis in *C. crescentus*, even in other α-proteobacteria.

## Methods

### Bacterial strains and growth conditions

All *Escherichia coli* strains used in this thesis were grown aerobically in Luria-Bertani (LB) broth (Sigma). *E. coli* Top10 was used for cloning purpose while *E. coli* MT607 was used for triparental matings. Thermo-competent cells were used for transformation of *E. coli*. All *Caulobacter crescentus* strains used are derived from the synchronizable wild-type strain NA1000. The traditional synthetic media supplemented with glucose (M2G) was modified by replacing potassium salts by their equivalent sodium salts to obtain a medium without any K^+^ (K_0_) to which the desired K^+^ concentrations could be added. The source of K^+^ consists in both KH_2_PO_4_ and K_2_HPO_4_ mixed to an equivalent ratio in terms of K^+^ concentration. For each experiment, cells were first grown at 30 °C; in Peptone Yeast Extract (PYE), then adapted overnight at 30 °C in K_0_ media supplemented with 0.5 mM K^+^, washed twice in K_0_ media and finally grown at 30 °C in K_0_ media supplemented with the desired K^+^ concentration. For growth curve measurements, cultures were diluted to a final OD_660_ of 0.02 in a 96-well plate. Growth was monitored during 48 h with continuous shaking at 30 °C, in an automated plate reader (Bioscreen C, Lab Systems) measuring OD_660_ every 10 min. For *E. coli*, antibiotics were used at the following concentrations (μg/ml; in liquid/solid medium): chloramphenicol (20/30), kanamycin (30/50), oxytetracycline (12.5/12.5), spectinomycin (100/100), streptomycin (50/100) while for *C. crescentus*, media were supplemented with kanamycin (5/20), oxytetracycline (2.5/2.5), nalidixic acid (20) when appropriate. Genes under the control of the inducible xylose promoter (P*_xylX_*) were induced with 0.1 % xylose. Plasmid delivery into *C. crescentus* was achieved by either bi- or tri-parental mating using *E. coli* S17-1 and *E. coli* MT607 as helper strains, respectively. In-frame deletions were created by using the pNPTS138-derivative plasmids as follows. Integration of the plasmids in the *C. crescentus* genome after single homologous recombination were selected on PYE plates supplemented with kanamycin. Three independent recombinant clones were inoculated in PYE medium without kanamycin and incubated overnight at 30 °C. Then, dilutions were spread on PYE plates supplemented with 3% sucrose and incubated at 30 °C. Single colonies were picked and transferred onto PYE plates with and without kanamycin. Finally, to discriminate between mutated and wild-type loci, kanamycin-sensitive clones were tested by PCR on colony using locus-specific oligonucleotides.

### Spotting assays

Ten-fold serial dilutions (in M2G-K) were prepared in 96-well plates from 5 ml cultures in standard universal tubes grown overnight at 30 °C in the corresponding media. Cells (5μl) were then spotted on plates, incubated at 30 °C for two-to-three days and pictures were taken.

### Flow cytometry analysis

DNA content was assessed using fluorescence activated cell sorting (FACS). Cells were fixed in ice-cold 77% ethanol. Fixed samples were then washed in FACS staining buffer (10mM Tris pH 7.2, 1mM EDTA, 50mM NaCitrate, 0.01% Triton X-100) then incubated for 30 min at room temperature in FACS staining buffer containing 0.1 mg/ml RNaseA. Cells were then collected by centrifugation for 3 min at 9000 rpm, resuspended in 1ml FACS staining buffer containing 0.5 mM Sytox Green Nucleic acid stain (Life Technologies), and incubated for 5 min at RT in the dark. Samples were analysed in flow cytometer (FACS Verse, BD Biosciences) at laser excitation of 488 nm. At least 1000 cells were recorded for each experiment. Before analysis, settings were adjusted with samples of a wild-type (RH50), a G1 accumulating strain (*spoT_D81G_*, RH1752) and a G2 accumulating strain (Δ*spoT,* RH1755) grown in PYE. Data were analysed with the BD FACSuite V1.0.5 software.

### Synchronization of cells

Cells were grown in 300 ml K_0_ media supplemented with 0.5 mM K^+^, harvested by centrifugation for 15 min at 6000 rpm, 4 °C; and resuspended in 45 ml of ice cold K_0_ media combined with 20 ml of ice cold Ludox® (Sigma-Aldrich; Ref 420808). After inverting several times, the cell-Ludox suspension was centrifuged for 35 min at 9000 rpm, 4 °C. Swarmer cells, corresponding to the bottom band, were collected, washed twice in ice-cold K_0_ media, and finally resuspended in K_0_ media supplemented with adequate K^+^ concentrations for growth at 30 °C. Samples were withdrawn every 20 minutes for 140 min for FACS analysis.

### β-galactosidase assays

β-galactosidase assays were performed as already described in (37). Briefly, cultures were either grown to stationary phase directly in a 96-wells plate or grown in flasks from which samples were withdrawn throughout growth and placed at −80 °C in between time points. Permeabilization of cells was performed by incubating cells with lysis buffer (20 mg/ml polymyxin B, β-mercaptoethanol) for 30 min at 28 °C. Then, 50 μl of 4 mg/ml O-nitrophenyl- β-D-galactopyranoside (ONPG) were added. Lysis and ONPG solutions were prepared using Z buffer as a solvent (60 mM Na_2_HPO_4_, 40 mM NaH_2_PO_4_, 10 mM KCl, 1 mM MgSO_4_, pH 7.0). Both OD_420_ and OD_550_ were measured every minute for 30 min at 30 °C using a fluorimeter (SpectraMax®, Molecular Devices). Miller Units were calculated using the following formula: M.U. = where t is the reaction time in min v is the volume of culture used in ml. Then, ONPG hydrolysis was measured at 30 °C for 30 min. The activity of the β-galactosidase expressed in miller units (MU) was calculated using the following equation: MU = (OD_420_ × 1,000) / [OD_660_ × t × v] where “t” is the time of the reaction (min), and “v” is the volume of cultures used in the assays (ml). Experimental values were the average of three independent experiments.

### Protein purification and production of polyclonal antibodies

To immunize rabbits for production of polyclonal antibodies, an *E. coli* BL21 (DE3) pLysS strain carrying plasmid pET-28a-*kdpE* was grown in LB medium supplemented with kanamycin and chloramphenicol until an OD600 of 0.5. After addition of IPTG to a final concentration of 1 mM, the culture was incubated at 18 °C for 18 hrs. Cells were then harvested by centrifugation for 30 min at 5,000 x g, 4 °C. The pellet was resuspended in 20 ml of binding buffer (20 mM Tris-HCl pH 8.0, 500 mM NaCl, 10% glycerol, 10 mM MgCl_2_, 12.5 mM Imidazole) supplemented with complete EDTA-free protease cocktail inhibitor (Roche), 400 mg lysozyme (Sigma) and 10 mg DNaseI (Roche) and incubated for 30 min on ice. Cells were then lysed by sonication. After centrifugation at 12,000 rpm for 30 min at 4°C, the lysate was loaded on a Ni-NTA column and incubated 1 hr at 4 °C with end-over-end agitation. The column was then washed with 5 ml binding buffer, 3 ml Wash1 buffer (binding buffer with 25 mM imidazole), 3 ml Wash2 buffer (binding buffer with 50 mM imidazole), 3 ml Wash3 buffer (binding buffer with 75 mM imidazole). Proteins bound to the column were eluted with 3 ml Elution buffer (binding buffer with 100 mM imidazole) and aliquoted in 300 µl fractions. All the fractions containing the protein of interest (checked by Coomassie blue staining) were pooled and dialyzed in Dialysis buffer (20 mM Tris pH 8, 500 mM NaCl, 10% glycerol). Purified KdpE was used to immunize rabbits (CER Groupe, Marloie, Belgium).

### Tn-seq analysis

A mini-Tn5 was introduced in *C. crescentus* NA1000 WT strain by bi-parental mating. Briefly, overnight cultures of recipient and donor strains (grown in LB supplemented with kanamycin) were mixed in 95:5 ratio to a final volume of 1 ml. Cells were harvested by centrifugation at 9,000 rpm (7,600 x g) at room temperature for 2 min. The supernatant was removed, and 1 ml of fresh PYE medium was added to wash the pellet. A second centrifugation step was done to remove again the supernatant. Thereafter, the pellet was resuspended in 50 μl and spotted on PYE agar and incubated at 30 ° C for 4h. Cells were collected, resuspended in 10 ml M2G or M2G-K and washed twice with the same volume. Dilutions (10^-6^) were plated on M2G or M2G-K plates supplemented with aztreonam (5μg ml^-1^) and kanamycin (5μg ml^-1^) and incubated at 30 ° C for 5 days. Then, at least 3 x10^5^ colonies were collected in M2G or M2G-K supplemented with 10% glycerol and stored at −80 °C. Genomic DNA was then extracted following the *Nucleospin Tissue Kit* protocol (Macherey-Nagel) and resuspended in 50 μl elution buffer (5 mM Tris-HCl [pH 8.5]). DNA sequencing was performed using an Illumina NextSeq (paired-end 2×75) instrument (Fasteris, Geneva, Switzerland). Reads corresponding to the mini-Tn5 insertion sites were first filtered with the 5’-GGTTGAGATGTGTATAAGAGACAG sequence before being processed as described in (78). Briefly, filtered reads were mapped on the genome of *C. crescentus* NA1000 (GenBank accession number NC_011916.1) and converted to Sequence Alignment/Map (SAM) format using the Burrows-Wheeler Aligner (BWA) and SAMtools, respectively, from the Sourceforge server (https://sourceforge.net/). Next, the number of reads overlapping each genomic position was computed using custom Python scripts. The total number of reads for internal 80% of each ORF in each condition were then computed using custom Python scripts. For comparing strains, the total number of reads was normalized by the ratio of the number of reads between the two strains. Ward’s clustering analysis was performed to define the fitness categories as previously reported.

### Chromatin immunoprecipitation followed by deep sequencing (ChIP-Seq assay)

ChIP-Seq was performed as already described in (78). Briefly, 80 ml of mid-log-phase cells (OD_660_ of 0.6) were cross-linked in 1% formaldehyde and 10 mM sodium phosphate (pH 7.6) at room temperature (RT) for 10 min and then for 30 min on ice. Cross-linking was stopped by addition of 125 mM glycine and incubated for 5 min on ice. Cells were washed twice in phosphate buffer solution (PBS; 137 mM NaCl, 2.7 mM KCl, 10 mM Na_2_HPO_4_, 1.8 mM KH_2_PO_4_, pH 7.4) resuspended in 450 μl TES buffer (10 mM Tris-HCl [pH 7.5], 1 mM EDTA, and 100 mM NaCl), and lysed with 2 μl of Ready-lyse lysozyme solution for 5 min at RT. Protease inhibitors (Roche) were added, and the mixture was incubated for 10 min. Then, 550 μl of ChIP buffer (1.1% Triton X-100, 1.2 mM EDTA, 16.7 mM Tris-HCl [pH 8.1], 167 mM NaCl, protease inhibitors) were added to the lysate and incubated at 37°C for 10 min before sonication (2 x 8 bursts of 30 sec on ice using a Diagenode Bioruptor) to shear DNA fragments to a length of 300 to 500 bp. Lysate was cleared by centrifugation for 10 min at 12,500 rpm at 4°C, and protein content was assessed by measuring the OD_280_. Then, 7.5 mg of proteins was diluted in ChIP buffer supplemented with 0.01% SDS and precleared for 1 hr at 4°C with 50 μl of SureBeads Protein A Magnetic Beads (BioRad) and 100 μg bovine serum albumin (BSA). One microliter of polyclonal anti-KdpE antibodies was added to the supernatant before overnight incubation at 4°C under gentle agitation. Next, 80 μl of BSA presaturated protein A-agarose beads were added to the solution and incubated for 2 hrs at 4°C with rotation, washed once with low-salt buffer (0.1% SDS, 1% Triton X-100, 2 mM EDTA, 20 mM Tris-HCl [pH 8.1], 150 mM NaCl), once with high-salt buffer (0.1% SDS, 1% Triton X-100, 2 mM EDTA, 20 mM Tris-HCl [pH 8.1], 500 mM NaCl), once with LiCl buffer (0.25 M LiCl, 1% NP-40, 1% deoxycholate, 1 mM EDTA, 10 mM Tris-HCl [pH 8.1]), and once with TE buffer (10 mM Tris-HCl [pH 8.1] 1 mM EDTA) at 4°C, followed by a second wash with TE buffer at RT. The DNA-protein complexes were eluted twice in 250 μl freshly prepared elution buffer (0.1 M NaHCO3, 1% SDS). NaCl was added at a concentration of 300 mM to the combined eluates (500 μl) before overnight incubation at 65°C to reverse the cross-link. The samples were treated with 20 μg of proteinase K in 40mM EDTA and 40 mM Tris-HCl (pH 6.5) for 2 h at 45°C. DNA was extracted using a Nucleospin PCR cleanup kit (Macherey-Nagel) and resuspended in 50 μl elution buffer (5 mM Tris-HCl [pH 8.5]). DNA sequencing was performed using an Illumina NextSeq 550 (paired-end 2×75) instrument (Seqalis). NGS data were analysed as described in (78).

### RNA-Seq

WT and Δ*kdpE* cells were grown overnight to OD_660_ ∼ 0.3 in M2G-K 0,025 mM K^+^ and WT in M2G-K 0,5 mM K^+^. Thereafter, total RNA was extracted with RNeasy® Protect Bacteria Kit from Qiagen and following manufacturer’s instructions. The quantity and quality (A260/A280 ration) of RNA was determined with a Thermo Scientific™ Nanodrop ™ One Microvolume UV-Vis Spectrophotometer. RNA-Seq TTRNA libraries were prepared according to the manufacturer’s instructions and sequenced with Illumina NovaSeq 6000 (paired-end 2×100) instrument (Seqalis). NGS data were analysed using Galaxy (https://usegalaxy.org) (37). Briefly, FastQC was used to evaluate the quality of the reads; HISAT2 was used to map the reads onto the NA1000 reference genome (NC_011916.1) and generate bam files; featureCounts was used to generate counts tables using bam files and DESeq2 was used to determine differentially expressed genes. The Volcano plot was generated using GraphPad Prism 9 software.

### MEME analysis

Sequences corresponding to the peaks founds in ChIP-seq were analysed for a KdpE conserved DNA binding motif using MEME (37) (**Table S6**). As searching parameters, the classical motif discovery mode, zoops motif site distribution, 10 minimum motif width and 20 maximum motif width, 2 as maximum number of motifs and a p-value<0.001 as cut-off were used. The motif with highest site count was selected.

### Statistical analyses

All the statistical analyses were performed using GraphPad Prism 9 software. A *P* value of <0.05 was considered as significant.

### Data availability

ChIP-Seq, RNA-Seq and Tn-Seq data have been deposited to the Gene Expression Omnibus (GEO) repository with the accession numbers [GSE253227, GSE253228, GSE253229].

### Molecular dynamics simulations

Protein systems were prepared for simulations from the three-dimensional model of Kup from *Caulobacter crescentus* (UniProt ID: Q9ABT9 or B8GXM1) generated with AlphaFold2 (79). A87P, G253S, and S456R mutants were manually edited from the WT structure at the corresponding position via the PyMOL software (version 2.5.2). As the putative potassium binding site of Kup is located within its transmembrane helixes, the N- (residues 1 to 38) and C-terminal (residues 499 to 665) domains were truncated to reduce computational costs. The resulting structural models (WT, A87P, G253S, and S456R) were properly protonated at pH 7.5 via the ProteinPrepare tool implemented in PlayMolecule (80). The CHARMMGUI suite was used for building the lipid bilayer, the insertion of Kup, and for the generation of input files using the Membrane Builder interface (81, 82). Based on the reported composition of *Caulobacter* inner membrane, a mix of 90% 1,2-dioleoyl-sn-glycero-3-phosphoglycerol (DOPG) and 10% tetraoleoyl cardiolipin (TOCL2) was selected to better reflect the native environment of Kup (41, 42). Each protein structure was oriented along the z axis, perpendicular to the lipid bilayer, with the transmembrane helix constituted from residue R77 to M104. The upper leaflet contained 65 DOPG and 7 TOCL2 molecules, whereas the lower leaflet was composed of 63 and 7 DOPG and TOCL2 molecules, respectively. KUP-embedded lipidic systems were solvated in TIP3P water, neutralized with K^+^ cations, and supplemented with 150 mM KCl. All-atom molecular dynamics (MD) simulations were carried out with the GROMACS 2023.1 suite package (83) on Kup WT, A87P, G253S, and S456R systems, using CHARMM36m force field ((84). Overall calculation efficiency was improved by hydrogen mass repartitioning (HMR), allowing to increase the time step to 4 fs. The energy of each system was first minimized within 5000 steps using the steepest descent method. Systems were further equilibrated successively at a temperature of 300 K and a pressure of 1 bar by the use of Berendsen thermostat and barostat, respectively, for a total of 1875 ps. Production MD were performed at 300 K and 1 bar with the Nosé-Hoover thermostat and the Parinello-Rahman barostat for 1 µs each. Long-range electrostatic interactions were taken into account by means of the Particle Mesh Ewald (PME) summation. Periodic boundary conditions (PBC) were set in all three dimensions, and atom trajectories were recorded every 4 ps. The final frame of each run was extracted and visualized with the PyMOL software (version 2.5.2). The latter was also used to create the presented snapshots and figures.

## Acknowledgments

We thank the members of the Hallez lab for critical reading of the manuscript and helpful discussions. The authors are also appreciative to the PTCI high-performance computing resource of the University of Namur. The present research benefited from computational resources made available on Lucia, the Tier-1 supercomputer of the Walloon Region, infrastructure funded by the Walloon Region under the grant agreement n°1910247. This work was supported by the Fonds de la Recherche Scientifique – FNRS (F.R.S. – FNRS) with a Welbio Starting Grant (WELBIO-CR-2019S-05) to R.H. A.Q-Y. was supported by a postdoctoral fellowship from the University of Namur (UNamur). L.L. and J.M. are both supported by the F.R.S. – FNRS, as a Postdoctoral Researcher for L.L. and a Research Fellow for J.M. C.M and R.H. are Senior Research Associates of the F.R.S. – FNRS.

