## Supplementary figures for "Regulation of potassium homeostasis in *Caulobacter crescentus*"

Figure S1

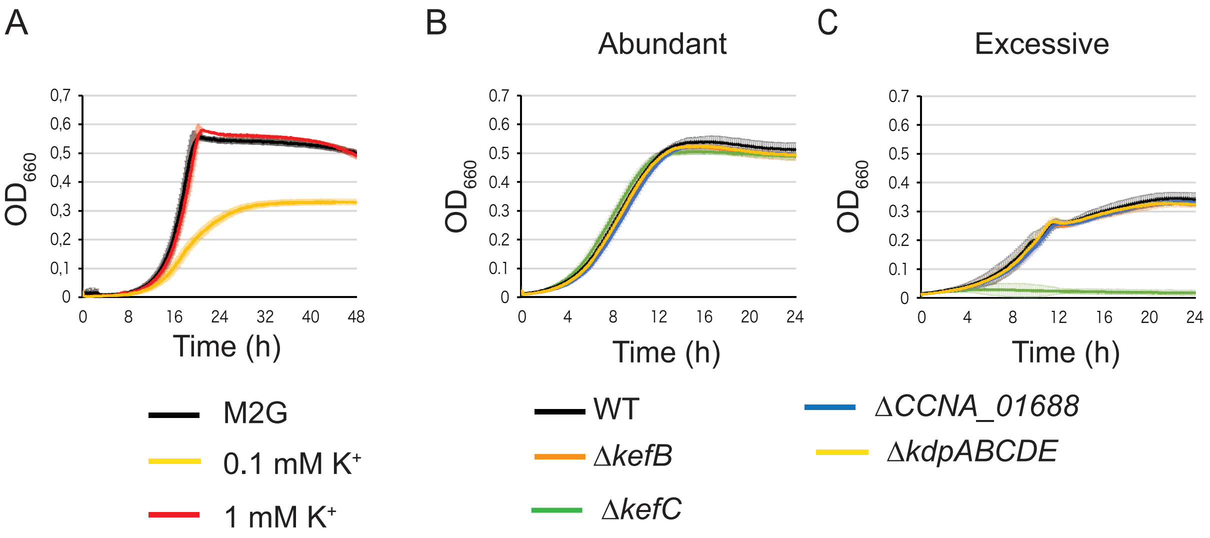


**Figure S1. Growth of *C. crescentus* with different $K^+$ concentrations.** (A) Growth of WT in M2G-K supplemented with different $K^+$ concentrations using KCl as $K^+$ source. Growth of WT, Δ*kefB*, Δ*kefC*, ΔCCNA_01688 and Δ*kdpABCDE* mutants in complex media PYE supplemented with (B) abundant (no extra $K^+$ added) and (C) excessive (50 mM) $K^+$ concentrations. Data represent average, n=3, and error bars= ±SD. Proportions of WT cells in G1, S and G2 phases at time 0, 60 and 120 min in M2G-K, 0.025 mM $K^+$ and 0.5 mM $K^+$ (D).

Figure S2

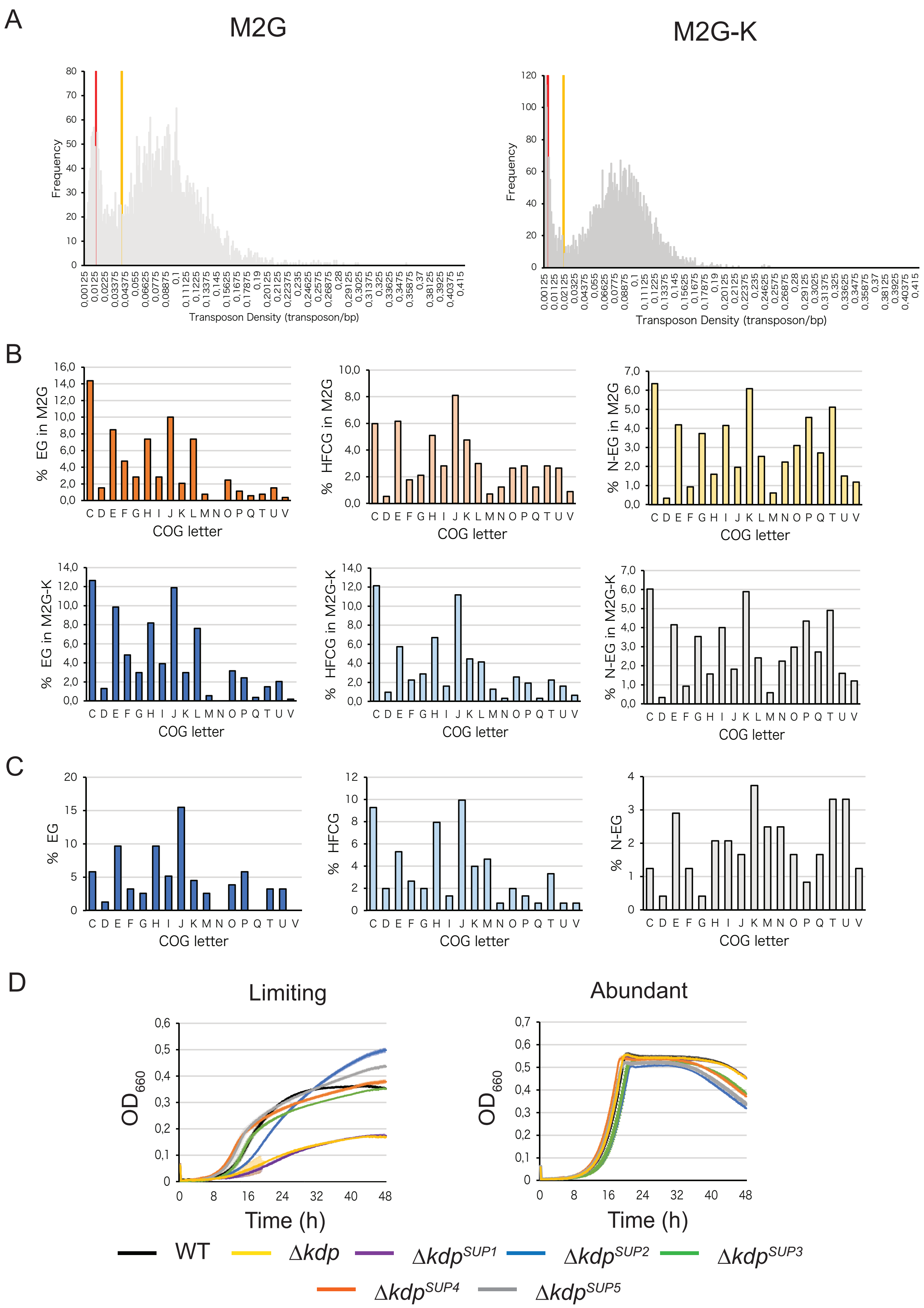

A
M2G
M2G-K
Frequency
Transposon Density (transposon/bp)
B
% EG in M2G
% HFCG in M2G
% N-EG in M2G
% EG in M2G-K
% HFCG in M2G-K
% N-EG in M2G-K
COG letter
C
% EG
% HFCG
% N-EG
COG letter
D
Limiting
Abundant
OD660
Time (h)
WT
Δkdp
ΔkdpSUP1
ΔkdpSUP2
ΔkdpSUP3
ΔkdpSUP4
ΔkdpSUP5

**Figure S2. Categories from the Tn-seq data and** Δ*kdp* suppressors fitness**.** (A) Transposon insertion frequencies and fitness-cost categories defined for genes of bacteria grown in M2G (abundant $K^+$) and M2G-K (limiting $K^+$) plates. Genes with transposon density up to the red line were considered as essential, while those between the red and yellow line as high-fitness cost, and those beyond the yellow line as non-essential. (B) Representation of COG categories for genes classified as essential, high-fitness cost and non-essential (EG, HFCG, N-EG, respectively) for cells grown in M2G and M2G-K media. (C) Same representation as in panel B, but for genes that changed in their fitness cost in analyses done in M2G and M2G-K media. COG categories: C, Energy production and conversion; D, Cell cycle control, cell division, chromosome partitioning; E, Amino acid transport and metabolism; F, Nucleotide transport and metabolism; G, Carbohydrate transport and metabolism; H, Coenzyme transport and metabolism; I, Lipid transport and metabolism; J, Translation, ribosomal structure and biogenesis; K, Transcription; L, Replication, recombination and repair; M, Cell wall/membrane/envelope biogenesis; N, Cell motility; O, Posttranslational modification, protein turnover, chaperones; P, Inorganic ion transport and metabolism; Q, Secondary metabolites biosynthesis, transport and catabolism; T, Signal transduction mechanisms; U, Intracellular trafficking, secretion and vesicular transport; and V, Defense mechanisms. Genes classified as R, General function prediction only; and S, Unknown function were not included in the data representation. (D) Growth of WT, Δ*kdp* and Δ*kdp* suppressors ($\Delta kdp^{SUP}$) in minimal media M2G-K supplemented with limiting (0.025 mM) or abundant (0.5 mM). Data represent average, n=3, and error bars= ±SD.

Figure S3

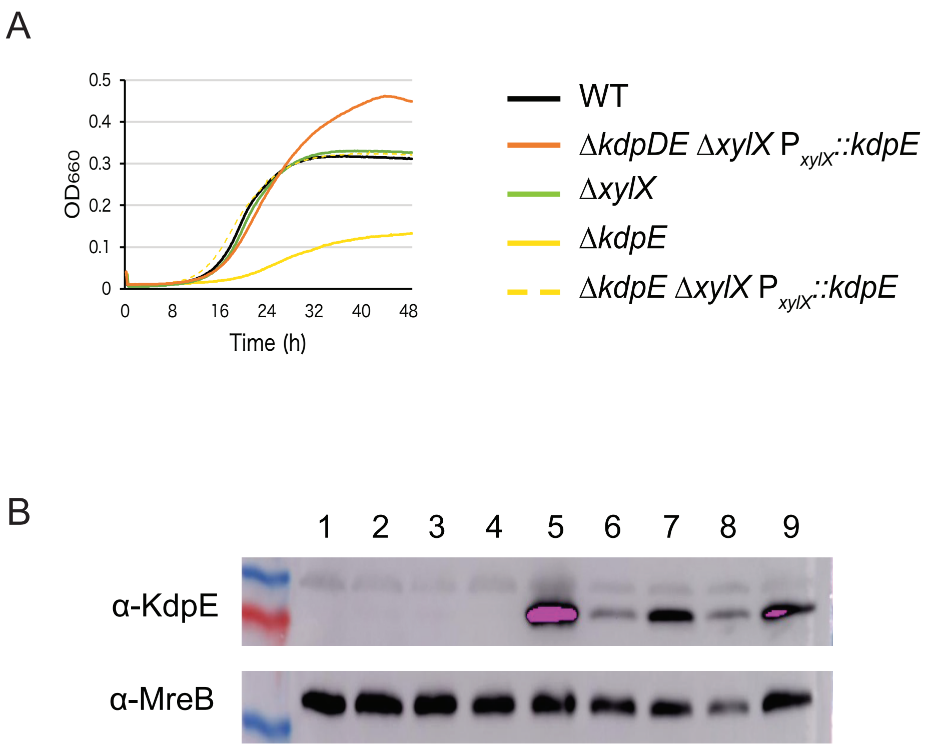


**Figure S3. Functionality of $P_{xylX}$::*kdpE* in Δ*kdpE* Δ*xylX* and Δ*kdpDE* Δ*xylX* strains for ChIP-seq experiments.** (A) Growth of WT, Δ*kdpE* $P_{xylX}$::*kdpE* Δ*xylX* and Δ*kdpDE* $P_{xylX}$::*kdpE* Δ*xylX* in M2G-K supplemented with 0.025 mM $K^+$ and 0.01% xylose. (B) Western blot detection of KdpE in Δ*kdpE* (1), Δ*kdpD* (2), WT (3, 4) Δ*kdpE* $P_{xylX}$::*kdpE* Δ*xylX* (5, 7), Δ*kdpDE* $P_{xylX}$::*kdpE* Δ*xylX* (6, 8) Δ*kdpE* $P_{xylX}$::*kdpE* (9). Protein crude extracts loaded in wells 1-3 and 9 come from cells grown in PYE, 4 in M2G, 5-6 in M2G-K supplemented with 0.025 mM $K^+$, 7-8 in in M2G-K supplemented with 0.5 mM $K^+$.

Figure S4

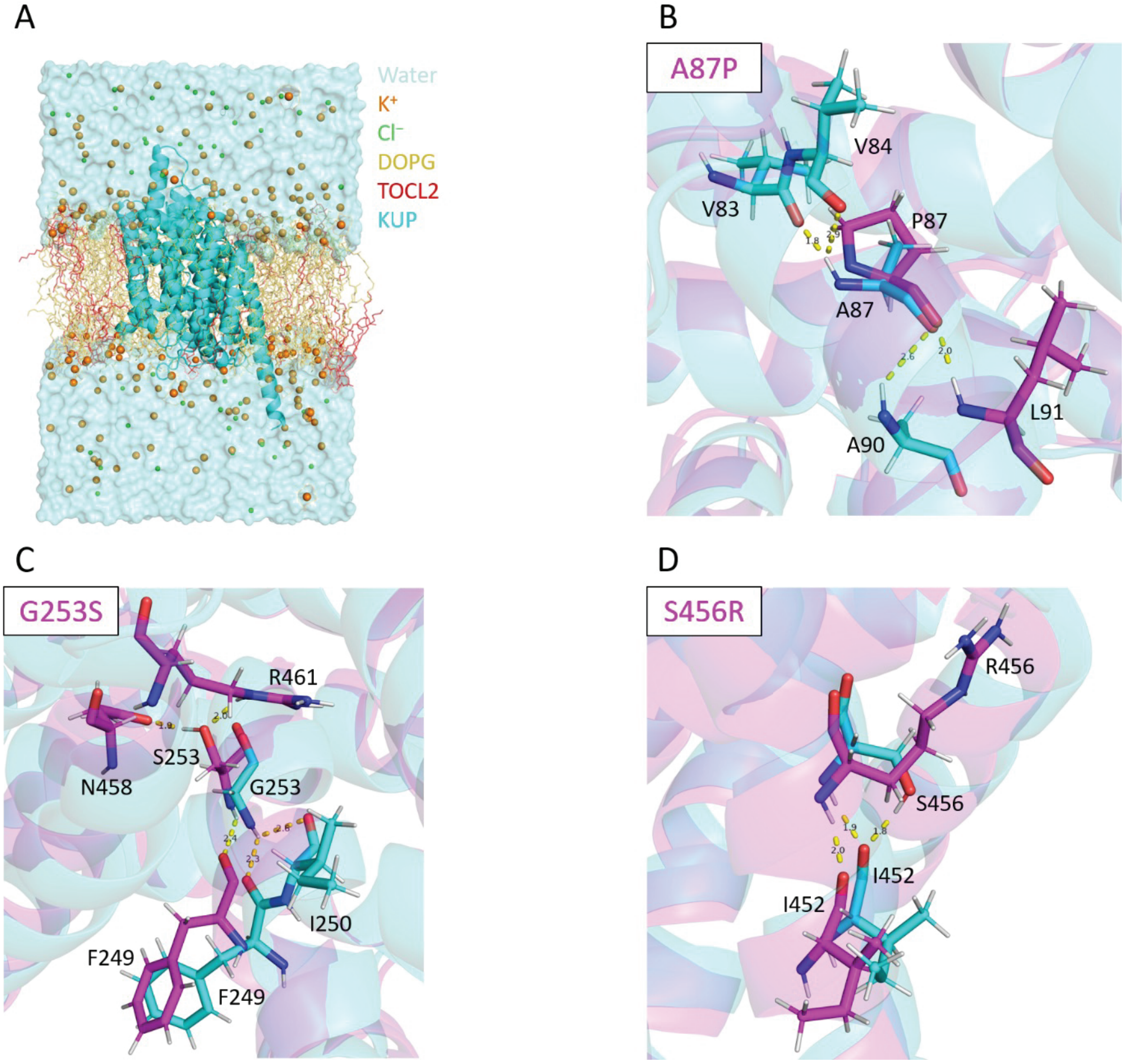


**Figure S4 Molecular dynamics-highlighted structural changes of mutated sites.** (A) Representation of the Kup transmembrane domains embedded in a DOPG-TOCL2 lipid bilayer solvated in water and 150 mM KCl at the initial state of the simulation after the minimization and equilibration steps. Water molecules are represented as van der Waals surface in pale blue, $K^+$ and $Cl^-$ ions as spheres in orange and green, respectively, DOPG and TOCL2 as lines in yellow and red, respectively, and Kup as cartoon in cyan. (B) Snapshot of the mutated site of Kup A87P, (C) G253S, and (D)

S456R after 1 $\mu$s of simulation. On each of these panels, aligned Kup WT (cyan) and mutated Kup (magenta) are shown as cartoon, and structurally relevant residues are represented as sticks in the corresponding color. Presence of H bonds is indicated by yellow dashes with their associated length in Å.

### Figure S5

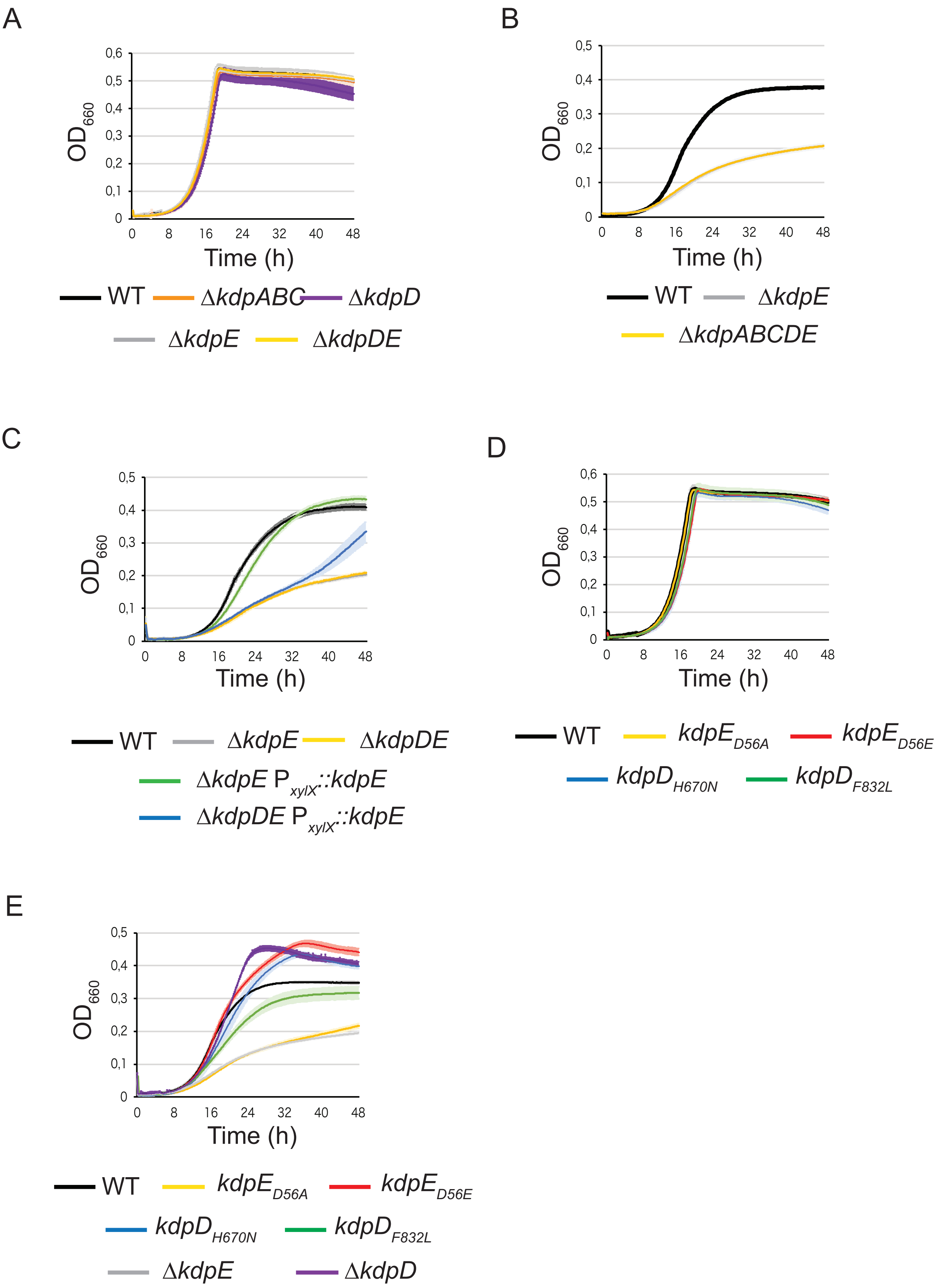
A
B
C
D
E
$OD_{660}$
Time (h)
0 8 16 24 32 40 48
0 0,1 0,2 0,3 0,4 0,5 0,6
WT
ΔkdpABC
ΔkdpD
ΔkdpE
ΔkdpDE
ΔkdpABCDE
ΔkdpE $P_{xylX}$::kdpE
ΔkdpDE $P_{xylX}$::kdpE
$kdpE_{D56A}$
$kdpE_{D56E}$
$kdpD_{H670N}$
$kdpD_{F832L}$

**Figure S5. Additional characterisation of *kdp* mutants.** (A) Growth of *kdp* mutants in minimal media M2G-K supplemented with abundant (0.5 mM) $K^+$ concentration. (B) Comparison of growth between Δ*kdpE* and Δ*kdpABCDE* mutants in minimal media M2G-K supplemented with limiting (0.025 mM) $K^+$ concentration. (C) Growth of Δ*kdpE* $P_{xylX}$::*kdpE* in minimal media M2G-K supplemented with limiting (0.025 mM) $K^+$ concentration and without xylose. (D) Growth of *kdpD* and *kdpE* catalytic mutants in minimal media M2G-K supplemented with abundant (0.5 mM) $K^+$ concentration. (E) Comparison of growth between catalytic point and KO mutants of *kdpD and kdpE* in minimal media M2G-K supplemented with limiting (0.025 mM) $K^+$ concentration.

Figure S6

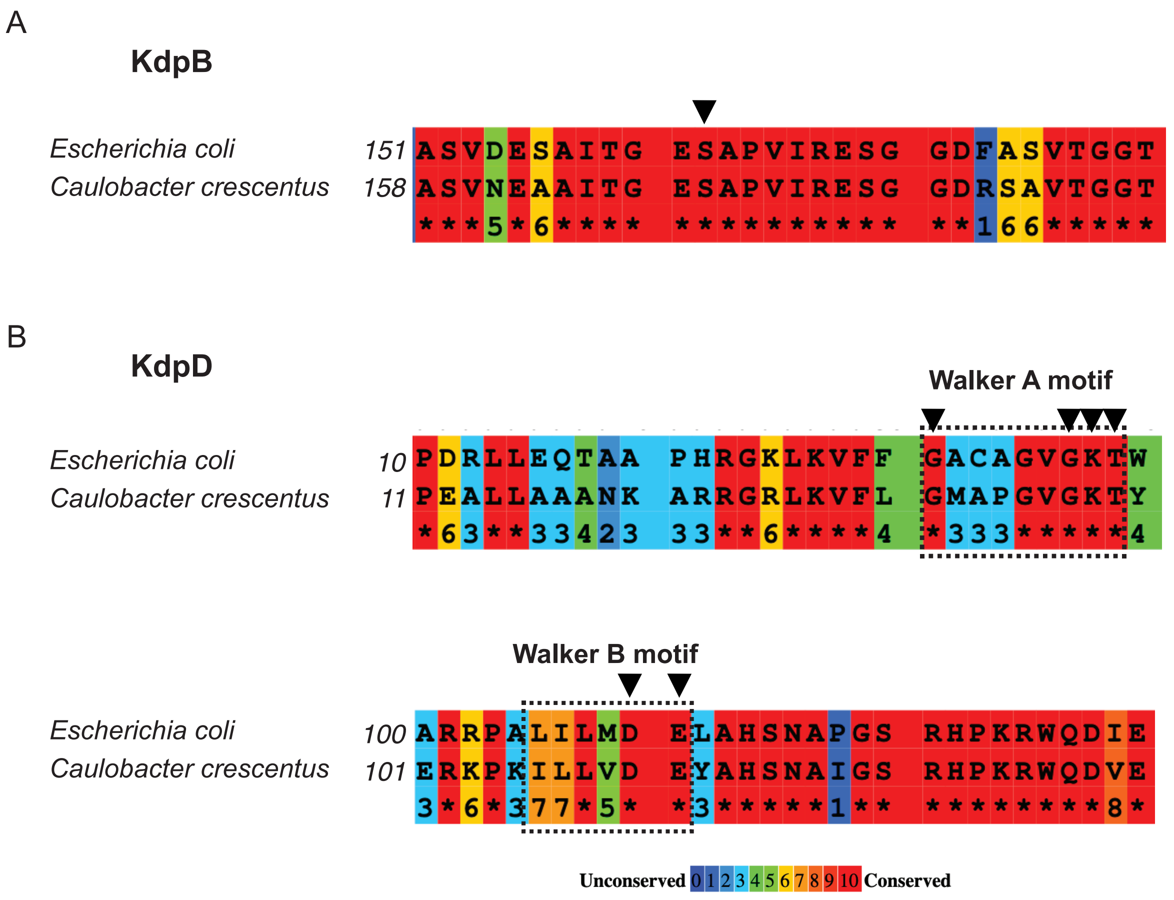

**Figure S6. Sequence conservation of KdpB and KdpD.** (A) The phosphorylated serine residue in *Escherichia coli* KdpB (black arrowheads) is conserved in *Caulobacter crescentus*. (B) The Walker A and B motifs in *Escherichia coli* KdpD (dotted lines) are conserved in *Caulobacter crescentus*. Black arrowheads point to critical residues.
