## Supplementary tables and methods for "Regulation of potassium homeostasis in *Caulobacter crescentus*"

Quintero-Yanes *et al.,* 2024.

Table S1A. Bacterial strains.

| Name | Description and relevant genotype | Reference/Source |
|---|---|---|
| **E. coli strains** | | |
| RH317 | DH10B | Life technology |
| MT607 | helper strain | Casadaban & Cohen. 1980 |
| Top10 | Thermocompetent cells | Life technology |
| ***Caulobacter crescentus* strains** | | |
| WT | Lab NA1000 strain | |
| Δ*kdpD* | NA1000 Δ*kdpD* | This study |
| Δ*kdpABC* | NA1000 Δ*kdpABC* | This study |
| Δ*kdpE* | NA1000 Δ*kdpE* | This study |
| Δ*kdpABCDE* | NA1000 Δ*kdpABCDE* | This study |
| Δ*kefB* | NA1000 Δ*kefB* | This study |
| Δ*kefC* | NA1000 Δ*kefC* | This study |
| ΔCCNA_01688 | NA1000 ΔCCNA_01688 | This study |
| Δ*kup* | NA1000 Δ*kup* | This study |
| $kdpE_{D56E}$ | NA1000 $kdpE_{D56E}$ | This study |
| $kdpE_{D56A}$ | NA1000 $kdpE_{D56A}$ | This study |
| $kdpD_{H670N}$ | NA1000 $kdpD_{H670N}$ | This study |
| $kdpD_{F832L}$ | NA1000 $kdpE_{F832L}$ | This study |
| WT $P_{kdp}$::*lacZ* | NA1000 pRKlac290 $P_{kdp}$::*lacZ* | This study |
| Δ*kdpD* $P_{kdp}$::*lacZ* | NA1000 Δ*kdpD* pRKlac290 $P_{kdp}$::*lacZ* | This study |
| Δ*kdpABC* $P_{kdp}$::*lacZ* | NA1000 Δ*kdpABC* pRKlac290 $P_{kdp}$::*lacZ* | This study |
| Δ*kdpE* $P_{kdp}$::*lacZ* | NA1000 Δ*kdpE* pRKlac290 $P_{kdp}$::*lacZ* | This study |
| $kdpE_{D56E}$ $P_{kdp}$::*lacZ* | NA1000 $kdpE_{D56E}$ pRKlac290 $P_{kdp}$::*lacZ* | This study |
| $kdpE_{D56A}$ $P_{kdp}$::*lacZ* | NA1000 $kdpE_{D56A}$ pRKlac290 $P_{kdp}$::*lacZ* | This study |
| $kdpD_{H670N}$ $P_{kdp}$::*lacZ* | NA1000 $kdpD_{H670N}$ pRKlac290 $P_{kdp}$::*lacZ* | This study |
| $kdpD_{F832L}$ $P_{kdp}$::*lacZ* | NA1000 $kdpE_{F832L}$ pRKlac290 $P_{kdp}$::*lacZ* | This study |
| WT EV | NA1000 pMR10 | This study |
| Δ*kdpD* EV | NA1000 Δ*kdpD* pMR10 | This study |
| Δ*kdpD* $kdpD^{+}$ | NA1000 Δ*kdpD* pMR10 $P_{kdp}$::*kdpD* | This study |
| Δ*kdpE* $kdpE^{++}$ | NA1000 Δ*kdpE* $P_{xyl}$::*kdpE* | This study |
| Δ*kdpDE* $kdpE^{++}$ | NA1000 Δ*kdpDE* $P_{xyl}$::*kdpE* | This study |
| $kup_{A87P}$ | NA1000 $kup_{A87P}$ | This study |
| $kup_{G253S}$ | NA1000 $kup_{G253S}$ | This study |
| $kup_{S456R}$ | NA1000 $kup_{S456R}$ | This study |
| Δ*kdpABCDE* $kdpkup_{A87P}$ | NA1000 Δ*kdpABCDE* $kup_{A87P}$ | This study |
| Δ*kdpABCDE* $kup_{G253S}$ | NA1000 Δ*kdpABCDE* $kup_{G253S}$ | This study |
| Δ*kdpABCDE* $kup_{S456R}$ | NA1000 Δ*kdpABCDE* $kup_{S456R}$ | This study |

Table S1B. Plasmids.

| Description | Reference/Source |
|---|---|
| pNPTS138 | M. R. Alley, Imperial College London (UK), unpublished |
| pXC-5 | Thanbichler *et al.,* 2007 |
| pMR10 | Shapiro, L. |
| pRKlac290 | Gober & Shapiro, 1992 |
| pRKlac290-$P_{pilA}$::*lacZ* | Skerker & Shapiro, 2000 |
| pNPTS138-Δ*kdpD* | This study |
| pNPTS138-Δ*kdpABC* | This study |
| pNPTS138-Δ*kdpE* | This study |
| pNPTS138-Δ*kdpDE* | This study |
| pNPTS138-Δ*kdpABCDE* | This study |
| pNPTS138-Δ*kefB* | This study |
| pNPTS138-Δ*kefC* | This study |
| pNPTS138-ΔCCNA_01688 | This study |
| pNPTS138-Δ*kup* | This study |
| pNPTS138-$kdpE_{D56E}$ | This study |
| pNPTS138-$kdpE_{D56A}$ | This study |
| pNPTS138-$kdpE_{H670N}$ | This study |
| pNPTS138-$kdpE_{F832L}$ | This study |
| pNPTS138-$kup_{A87P}$ | This study |
| pNPTS138-$kup_{G253S}$ | This study |
| pNPTS138-$kup_{S456R}$ | This study |
| pRKlac290-$P_{kdp}$::*lacZ* | This study |
| pMR10-$P_{kdp}$::*kdpD* | This study |
| pXC-5-$P_{xyl}$::*kdpE* | This study |
| pET-28a-*kdpE* | This study |

### Table S1C. Oligonucleotides.

| ID | Description | 5' to 3' sequence |
|---|---|---|
| 1990 | *kdpA*_for1 | ttagttacttaggggatccatggtcctgggttcgagtcc |
| 1991 | *kdpA*_rev1 | cctaagtaactaagaattcctcagggtcagggcgatttc |
| 2000 | *kdpC*_for1 | ttagttacttagggaattccactcgccagccagaaggac |
| 2001 | *kdpC*_rev1 | cctaagtaactaaaagcttatgtcgtcggcgcgagagag |
| 2002 | *kdpD*_for1 | ttagttacttaggaagcttgttcaccgtcctgctcg |
| 2003 | *kdpD*_rev1 | cctaagtaactaagaattctccggtcgtttcgggtcgtc |
| 2004 | *kdpD*_for2 | ttagttacttagggaattcgggcacgcggattctcatcag |
| 2005 | *kdpD*_rev2 | cctaagtaactaaggatcctttgggcgacagcttcacg |
| 2006 | *kefB*_for1 | ttagttacttaggggatccggcgtcaccttcaaggatc |
| 2007 | *kefB*_rev1 | cctaagtaactaagaattcatcaggtagcccagcaccgag |
| 2008 | *kefB*_for2 | ttagttacttagggaattccagcctggacgaagagctgc |
| 2009 | *kefB*_rev2 | cctaagtaactaaaagcttctgacgccgctgacggtctatc |
| 2010 | *kefC*_for1 | ttagttacttaggggatccgggtgaggaggaagcagtc |
| 2011 | *kefC*_rev1 | cctaagtaactaagaattcgtcgccaggaacaggaccag |
| 2012 | *kefC*_for2 | ttagttacttagggaattccgcgacgagtaccgcaagaag |
| 2013 | *kefC*_rev2 | cctaagtaactaaaagcttgggcgtctcgggcagaattg |
| 2014 | *kup*_for1 | ttagttacttaggggatcctgatggtctcggcgggcttg |
| 2015 | *kup*_rev1 | cctaagtaactaagaattcccgatgtcgccgaacaccac |
| 2016 | *kup*_for2 | ttagttacttagggaattcgccaacccgacggacttcttc |

| | | |
|---|---|---|
| 2017 | *kup*_rev2 | cctaagtaactaaaagcttaggcgagcacgaccttcttc |
| 2018 | *CCNA_01688*_for1 | ttagttacttaggggatccttcagcaagggcgcgaagg |
| 2019 | *CCNA_01688*_rev1 | cctaagtaactaagaattcgaaggccaggttgcggaagg |
| 2020 | *CCNA_01688*_for2 | ttagttacttagggaattctggcggtcctctccaagctg |
| 2021 | *CCNA_01688*_rev2 | cctaagtaactaaaagcttgtcaccctcgaaccgcatgtc |
| 2034 | *kdpD_rev* | ggatccccgggtactgcagtcatagcaggtgatgcggcgtcgtg |
| 2072 | *kdpE*_for1 | ggatccccgggtaggatccgctatgtcgccaacctgctcg |
| 2073 | *kdpE*_for2 | ggatccccgggtagaattctcgatgacgaggatgcggtg |
| 2074 | *kdpE*_rev1 | cctaagtaactaagaattcgaccgaacctggcgtaggctatc |
| 2075 | *kdpE*_rev2 | cctaagtaactaaaagcttctcgtaacgcacaagcacccagg |
| 2113 | *kdpE*_rev2' | cctaagtaactaaggatccctcgtaacgcacaagcacccagg |
| 2153 | *kdpE_for* | ttagttacttaggcatatgagcgcgcttcgccaccgcat |
| 2154 | *kdpE_rev* | cctaagtaactaagagctctgccggtagtcttggcgacg |
| 2171 | P$_{kdp}$*_for* | ttagttacttaggggtaccctggatcagcgacaggtcgg |
| 2172 | P$_{kdp}$*_rev* | cctaagtaactaaaagcttggaattggtccgtcagaacttgtcg |
| 2588 | *kdpD*$_{H670N}$*_for1* | acgcgtcacggccgaagctaaaagtgtttctgggcatgg |
| 2589 | *kdpD*$_{H670N}$*_for2* | tgaactcggtcagcaatgacctgcgcacgc |
| 2590 | *kdpD*$_{H670N}$*_rev1* | gcgtgcgcaggtcattgctgaccgagttca |
| 2591 | *kdpD*$_{H670N}$*_rev2* | gcaggatatctggatccacaggtgatgcggcgtcgtga |
| 2592 | *kdpE*$_{D56E}$*/kdpE*$_{D56A}$*_for1* | acgcgtcacggccgaagctaacgatgaaccgcaaatccat |
| 2593 | *kdpE*$_{D56A}$*_for2* | tggtgctggctctgggcttgcccgacat |
| 2594 | *kdpE*$_{D56A}$*_rev1* | atgtcgggcaagcccagagccagcacca |
| 2595 | *kdpE*$_{D56E}$*/kdpE*$_{D56A}$*_rev2* | gcaggatatctggatccaccctggagccgatagcctacg |
| 2596 | *kdpE*$_{D56E}$*_for2* | tggtgctggaactgggcttgcccgacat |
| 2597 | *kdpE*$_{D56E}$*_rev1* | atgtcgggcaagcccagttccagcacca |
| 3270 | *kup_for* | taagtaactaaggatccgcttccgaagccccccacgc |
| 3271 | *kup_rev* | taagtaactaaggtaccgcttccgaagccccccacgc |
| 3788 | *kdpE*$_{F832L}$*_for1* | gccgaagctagcgaattcgtggatccagatcggctctgatgaactcggtc |
| 3789 | *kdpE*$_{F832L}$*_rev1* | ctcctccatgcggcgtaacttgtcgaacac |
| 3790 | *kdpE*$_{F832L}$*_for2* | gtgttcgacaagttacgccgcatggaggag |
| 3791 | *kdpE*$_{F832L}$*_rev2* | gccggctggcgccaagcttctctgcaggataactctttgggcgacagc |

### Table S2. Genes changing in fitness category in the M2G and M2G-K Tn-seq analysis.

| GeneID | Name | M2G #Insertions | M2G #Insertions/pb | M2G-K #Insertions | M2G-K #Insertions/pb | COG Letter |
|---|---|---|---|---|---|---|
| CCNA_02026 | hypothetical protein | 0 | 0 | 0,917117309 | 0,005095096 | C |
| *CCNA_03639* | *ferredoxin, 2Fe-2s* | *0* | 0 | 1,834234617 | 0,007164979 | K |
| CCNA_03918 | hypothetical protein | 0 | 0 | 0,917117309 | 0,008491827 | NA |
| CCNA_R0161 | small non-coding RNA | 0 | 0 | 0,917117309 | 0,014557418 | NA |
| CCNA_R0038 | tRNA Pro | 0 | 0 | 0,917117309 | 0,015285288 | NA |
| CCNA_R0036 | tRNA Glu | 0 | 0 | 0,917117309 | 0,015285288 | NA |
| CCNA_R0182 | small non-coding RNA | 0 | 0 | 1,834234617 | 0,018527622 | NA |
| CCNA_00149 | transcriptional regulator, Cro/CI family | 0 | 0 | 3,668469234 | 0,020267786 | NA |

| | | | | | | |
|---|---|---|---|---|---|---|
| *CCNA_00808* | *LSU ribosomal protein L34P* | *0* | 0 | 2,751351926 | 0,025475481 | NA |
| CCNA_R0198 | small non-coding RNA | 0 | 0 | 1,834234617 | 0,02583429 | NA |
| CCNA_R0039 | tRNA Arg | 0 | 0 | 1,834234617 | 0,030570577 | NA |
| CCNA_R0033 | tRNA Asn | 0 | 0 | 1,834234617 | 0,030570577 | NA |
| CCNA_01581 | hypothetical protein | 0 | 0 | 8,254055777 | 0,031030285 | NA |
| CCNA_R0131 | small non-coding RNA | 0 | 0 | 1,834234617 | 0,03743336 | NA |
| CCNA_R0139 | small non-coding RNA | 0 | 0 | 3,668469234 | 0,05822967 | NA |
| CCNA_03737 | YggU superfamily protein | 0 | 0 | 14,67387694 | 0,062177445 | NA |
| CCNA_R0136 | small non-coding RNA | 0 | 0 | 4,585586543 | 0,066457776 | NA |
| CCNA_01255 | hypothetical protein | 0 | 0 | 8,254055777 | 0,069361813 | NA |
| CCNA_03919 | hypothetical protein | 0 | 0 | 4,585586543 | 0,07164979 | NA |
| CCNA_00739 | hypothetical protein | 0 | 0 | 5,502703851 | 0,084656982 | NA |
| CCNA_R0150 | small non-coding RNA | 0 | 0 | 6,41982116 | 0,085597615 | NA |
| CCNA_01482 | hypothetical protein | 0 | 0 | 6,41982116 | 0,085597615 | NA |
| CCNA_03934 | hypothetical protein | 0 | 0 | 3,668469234 | 0,101901923 | NA |
| CCNA_00147 | hypothetical protein | 0 | 0 | 13,75675963 | 0,122828211 | S |
| CCNA_03770 | malate dehydrogenase | 1 | 0,001298701 | 4,585586543 | 0,005955307 | C |
| CCNA_01812 | 2-C-methyl-D-erythritol 4-phosphate cytidylyltransferase/2-C-methyl-D-erythritol 2,4-cyclodiphosphat | 2 | 0,002178649 | 4,585586543 | 0,004995192 | I |
| CCNA_01321 | LSU ribosomal protein L6P | 1 | 0,00234192 | 2,751351926 | 0,006443447 | J |
| CCNA_01968 | hypothetical protein | 2 | 0,002877698 | 4,585586543 | 0,006597966 | S |
| CCNA_00678 | LSU ribosomal protein L11P | 1 | 0,002898551 | 1,834234617 | 0,005316622 | J |
| CCNA_03869 | chromosome partitioning protein parA | 2 | 0,00311042 | 5,502703851 | 0,00855786 | D |
| CCNA_00052 | predicted metal-dependent hydrolase | 1 | 0,003289474 | 1,834234617 | 0,006033667 | NA |
| CCNA_02591 | glutaredoxin | 1 | 0,003663004 | 3,668469234 | 0,013437616 | O |
| ***CCNA_01305*** | ***SSU ribosomal protein S10P*** | ***1*** | 0,004065041 | 3,668469234 | 0,014912477 | J |
| CCNA_01624 | orotate phosphoribosyltransferase | 2 | 0,004273504 | 2,751351926 | 0,005878957 | F |
| CCNA_01409 | delta-aminolevulinic acid dehydratase | 4 | 0,004975124 | 5,502703851 | 0,006844159 | H |
| CCNA_01749 | acyl carrier protein | 1 | 0,005319149 | 0,917117309 | 0,004878284 | IQ |
| CCNA_00309 | cytosolic protein | 1 | 0,005347594 | 1,834234617 | 0,009808741 | NA |
| CCNA_03957 | SirA/YedF/YeeD family regulatory protein | 1 | 0,005347594 | 14,67387694 | 0,07846993 | NA |
| CCNA_02640 | phospho-N-acetylmuramoyl-pentapeptide- transferase | 5 | 0,005617978 | 6,41982116 | 0,007213282 | M |
| CCNA_00100 | ribulose-phosphate 3-epimerase | 3 | 0,0056926 | 2,751351926 | 0,005220782 | NA |
| *CCNA_03373* | *thiamin-phosphate pyrophosphorylase* | *3* | 0,005813953 | 2,751351926 | 0,005332077 | H |
| CCNA_01043 | hypothetical protein | 1 | 0,005847953 | 11,92252501 | 0,069722368 | NA |

| | | | | | | |
|---|---|---|---|---|---|---|
| CCNA_01987 | hypothetical protein | 4 | 0,005934718 | 3,668469234 | 0,005442833 | S |
| CCNA_03643 | succinate dehydrogenase membrane anchor subunit | 2 | 0,005970149 | 1,834234617 | 0,005475327 | C |
| CCNA_01484 | hypothetical protein | 2 | 0,005970149 | 6,41982116 | 0,019163645 | NA |
| CCNA_00761 | hypothetical protein | 1 | 0,005988024 | 0,917117309 | 0,00549172 | K |
| *CCNA_02789* | *transcriptional regulator, Xre family* | *1* | 0,005988024 | 1,834234617 | 0,010983441 | NA |
| CCNA_00196 | 3-isopropylmalate dehydratase large subunit | 7 | 0,006081668 | 7,336938469 | 0,006374404 | E |
| CCNA_02596 | SSU ribosomal protein S4P | 3 | 0,006085193 | 2,751351926 | 0,005580836 | J |
| CCNA_01896 | hypothetical protein | 1 | 0,006097561 | 15,59099425 | 0,095067038 | NA |
| *CCNA_03865* | *leucyl-tRNA synthetase* | *13* | 0,006283229 | 18,34234617 | 0,00886532 | J |
| CCNA_02706 | hypothetical protein | 2 | 0,006289308 | 4,585586543 | 0,014420083 | NA |
| CCNA_03350 | hypothetical protein | 1 | 0,006369427 | 22,01081541 | 0,140196276 | F |
| CCNA_03144 | DNA primase | 10 | 0,006472492 | 9,171173086 | 0,005936034 | L |
| CCNA_03702 | SSU ribosomal protein S1P | 9 | 0,00658376 | 11,0054077 | 0,008050774 | J |
| CCNA_02033 | NADH-quinone oxidoreductase chain A | 2 | 0,006644518 | 2,751351926 | 0,009140704 | C |
| CCNA_00972 | hypothetical protein | 1 | 0,006711409 | 11,92252501 | 0,080016946 | NA |
| CCNA_00874 | biotin synthesis protein bioC | 5 | 0,006858711 | 5,502703851 | 0,007548291 | H |
| CCNA_03879 | uroporphyrinogen decarboxylase | 6 | 0,006872852 | 5,502703851 | 0,006303212 | NA |
| CCNA_R0129 | small non-coding RNA | 1 | 0,006944444 | 2,751351926 | 0,019106611 | NA |
| CCNA_01176 | hypothetical protein | 2 | 0,006968641 | 2,751351926 | 0,009586592 | NA |
| CCNA_00341 | succinyl-CoA synthetase subunit alpha | 5 | 0,007062147 | 4,585586543 | 0,006476817 | C |
| CCNA_00918 | peptide release factor-glutamine N5-methyltransferase | 5 | 0,007194245 | 4,585586543 | 0,006597966 | J |
| CCNA_02389 | glucosamine-1-phosphate acetyltransferase/UDP-N-acetylglucosamine pyrophosphorylase | 8 | 0,007207207 | 6,41982116 | 0,005783623 | M |
| CCNA_03787 | heme exporter protein CcmD | 1 | 0,007518797 | 1,834234617 | 0,013791238 | NA |
| CCNA_01942 | Rrf2 family protein | 3 | 0,007894737 | 1,834234617 | 0,004826933 | K |
| CCNA_01633 | GTP-binding protein era | 6 | 0,007894737 | 3,668469234 | 0,004826933 | R |
| CCNA_02248 | hypothetical protein | 1 | 0,008196721 | 8,254055777 | 0,067656195 | NA |
| CCNA_00373 | ATP synthase A chain | 5 | 0,008210181 | 3,668469234 | 0,006023759 | C |
| CCNA_02217 | hypothetical protein | 2 | 0,008264463 | 6,41982116 | 0,026528187 | NA |
| CCNA_03359 | phosphoglycerate kinase | 8 | 0,008403361 | 8,254055777 | 0,008670227 | G |
| CCNA_03704 | 3-phosphoshikimate 1-carboxyvinyltransferase | 9 | 0,008450704 | 5,502703851 | 0,005166858 | E |
| CCNA_02644 | putative cell division protein | 3 | 0,008450704 | 1,834234617 | 0,005166858 | NA |
| CCNA_03766 | 16S rRNA processing protein rimM | 4 | 0,008602151 | 9,171173086 | 0,019722953 | J |
| CCNA_00098 | hypothetical protein | 2 | 0,00862069 | 19,25946348 | 0,083014929 | NA |

| | | | | | | |
|---|---|---|---|---|---|---|
| CCNA_02526 | dihydroorotase | 9 | 0,008645533 | 8,254055777 | 0,007928968 | F |
| CCNA_02520 | aspartyl/glutamyl-tRNA(Asn/Gln) amidotransferase subunit C | 2 | 0,008733624 | 1,834234617 | 0,008009758 | J |
| CCNA_01758 | 4-hydroxythreonine-4-phosphate dehydrogenase PdxA | 7 | 0,008739076 | 4,585586543 | 0,005724827 | H |
| CCNA_01805 | dihydrolipoamide dehydrogenase | 10 | 0,008928571 | 6,41982116 | 0,005731983 | C |
| CCNA_00677 | LSU ribosomal protein L1P | 5 | 0,00907441 | 3,668469234 | 0,006657839 | J |
| CCNA_00343 | dihydrolipoamide succinyltransferase component (E2) of 2-oxoglutarate dehydrogenase complex | 9 | 0,00931677 | 7,336938469 | 0,007595174 | C |
| CCNA_03331 | phosphomethylpyrimidine kinase/hydroxymethylpyrimidine kinase | 6 | 0,00933126 | 3,668469234 | 0,00570524 | H |
| CCNA_03582 | 3 2 ,5 -bisphosphate nucleotidase/myo-inositol 1-(4-)phosphatase | 6 | 0,009419152 | 4,585586543 | 0,007198723 | P |
| CCNA_02030 | hypothetical protein | 3 | 0,00952381 | 1,834234617 | 0,005822967 | NA |
| CCNA_02080 | Sec-independent protein translocase protein tatC | 7 | 0,009695291 | 5,502703851 | 0,007621473 | U |
| CCNA_01395 | electron transfer flavoprotein-ubiquinone oxidoreductase | 13 | 0,009701493 | 6,41982116 | 0,004790911 | C |
| CCNA_R0188 | small non-coding RNA | 2 | 0,009756098 | 4,585586543 | 0,022368715 | NA |
| CCNA_00684 | transcriptional activator chrR | 5 | 0,009803922 | 6,41982116 | 0,012587885 | T |
| CCNA_00749 | ferrous iron transport protein B | 15 | 0,00990099 | 10,08829039 | 0,006658938 | P |
| CCNA_01178 | oxidoreductase | 4 | 0,00990099 | 3,668469234 | 0,009080369 | S |
| CCNA_00212 | N-carbamoylputrescine amidase | 7 | 0,00997151 | 3,668469234 | 0,00522574 | R |
| CCNA_00013 | protein-PII uridylyltransferase | 23 | 0,010186005 | 16,50811155 | 0,007310944 | O |
| CCNA_01965 | hypothetical protein | 1 | 0,010204082 | 8,254055777 | 0,084225059 | NA |
| CCNA_01751 | hypothetical protein | 9 | 0,010227273 | 8,254055777 | 0,009379609 | R |
| CCNA_00004 | dephospho-CoA kinase | 5 | 0,010438413 | 3,668469234 | 0,0076586 | H |
| CCNA_02224 | methylenetetrahydrofolate reductase | 8 | 0,010582011 | 3,668469234 | 0,004852473 | E |
| CCNA_01447 | homoserine dehydrogenase | 11 | 0,010669253 | 9,171173086 | 0,008895415 | E |
| CCNA_00494 | transport protein | 4 | 0,010840108 | 3,668469234 | 0,009941651 | C |
| CCNA_03527 | glutamate-cysteine ligase | 11 | 0,010858835 | 5,502703851 | 0,005432087 | NA |
| CCNA_R0147 | small non-coding RNA | 1 | 0,010869565 | 1,834234617 | 0,019937333 | NA |
| CCNA_03397 | cytosolic protein | 2 | 0,010869565 | 4,585586543 | 0,024921666 | S |
| ***CCNA_03911*** | ***hypothetical protein*** | ***1*** | 0,010989011 | 9,171173086 | 0,100782122 | NA |
| CCNA_03972 | hypothetical protein | 1 | 0,010989011 | 9,171173086 | 0,100782122 | NA |
| CCNA_03584 | histidine phosphotransferase chpT | 6 | 0,011090573 | 4,585586543 | 0,00847613 | NA |
| CCNA_03116 | hypothetical protein | 2 | 0,011111111 | 3,668469234 | 0,020380385 | NA |
| CCNA_00211 | agmatine deiminase | 9 | 0,011180124 | 7,336938469 | 0,009114209 | NA |
| CCNA_01959 | 3-dehydroquinate dehydratase | 4 | 0,011363636 | 1,834234617 | 0,005210894 | E |

| | | | | | | |
|---|---|---|---|---|---|---|
| CCNA_00931 | GTP cyclohydrolase II/3,4-dihydroxy-2-butanone-4-phosphate synthase | 11 | **0,011434511** | 11,92252501 | **0,012393477** | H |
| CCNA_00670 | O-antigen export system ATP-binding protein | 7 | **0,011627907** | 4,585586543 | **0,007617253** | GM |
| CCNA_03851 | imidazole glycerol phosphate synthase, glutamine amidotransferase subunit | 6 | **0,011695906** | 3,668469234 | **0,007151012** | E |
| *CCNA_03703* | *cytidylate kinase* | *6* | **0,011695906** | 5,502703851 | **0,010726518** | F |
| CCNA_02126 | hypothetical protein | 2 | **0,011695906** | 2,751351926 | **0,016089777** | NA |
| CCNA_01547 | methionyl-tRNA synthetase | 16 | **0,011704462** | 7,336938469 | **0,005367182** | J |
| CCNA_02635 | cell division protein ftsW | 11 | **0,011727079** | 7,336938469 | **0,007821896** | D |
| CCNA_03654 | hypothetical protein | 1 | **0,011764706** | 6,41982116 | **0,075527308** | NA |
| CCNA_03910 | hypothetical protein | 1 | **0,011764706** | 6,41982116 | **0,075527308** | NA |
| CCNA_00410 | acetyltransferase | 5 | **0,011792453** | 9,171173086 | **0,021630125** | R |
| CCNA_03842 | molybdopterin biosynthesis MoeB protein | 7 | **0,01188455** | 5,502703851 | **0,009342451** | NA |
| CCNA_00930 | riboflavin synthase subunit alpha | 6 | **0,012170385** | 3,668469234 | **0,007441114** | H |
| CCNA_00315 | 3-polyprenyl-4-hydroxybenzoate decarboxylase | 15 | **0,012264922** | 7,336938469 | **0,005999132** | H |
| CCNA_00033 | polyribonucleotide nucleotidyltransferase/polynucleotide adenylyltransferase | 21 | **0,012280702** | 9,171173086 | **0,005363259** | J |
| CCNA_02623 | cell division protein FtsZ | 15 | **0,012295082** | 6,41982116 | **0,005262148** | D |
| CCNA_02081 | Sec-independent protein translocase protein tatB | 6 | **0,012448133** | 13,75675963 | **0,028540995** | U |
| CCNA_03719 | undecaprenyl-diphosphatase (bacitracin resistance protein) | 8 | **0,0125** | 7,336938469 | **0,011463966** | V |
| CCNA_01983 | citrate synthase | 13 | **0,012695313** | 5,502703851 | **0,005373734** | C |
| CCNA_03626 | ATP phosphoribosyltransferase | 10 | **0,012987013** | 15,59099425 | **0,020248044** | E |
| CCNA_00222 | glucose-6-phosphate isomerase/glucose-6 phosphate 1-epimerase | 17 | **0,013127413** | 23,84505002 | **0,018413166** | G |
| CCNA_02247 | pantoate-beta-alanine ligase | 9 | **0,013138686** | 9,171173086 | **0,013388574** | H |
| CCNA_03305 | SSU ribosomal protein S7P | 5 | **0,013192612** | 1,834234617 | **0,004839669** | J |
| CCNA_02024 | NADH-quinone oxidoreductase chain F | 14 | **0,013270142** | 7,336938469 | **0,006954444** | C |
| CCNA_02879 | genetic exchange related protein | 1 | **0,013333333** | 0,917117309 | **0,012228231** | NA |
| ***CCNA_02816*** | ***hypothetical protein*** | ***4*** | **0,013468013** | 11,92252501 | **0,040143182** | NA |
| CCNA_03516 | protoheme IX farnesyltransferase coxE | 11 | **0,013496933** | 0,917117309 | **0,001125297** | H |
| CCNA_03734 | glutamine-dependent NAD(+) synthetase | 22 | **0,013513514** | 5,502703851 | **0,003380039** | H |
| CCNA_03314 | ferredoxin-NADP reductase | 9 | **0,013615734** | 0 | **0** | H |
| CCNA_02041 | ATP-dependent endopeptidase clp, proteolytic subunit ClpP | 7 | **0,013645224** | 1,834234617 | **0,003575506** | OU |
| CCNA_00324 | TolR protein | 5 | **0,013661202** | 10,08829039 | **0,027563635** | U |
| CCNA_00899 | bacterial peptide chain Release factor 1 (RF-1) | 11 | **0,013664596** | 0,917117309 | **0,001139276** | NA |
| CCNA_01792 | LSU ribosomal protein L32P | 2 | **0,01369863** | 0 | **0** | J |

| | | | | | | |
|---|---|---|---|---|---|---|
| CCNA_0163 7 | topoisomerase IV subunit A | 25 | 0,013713659 | 2,75135192 6 | 0,001509244 | L |
| CCNA_0261 2 | alanyl-tRNA synthetase | 29 | 0,01371807 | 3,66846923 4 | 0,001735321 | J |
| CCNA_0011 7 | glucosamine-fructose-6-phosphate aminotransferase, isomerizing | 20 | 0,013736264 | 3,66846923 4 | 0,002519553 | M |
| CCNA_0010 6 | 2-polyprenyl-6-methoxyphenol hydroxylase | 14 | 0,013820336 | 4,58558654 3 | 0,004526739 | HC |
| ***CCNA_0032 0*** | ***LSU ribosomal protein L27P*** | ***3*** | 0,013953488 | 0,91711730 9 | 0,004265662 | J |
| CCNA_0342 8 | cytosolic protein | 3 | 0,013953488 | 6,41982116 | 0,029859633 | S |
| CCNA_0200 0 | SSU ribosomal protein S2P | 9 | 0,013975155 | 2,75135192 6 | 0,004272286 | J |
| ***CCNA_0130 6*** | ***LSU ribosomal protein L3P*** | ***9*** | 0,01399689 | 0 | 0 | J |
| CCNA_0206 6 | Phytoene synthase family protein | 8 | 0,014010508 | 0,91711730 9 | 0,00160616 | I |
| CCNA_0155 0 | sulfate adenylyltransferase subunit 2 | 11 | 0,014030612 | 0 | 0 | EH |
| CCNA_0337 8 | myo-inositol-1(or 4)-monophosphatase | 9 | 0,0140625 | 0,91711730 9 | 0,001432996 | NA |
| CCNA_0168 7 | 23S rRNA Um2552 2 -O-methyltransferase | 8 | 0,014084507 | 0 | 0 | J |
| CCNA_0372 2 | glutamate synthase (NADPH) large chain | 51 | 0,014088398 | 12,8396423 2 | 0,003546863 | E |
| CCNA_0080 6 | inner membrane protein translocase component yidC | 21 | 0,014218009 | 3,66846923 4 | 0,00248373 | U |
| CCNA_0037 9 | thiol:disulfide interchange protein dsbA | 7 | 0,014227642 | 0,91711730 9 | 0,00186406 | O |
| CCNA_0141 7 | 5-aminolevulinic acid synthase | 14 | 0,014285714 | 3,66846923 4 | 0,003743336 | H |
| CCNA_0004 8 | S-adenosylmethionine synthetase | 14 | 0,014300306 | 1,83423461 7 | 0,00187358 | H |
| CCNA_0132 3 | SSU ribosomal protein S5P | 7 | 0,01443299 | 0 | 0 | J |
| CCNA_R019 2 | small non-coding RNA | 1 | 0,014492754 | 0 | 0 | NA |
| CCNA_0195 0 | bacterial peptide chain Release factor 2 (RF-2) | 13 | 0,014590348 | 2,75135192 6 | 0,003087937 | J |
| CCNA_0199 4 | 1-deoxy-D-xylulose 5-phosphate reductoisomerase | 14 | 0,01459854 | 0,91711730 9 | 0,000956327 | I |
| CCNA_0282 2 | transcriptional regulator, MerR family | 10 | 0,014619883 | 2,75135192 6 | 0,004022444 | K |
| CCNA_0238 8 | ribose 5-phosphate isomerase | 8 | 0,014625229 | 0,91711730 9 | 0,001676631 | G |
| CCNA_0196 0 | biotin carboxyl carrier protein of acetyl-CoA | 6 | 0,014634146 | 1,83423461 7 | 0,004473743 | I |
| *CCNA_03873* | *hypothetical protein* | *3* | 0,014634146 | 12,8396423 2 | 0,062632402 | NA |
| CCNA_0179 5 | enolase | 15 | 0,014648438 | 0,91711730 9 | 0,000895622 | G |
| ***CCNA_R009 0*** | ***tRNA Leu*** | ***1*** | 0,014705882 | 0 | 0 | NA |
| CCNA_R005 3 | tRNA Leu | 1 | 0,014705882 | 0 | 0 | NA |
| CCNA_R018 0 | small non-coding RNA | 1 | 0,014705882 | 1,83423461 7 | 0,026974038 | NA |
| CCNA_0225 6 | Mg2+ transporter mgtE | 17 | 0,014808362 | 0 | 0 | P |
| CCNA_0093 9 | transcriptional regulator, MarR family | 6 | 0,014888337 | 40,3531615 8 | 0,100131915 | K |
| CCNA_R002 5 | tRNA Gly | 4 | 0,014925373 | 11,0054077 | 0,041064954 | NA |
| CCNA_0200 3 | DNA polymerase III alpha subunit | 41 | 0,014936248 | 6,41982116 | 0,002338733 | L |
| CCNA_0202 9 | NADH-quinone oxidoreductase chain D | 15 | 0,015 | 2,75135192 6 | 0,002751352 | C |

| | | | | | | |
|---|---|---|---|---|---|---|
| *CCNA_03866* | *hypothetical protein* | *6* | 0,015 | 1,834234617 | 0,004585587 | S |
| CCNA_02008 | prolyl-tRNA synthetase | 16 | 0,015023474 | 3,668469234 | 0,003444572 | J |
| CCNA_03798 | tRNA 2-methylthioadenosine synthase-like protein | 16 | 0,015023474 | 3,668469234 | 0,003444572 | J |
| CCNA_00130 | Kup system potassium uptake protein | 24 | 0,015028178 | 96,2973174 | 0,060298884 | P |
| *CCNA_00536* | *DNA-directed RNA polymerase subunit beta* | *49* | 0,01504914 | 13,75675963 | 0,004225049 | K |
| CCNA_03780 | hypothetical protein | 9 | 0,01512605 | 0,917117309 | 0,001541374 | S |
| CCNA_03510 | threonine synthase | 17 | 0,015219338 | 3,668469234 | 0,003284216 | E |
| CCNA_03469 | arginyl-tRNA synthetase | 22 | 0,015256588 | 2,751351926 | 0,001908011 | J |
| CCNA_03544 | nicotinate-nucleotide adenylyltransferase | 8 | 0,015384615 | 1,834234617 | 0,003527374 | H |
| CCNA_01997 | ribosome Recycling factor (RRF) | 7 | 0,015486726 | 0 | 0 | J |
| CCNA_03562 | ATP synthase subunit alpha | 19 | 0,015497553 | 2,751351926 | 0,00224417 | C |
| CCNA_02306 | histidinol-phosphate aminotransferase | 14 | 0,015607581 | 3,668469234 | 0,004089709 | E |
| CCNA_02007 | lipoprotein releasing system transmembrane protein lolE | 16 | 0,015625 | 4,585586543 | 0,004478112 | M |
| CCNA_02182 | phosphoserine phosphatase | 11 | 0,015669516 | 0 | 0 | NA |
| CCNA_03164 | protein translocase subunit secA | 35 | 0,0157871 | 9,171173086 | 0,004136749 | U |
| CCNA_02462 | hypothetical protein | 4 | 0,015810277 | 9,171173086 | 0,036249696 | S |
| CCNA_01441 | SSU ribosomal protein S9P | 6 | 0,015831135 | 0 | 0 | J |
| *CCNA_03607* | *ribonucleoside-diphosphate reductase subunit alpha* | *24* | 0,01592568 | 6,41982116 | 0,004260001 | F |
| CCNA_00364 | deoxyhypusine synthase | 14 | 0,015981735 | 3,668469234 | 0,00418775 | O |
| ***CCNA_01330*** | ***DNA-directed RNA polymerase subunit alpha*** | ***13*** | 0,016009852 | 0,917117309 | 0,001129455 | K |
| *CCNA_01669* | *sulfate transport system permease protein cysW* | *11* | 0,016058394 | 0 | 0 | P |
| CCNA_02337 | nonspecific lipid-transfer protein | 4 | 0,016260163 | 0 | 0 | I |
| CCNA_01132 | sensory transduction histidine kinase/receiver protein CckA | 27 | 0,016274864 | 5,502703851 | 0,00331688 | T |
| CCNA_00894 | 1-hydroxy-2-methyl-2-(E)-butenyl 4-diphosphate synthase | 15 | 0,016286645 | 3,668469234 | 0,003983137 | I |
| CCNA_00265 | AMP nucleosidase | 19 | 0,016323024 | 59,61262506 | 0,051213595 | F |
| CCNA_00168 | putative capsule polysaccharide export protein | 8 | 0,016359918 | 63,28109429 | 0,129409191 | M |
| CCNA_00293 | phosphate transport system permease protein pstA | 17 | 0,016425121 | 2,751351926 | 0,002658311 | P |
| CCNA_02759 | methionine aminopeptidase | 11 | 0,016566265 | 0,917117309 | 0,001381201 | J |
| CCNA_03142 | RNA polymerase sigma factor rpoD | 26 | 0,016571064 | 6,41982116 | 0,004091664 | K |
| CCNA_03334 | cell division protein ftsH | 25 | 0,01662234 | 3,668469234 | 0,002439142 | D |
| CCNA_00372 | ATP synthase C chain | 3 | 0,016666667 | 0 | 0 | C |
| CCNA_01562 | 4-hydroxy-2-oxoglutarate aldolase/2-dehydro-3-deoxyphosphogluconate aldolase | 9 | 0,016666667 | 0,917117309 | 0,001698365 | G |

| | | | | | | |
|---|---|---|---|---|---|---|
| CCNA_00020 | hypothetical protein | 3 | 0,016666667 | 11,0054077 | 0,061141154 | NA |
| CCNA_R0172 | small non-coding RNA | 1 | 0,016666667 | 6,41982116 | 0,106997019 | NA |
| CCNA_01815 | nitrogen assimilation regulatory protein | 19 | 0,016710642 | 3,668469234 | 0,003226446 | T |
| CCNA_02016 | NADH-quinone oxidoreductase chain M | 20 | 0,016835017 | 5,502703851 | 0,004631906 | C |
| CCNA_00308 | hypothetical protein | 9 | 0,016885553 | 1,834234617 | 0,003441341 | T |
| CCNA_01998 | uridylate kinase | 10 | 0,016891892 | 0 | 0 | F |
| CCNA_R0089 | tRNA Ala | 1 | 0,016949153 | 0 | 0 | NA |
| CCNA_R0058 | tRNA Phe | 1 | 0,016949153 | 0 | 0 | NA |
| CCNA_R0022 | tRNA Gln | 1 | 0,016949153 | 0 | 0 | NA |
| CCNA_R0055 | tRNA Met | 1 | 0,016949153 | 3,668469234 | 0,062177445 | NA |
| CCNA_01614 | hypothetical protein mreD | 7 | 0,01703163 | 0 | 0 | NA |
| *CCNA_01670* | *sulfate transport ATP-binding protein cysA* | *14* | 0,017177914 | 3,668469234 | 0,004501189 | P |
| CCNA_01760 | organic solvent tolerance protein | 33 | 0,017196456 | 1,834234617 | 0,000955828 | M |
| CCNA_00370 | ATP synthase B chain | 7 | 0,017199017 | 0 | 0 | C |
| CCNA_02420 | TonB accessory protein exbD | 6 | 0,017241379 | 0,917117309 | 0,002635395 | U |
| CCNA_00564 | sensory transduction protein kinase CenK | 21 | 0,017427386 | 2,751351926 | 0,00228328 | T |
| CCNA_01753 | guanylate kinase | 9 | 0,01754386 | 1,834234617 | 0,003575506 | F |
| CCNA_00491 | GTP cyclohydrolase I | 9 | 0,017647059 | 0,917117309 | 0,001798269 | H |
| CCNA_01986 | lipid-A-disaccharide synthase | 17 | 0,017782427 | 0,917117309 | 0,000959328 | M |
| CCNA_01450 | single-stranded-DNA-specific exonuclease recJ | 26 | 0,017857143 | 2,751351926 | 0,001889665 | H |
| CCNA_02280 | 2-polyprenyl-3-methyl-6-methoxy-1,4-benzoquinol hydroxylase | 8 | 0,017857143 | 0,917117309 | 0,002047137 | L |
| CCNA_00667 | capsular polysaccharide biosynthesis protein | 17 | 0,017932489 | 48,60721735 | 0,051273436 | G |
| CCNA_02194 | fumarate hydratase | 20 | 0,017969452 | 0 | 0 | C |
| CCNA_02036 | DNA-binding protein HU | 4 | 0,018018018 | 16,50811155 | 0,074360863 | L |
| CCNA_02198 | trans-isoprenyl diphosphate synthase | 14 | 0,018087855 | 0,917117309 | 0,001184906 | NA |
| CCNA_00071 | uroporphyrinogen-III synthase | 10 | 0,018248175 | 1,834234617 | 0,003347143 | H |
| CCNA_02019 | NADH-quinone oxidoreductase chain J | 9 | 0,018255578 | 0,917117309 | 0,001860279 | C |
| CCNA_02518 | aspartyl/glutamyl-tRNA(Asn/Gln) amidotransferase subunit B | 22 | 0,018394649 | 3,668469234 | 0,003067282 | NA |
| CCNA_R0162 | small non-coding RNA | 1 | 0,018518519 | 0 | 0 | E |
| CCNA_00559 | homoserine O-acetyltransferase | 17 | 0,018518519 | 0,917117309 | 0,000999038 | I |
| CCNA_01746 | malonyl-CoA-(acyl-carrier-protein) transacylase | 14 | 0,018518519 | 1,834234617 | 0,002426236 | NA |
| CCNA_01963 | hypothetical protein | 5 | 0,018518519 | 11,92252501 | 0,0441575 | NA |
| CCNA_01007 | hypothetical protein | 4 | 0,018604651 | 14,67387694 | 0,06825059 | NA |
| CCNA_02047 | glutamine synthetase | 21 | 0,01863354 | 2,751351926 | 0,002441306 | E |

| | | | | | | |
|---|---|---|---|---|---|---|
| CCNA_00238 | two-component sensor histidine kinase chvG | 24 | 0,018691589 | 41,27027889 | 0,032141962 | NA |
| CCNA_R0088 | Minimal medium sRNA | 2 | 0,018691589 | 11,0054077 | 0,102854278 | T |
| *CCNA_00443* | *chemotaxis protein cheW* | *7* | 0,018766756 | 23,84505002 | 0,063927748 | NT |
| CCNA_02028 | NTF2 enzyme family protein | 5 | 0,018796992 | 0,917117309 | 0,003447809 | S |
| CCNA_00257 | adenosylhomocysteinase | 21 | 0,018867925 | 3,668469234 | 0,003296019 | H |
| CCNA_01101 | bacterial protein translation initiation factor 3 (IF-3) | 8 | 0,019184652 | 1,834234617 | 0,004398644 | J |
| CCNA_03854 | phosphoribosyl-ATP pyrophosphatase | 5 | 0,019305019 | 0 | 0 | E |
| CCNA_01459 | hypothetical protein | 4 | 0,019512195 | 14,67387694 | 0,071579887 | NA |
| CCNA_02070 | protein translocase subunit secD | 25 | 0,019561815 | 5,502703851 | 0,004305715 | U |
| CCNA_03606 | hypothetical protein | 2 | 0,01980198 | 0 | 0 | NA |
| CCNA_03534 | 4-hydroxybenzoate polyprenyltransferase | 15 | 0,019815059 | 2,751351926 | 0,003634547 | NA |
| CCNA_01317 | LSU ribosomal protein L24P | 5 | 0,01984127 | 0 | 0 | J |
| CCNA_00292 | phosphate transport system permease protein pstC | 23 | 0,02005231 | 2,751351926 | 0,002398738 | P |
| CCNA_03937 | hypothetical protein | 3 | 0,020134228 | 20,17658079 | 0,135413294 | NA |
| *CCNA_03736* | *YGGT family protein* | *6* | 0,02020202 | 17,42522886 | 0,058670804 | NA |
| CCNA_R0106 | small non-coding RNA | 2 | 0,02020202 | 7,336938469 | 0,07411049 | S |
| CCNA_01159 | transporter, major facilitator superfamily | 22 | 0,020313943 | 1,834234617 | 0,001693661 | G |
| CCNA_01458 | hypothetical protein | 5 | 0,020325203 | 23,84505002 | 0,096931098 | NA |
| CCNA_01652 | phosphopantetheine adenylyltransferase | 8 | 0,020356234 | 0 | 0 | H |
| CCNA_03393 | phosphoribosylaminoimidazole carboxylase carboxyltransferase subunit | 8 | 0,020356234 | 0 | 0 | NA |
| CCNA_R0183 | small non-coding RNA | 4 | 0,020408163 | 4,585586543 | 0,02339585 | NA |
| CCNA_03550 | hypothetical protein | 2 | 0,020408163 | 8,254055777 | 0,084225059 | NA |
| CCNA_03197 | alpha/beta hydrolase family protein | 12 | 0,020408163 | 54,10992121 | 0,092023676 | R |
| *CCNA_02185* | *acetolactate synthase large subunit* | *30* | 0,020761246 | 2,751351926 | 0,00190405 | EH |
| ***CCNA_01308*** | ***LSU ribosomal protein L23P*** | ***5*** | 0,021186441 | 0 | 0 | J |
| CCNA_01120 | hypothetical protein | 5 | 0,021276596 | 14,67387694 | 0,06244203 | NA |
| CCNA_00721 | chaperonin GroEL | 28 | 0,021292776 | 0,917117309 | 0,000697428 | O |
| CCNA_R0013 | small non-coding RNA | 2 | 0,02173913 | 9,171173086 | 0,099686664 | NA |
| CCNA_01432 | 3-oxoacyl-(acyl-carrier-protein) synthase III | 17 | 0,021766965 | 1,834234617 | 0,002348572 | I |
| CCNA_02490 | 3-ketoacyl-CoA thiolase | 21 | 0,02189781 | 22,92793271 | 0,023908168 | I |
| *CCNA_02818* | *hypothetical protein* | *2* | 0,021978022 | 0 | 0 | NA |
| CCNA_01842 | hypothetical protein | 4 | 0,022099448 | 8,254055777 | 0,045602518 | NA |
| CCNA_03852 | 1-(5-phosphoribosyl)-5-((5-phosphoribosylamino)methylideneamino) imidazole-4-carboxamide isomerase | 13 | 0,022108844 | 0,917117309 | 0,001559723 | E |

| | | | | | | |
|---|---|---|---|---|---|---|
| *CCNA_R0032* | *cell cycle sRNA* | *1* | **0,022222222** | 0 | **0** | NA |
| CCNA_00231 | tRNA-specific adenosine deaminase | 8 | **0,02247191** | 0,917117309 | **0,002576172** | FJ |
| *CCNA_03427* | *hypothetical protein* | *6* | **0,02247191** | 19,25946348 | **0,072132822** | NA |
| CCNA_03907 | conserved hypothetical protein | 6 | **0,02247191** | 19,25946348 | **0,072132822** | NA |
| CCNA_01740 | SSU ribosomal protein S18P | 5 | **0,022522523** | 0 | **0** | J |
| CCNA_02107 | hypothetical protein | 4 | **0,02259887** | 12,83964232 | **0,072540352** | NA |
| CCNA_03955 | hypothetical protein | 1 | **0,022727273** | 0 | **0** | NA |
| CCNA_02039 | ATP-dependent endopeptidase clp, ATP-binding subunit ClpX | 23 | **0,022772277** | 1,834234617 | **0,001816074** | O |
| CCNA_01709 | hypothetical protein | 6 | **0,022813688** | 11,92252501 | **0,045332795** | NA |
| CCNA_00226 | hypothetical protein | 5 | **0,02283105** | 8,254055777 | **0,037689752** | S |
| CCNA_03182 | hypothetical protein | 7 | **0,023569024** | 21,0936981 | **0,071022553** | NA |
| CCNA_02025 | hypothetical protein | 5 | **0,023584906** | 4,585586543 | **0,021630125** | NA |
| CCNA_00389 | aminomethyltransferase family protein | 16 | **0,023633678** | 0,917117309 | **0,001354678** | R |
| CCNA_03631 | hypothetical protein | 2 | **0,023809524** | 6,41982116 | **0,076426442** | HE |
| CCNA_03321 | hypothetical protein | 7 | **0,023809524** | 29,34775387 | **0,099822292** | NA |
| CCNA_01903 | cytosolic protein | 16 | **0,023845007** | 36,68469234 | **0,054671673** | S |
| CCNA_02279 | disulfide bond formation protein B | 12 | **0,023856859** | 0,917117309 | **0,001823295** | O |
| CCNA_00023 | putative periplasmic protein | 9 | **0,02393617** | 25,67928464 | **0,06829597** | R |
| *CCNA_03871* | *glucose inhibited division protein A* | *37* | **0,024057217** | 5,502703851 | **0,003577831** | D |
| CCNA_02078 | seryl-tRNA synthetase | 28 | **0,02417962** | 2,751351926 | **0,002375952** | J |
| CCNA_01421 | 6,7-dimethyl-8-ribityllumazine synthase | 9 | **0,024193548** | 0 | **0** | H |
| CCNA_03657 | tryptophan synthase subunit alpha | 16 | **0,024205749** | 0 | **0** | E |
| CCNA_02321 | O-succinylhomoserine sulfhydrylase | 23 | **0,024261603** | 2,751351926 | **0,00290227** | E |
| *CCNA_01331* | *hypothetical protein* | *4* | **0,024390244** | 4,585586543 | **0,027960894** | NA |
| *CCNA_00860* | *holdfast inhibitor HfiA* | *4* | **0,024390244** | 7,336938469 | **0,04473743** | NA |
| CCNA_01691 | 16S rRNA m(5)C 967 methyltransferase | 25 | **0,024485798** | 2,751351926 | **0,002694762** | J |
| CCNA_01969 | aspartyl-tRNA synthetase | 36 | **0,024489796** | 6,41982116 | **0,004367225** | J |
| CCNA_01269 | GcrB protein | 10 | **0,024813896** | 26,59640195 | **0,065996035** | S |
| CCNA_02804 | hypothetical protein | 4 | **0,025** | 7,336938469 | **0,045855865** | NA |
| *CCNA_03545* | *iojap protein family* | *11* | **0,025171625** | 10,08829039 | **0,023085333** | S |
| CCNA_01657 | hypothetical protein | 13 | **0,025193798** | 21,0936981 | **0,04087926** | NA |
| CCNA_00250 | poly(3-hydroxyalkanoate) depolymerase | 26 | **0,025316456** | 88,96037893 | **0,086621596** | I |
| CCNA_00561 | conserved cytosolic protein | 7 | **0,025362319** | 19,25946348 | **0,069780665** | S |
| CCNA_02620 | hypothetical protein | 6 | **0,025862069** | 27,51351926 | **0,118592755** | NA |
| *CCNA_01060* | *type I protein secretion ATP-binding protein RsaD* | *36* | **0,025936599** | 36,68469234 | **0,026429894** | R |

| | | | | | |
|---|---|---|---|---|---|
| *CCNA_R0116* | *small non-coding RNA* | *5* | 0,02617801 | 17,42522886 | 0,091231565 | NA |
| CCNA_02175 | hypothetical protein | 11 | 0,026190476 | 10,08829039 | 0,024019739 | NA |
| CCNA_01497 | ADP-L-glycero-D-manno-heptose-6-epimerase | 21 | 0,026217228 | 49,52433466 | 0,061828133 | MG |
| CCNA_R0018 | Stat phase sRNA | 5 | 0,026315789 | 7,336938469 | 0,038615466 | NA |
| CCNA_01292 | hypothetical protein | 9 | 0,026392962 | 11,92252501 | 0,034963416 | NA |
| CCNA_03997 | amelogenin/CpxP-related protein | 14 | 0,026415094 | 46,77298274 | 0,088250911 | NA |
| CCNA_02958 | hypothetical protein | 2 | 0,026666667 | 1,834234617 | 0,024456462 | NA |
| CCNA_03745 | formyltetrahydrofolate deformylase | 18 | 0,026706231 | 21,0936981 | 0,031296288 | F |
| CCNA_03484 | hypothetical protein | 10 | 0,02688172 | 36,68469234 | 0,098614764 | OU |
| *CCNA_03462* | *glycine dehydrogenase (decarboxylating)* | *34* | 0,026898734 | 89,87749624 | 0,071105614 | E |
| *CCNA_03325* | *hypothetical protein* | *7* | 0,026923077 | 15,59099425 | 0,059965362 | L |
| *CCNA_01405* | *cytosolic protein* | *7* | 0,026923077 | 22,92793271 | 0,088184357 | T |
| ***CCNA_03306*** | ***SSU ribosomal protein S12P*** | ***8*** | 0,026936027 | 0 | 0 | NA |
| CCNA_03069 | hypothetical protein | 8 | 0,026936027 | 18,34234617 | 0,061758741 | NA |
| CCNA_03968 | hypothetical protein | 3 | 0,027027027 | 3,668469234 | 0,033049272 | NA |
| CCNA_01001 | ribokinase | 20 | 0,027434842 | 44,93874812 | 0,061644373 | K |
| CCNA_01593 | Xaa-pro aminopeptidase | 40 | 0,027605245 | 73,36938469 | 0,050634496 | E |
| CCNA_03998 | GT1/WbuB-like N-acetyl-L-fucosamine transferase | 28 | 0,027613412 | 110,9711943 | 0,109439048 | NA |
| CCNA_01516 | NAD(P)H-dependent quinone reductase | 14 | 0,027667984 | 24,76216733 | 0,04893709 | C |
| CCNA_01858 | transcriptional regulator, AraC family | 20 | 0,027700831 | 53,1928039 | 0,073674244 | K |
| CCNA_01006 | flagellar hook-basal body complex protein FliE | 7 | 0,02811245 | 19,25946348 | 0,077347243 | NU |
| CCNA_02004 | transcriptional regulator, SidA | 2 | 0,028169014 | 1,834234617 | 0,02583429 | NA |
| *CCNA_03608* | *hypothetical protein* | *8* | 0,028268551 | 23,84505002 | 0,084258127 | NA |
| CCNA_02863 | Spr-family cell wall-associated hydrolase | 11 | 0,028277635 | 29,34775387 | 0,075444097 | M |
| ***CCNA_00044*** | ***cytosolic protein*** | ***13*** | 0,02832244 | 11,92252501 | 0,025975 | S |
| CCNA_01827 | adenosylcobyric acid synthase (glutamine-hydrolysing) | 33 | 0,028350515 | 48,60721735 | 0,041758778 | H |
| CCNA_03785 | hypothetical protein | 19 | 0,028443114 | 59,61262506 | 0,089240457 | NA |
| CCNA_02844 | antitoxin protein parD-3 | 5 | 0,028735632 | 4,585586543 | 0,026353946 | K |
| *CCNA_00049* | *HTH transcriptional regulator* | *10* | 0,028735632 | 36,68469234 | 0,105415783 | NA |
| *CCNA_00273* | *peptide deformylase* | *12* | 0,028776978 | 11,0054077 | 0,026391865 | J |
| CCNA_02282 | hypothetical protein | 12 | 0,028776978 | 19,25946348 | 0,046185764 | NA |
| CCNA_02425 | non-essential pilus assembly protein | 4 | 0,028776978 | 14,67387694 | 0,10556746 | S |
| ***CCNA_03558*** | ***ATP synthase subunit epsilon*** | ***6*** | 0,028846154 | 0 | 0 | C |
| CCNA_00769 | transcriptional regulator, TetR family | 15 | 0,02901354 | 36,68469234 | 0,070956852 | K |

| | | | | | | |
|---|---|---|---|---|---|---|
| CCNA_03836 | 3-hydroxydecanoyl-(acyl-carrier-protein) dehydratase | 12 | 0,029268293 | 0,917117309 | 0,002236871 | I |
| CCNA_R0052 | tRNA Leu | 2 | 0,029411765 | 0 | 0 | NA |
| ***CCNA_R0048*** | ***tRNA Leu*** | ***2*** | 0,029411765 | 1,834234617 | 0,026974038 | NA |
| CCNA_R0093 | Minimal medium sRNA CrfA | 3 | 0,029411765 | 7,336938469 | 0,071930769 | NA |
| ***CCNA_01665*** | ***potassium-transporting ATPase C chain kdpC*** | ***14*** | 0,029473684 | 1,834234617 | 0,003861547 | E |
| CCNA_00300 | glyoxalase superfamily protein | 14 | 0,029473684 | 44,02163081 | 0,092677117 | P |
| CCNA_01654 | peptidyl-prolyl cis-trans isomerase | 20 | 0,029498525 | 32,0991058 | 0,047343814 | O |
| CCNA_01326 | protein translocase subunit secY | 32 | 0,029547553 | 24,76216733 | 0,02286442 | U |
| CCNA_00306 | hypothetical protein | 17 | 0,029772329 | 12,83964232 | 0,022486239 | S |
| CCNA_01824 | hypothetical protein | 8 | 0,029962547 | 8,254055777 | 0,030914067 | NA |
| CCNA_R0140 | small non-coding RNA | 2 | 0,03030303 | 0 | 0 | NA |
| CCNA_00177 | type II secretion pathway protein I | 9 | 0,03030303 | 23,84505002 | 0,080286364 | NA |
| CCNA_R0096 | small non-coding RNA | 2 | 0,03030303 | 5,502703851 | 0,083374301 | NU |
| CCNA_00316 | glutamate 5-kinase | 28 | 0,030871003 | 0 | 0 | E |
| CCNA_01201 | hypothetical protein | 13 | 0,030952381 | 48,60721735 | 0,11573147 | NA |
| CCNA_02509 | glycosyltransferase hfsG | 23 | 0,030955585 | 52,27568659 | 0,070357586 | R |
| CCNA_00471 | GDP-L-fucose synthase | 24 | 0,031007752 | 56,86127313 | 0,073464177 | MG |
| CCNA_00507 | cytochrome c1 | 21 | 0,031019202 | 19,25946348 | 0,028448247 | C |
| CCNA_01383 | hypothetical protein | 9 | 0,031034483 | 9,171173086 | 0,031624735 | S |
| CCNA_01582 | hypothetical protein | 9 | 0,031034483 | 13,75675963 | 0,047437102 | S |
| CCNA_01788 | hypothetical protein | 9 | 0,031358885 | 6,41982116 | 0,022368715 | NA |
| CCNA_01142 | hypothetical protein | 12 | 0,031578947 | 23,84505002 | 0,062750132 | NA |
| CCNA_00243 | hypothetical protein | 12 | 0,031578947 | 23,84505002 | 0,062750132 | NA |
| CCNA_02380 | aspartate 1-decarboxylase | 9 | 0,031690141 | 16,50811155 | 0,058127153 | H |
| CCNA_00893 | transcriptional regulatory protein | 27 | 0,031690141 | 49,52433466 | 0,058127153 | S |
| CCNA_00666 | response regulator receiver domain protein | 9 | 0,03180212 | 22,92793271 | 0,08101743 | T |
| *CCNA_00161* | *hypothetical protein* | *16* | 0,032 | 44,02163081 | 0,088043262 | NA |
| CCNA_01952 | N-acetylmuramoyl-L-alanine amidase AmiC | 29 | 0,032329989 | 44,02163081 | 0,049076511 | NA |
| CCNA_01020 | transcriptional regulator, LacI family | 27 | 0,032335329 | 54,10992121 | 0,064802301 | K |
| CCNA_00124 | hypothetical protein | 17 | 0,032442748 | 31,18198849 | 0,059507612 | NA |
| CCNA_00295 | phosphate transport system protein phoU | 18 | 0,032490975 | 0,917117309 | 0,001655446 | P |
| CCNA_02305 | chorismate mutase-family protein | 22 | 0,032496307 | 1,834234617 | 0,002709357 | E |
| ***CCNA_00628*** | ***chemotaxis protein cheY*** | 10 | 0,03257329 | 17,42522886 | 0,056759703 | T |
| CCNA_01802 | hypothetical protein | 3 | 0,032608696 | 11,92252501 | 0,129592663 | NA |

| | | | | | | |
|---|---|---|---|---|---|---|
| CCNA_01752 | protein yicC | 23 | 0,032624113 | 15,59099425 | 0,022114885 | S |
| CCNA_03287 | transcriptional regulatory protein | 8 | 0,032653061 | 22,01081541 | 0,089840063 | N |
| CCNA_02721 | peptidase, M16 family | 78 | 0,032704403 | 159,5784117 | 0,066909187 | R |
| CCNA_03723 | PaaI thioesterase family protein | 12 | 0,032786885 | 30,26487118 | 0,082690905 | Q |
| CCNA_01501 | protein kinase C-like superfamily protein | 10 | 0,032894737 | 19,25946348 | 0,063353498 | NA |
| *CCNA_00468* | *hypothetical protein* | *3* | 0,032967033 | 3,668469234 | 0,040312849 | NA |
| CCNA_01779 | hypothetical protein | 7 | 0,033018868 | 15,59099425 | 0,073542426 | NA |
| CCNA_01797 | septum formation initiator | 8 | 0,033057851 | 5,502703851 | 0,022738446 | D |
| CCNA_01677 | carboxypeptidase G2 precursor | 33 | 0,03313253 | 65,11532891 | 0,065376836 | E |
| CCNA_03557 | hypothetical protein | 20 | 0,033167496 | 41,27027889 | 0,06844159 | NA |
| CCNA_02375 | hypothetical protein | 7 | 0,033175355 | 20,17658079 | 0,095623606 | NA |
| CCNA_00422 | 3-oxoacyl-(acyl-carrier protein) reductase | 20 | 0,033222591 | 35,76757503 | 0,059414576 | IQR |
| CCNA_R0045 | tRNA Asp | 2 | 0,033333333 | 0 | 0 | NA |
| *CCNA_R0015* | *tRNA-Val* | *2* | 0,033333333 | 0 | 0 | NA |
| CCNA_R0035 | tRA Pro | 2 | 0,033333333 | 0 | 0 | NA |
| ***CCNA_R0008*** | ***tRNA-Arg*** | ***2*** | 0,033333333 | 1,834234617 | 0,030570577 | NA |
| CCNA_02465 | UDP-glucose 6-dehydrogenase | 35 | 0,033492823 | 40,35316158 | 0,038615466 | M |
| ***CCNA_R0028*** | ***tRNA Lys*** | ***2*** | 0,033898305 | 0 | 0 | E |
| CCNA_03750 | threonine dehydratase | 34 | 0,033898305 | 3,668469234 | 0,003657497 | NA |
| CCNA_01863 | outer membrane efflux protein | 39 | 0,033972125 | 69,70091545 | 0,060715083 | MU |
| CCNA_00015 | molybdenum cofactor biosynthesis protein B | 15 | 0,034013605 | 49,52433466 | 0,112300079 | H |
| CCNA_02948 | hypothetical protein | 10 | 0,034129693 | 32,0991058 | 0,109553262 | NA |
| CCNA_00217 | thiol:disulfide interchange protein DsbD | 56 | 0,034398034 | 38,51892696 | 0,023660275 | NA |
| CCNA_02719 | hypothetical protein | 17 | 0,034552846 | 38,51892696 | 0,078290502 | NA |
| CCNA_02646 | cell division protein mraZ | 13 | 0,034574468 | 24,76216733 | 0,065856828 | S |
| CCNA_02183 | tRNA delta(2)-isopentenylpyrophosphate transferase | 25 | 0,03511236 | 0 | 0 | J |
| CCNA_00148 | hypothetical protein | 15 | 0,035128806 | 53,1928039 | 0,124573311 | NA |
| CCNA_01506 | cystine transport ATP-binding protein | 21 | 0,035472973 | 49,52433466 | 0,083655971 | E |
| CCNA_03950 | hypothetical protein | 3 | 0,035714286 | 9,171173086 | 0,109180632 | NA |
| CCNA_03987 | hypothetical protein | 3 | 0,035714286 | 15,59099425 | 0,185607074 | NA |
| CCNA_01338 | glyoxalase family protein | 13 | 0,035812672 | 33,93334042 | 0,093480277 | S |
| CCNA_01875 | acyl-CoA dehydrogenase, short-chain specific | 33 | 0,035986914 | 61,44685967 | 0,067008571 | I |
| CCNA_00463 | hypothetical protein | 8 | 0,036036036 | 24,76216733 | 0,111541294 | NA |
| CCNA_01351 | two-component response regulator protein | 21 | 0,036082474 | 45,85586543 | 0,078790147 | TK |

| | | | | | | |
|---|---|---|---|---|---|---|
| CCNA_00634 | chemotaxis protein methyltransferase | 24 | 0,036144578 | 41,27027889 | 0,062154034 | NT |
| CCNA_01154 | hypothetical protein | 20 | 0,036297641 | 32,0991058 | 0,05825609 | S |
| CCNA_02711 | holdfast attachment protein hfaA | 13 | 0,036619718 | 36,68469234 | 0,103337162 | NA |
| CCNA_01861 | hypothetical protein | 35 | 0,036649215 | 51,35856928 | 0,053778607 | NA |
| *CCNA_02544* | *hypothetical protein* | *8* | 0,036697248 | 13,75675963 | 0,063104402 | NA |
| CCNA_03844 | ATP-dependent endopeptidase hsl ATP-binding subunit hslU | 38 | 0,036750484 | 80,70632315 | 0,078052537 | O |
| CCNA_02132 | transcriptional regulator, LacI family | 30 | 0,036809816 | 61,44685967 | 0,07539492 | K |
| CCNA_01487 | transcriptional regulator, GntR family | 22 | 0,036912752 | 32,0991058 | 0,05385756 | K |
| CCNA_01618 | uracil-DNA glycosylase | 20 | 0,036968577 | 19,25946348 | 0,035599748 | L |
| CCNA_02311 | nan | 10 | 0,037037037 | 20,17658079 | 0,074728077 | NA |
| CCNA_01564 | hypothetical protein | 7 | 0,037234043 | 6,41982116 | 0,034147985 | NA |
| CCNA_03543 | gamma-glutamyl phosphate reductase | 38 | 0,037254902 | 3,668469234 | 0,003596538 | E |
| CCNA_02195 | acetyltransferase | 13 | 0,037356322 | 22,01081541 | 0,06324947 | NA |
| CCNA_R0079 | Minimal medium sRNA | 4 | 0,037383178 | 7,336938469 | 0,068569518 | NA |
| CCNA_00999 | MopJ | 16 | 0,037470726 | 29,34775387 | 0,068730103 | S |
| CCNA_03556 | quaternary ammonium compound-resistance protein | 10 | 0,037593985 | 15,59099425 | 0,05861276 | IQ |
| CCNA_03291 | hypothetical protein | 30 | 0,037593985 | 57,77839044 | 0,072403998 | P |
| CCNA_03971 | hypothetical protein | 2 | 0,037735849 | 1,834234617 | 0,0346082 | NA |
| CCNA_03732 | mannose-6-phosphate isomerase | 32 | 0,037825059 | 42,18739619 | 0,049866899 | NA |
| CCNA_03237 | hypothetical protein | 5 | 0,037878788 | 8,254055777 | 0,062530726 | NA |
| CCNA_03916 | hypothetical protein | 8 | 0,037914692 | 13,75675963 | 0,065197913 | NA |
| CCNA_02892 | hypothetical protein | 9 | 0,038135593 | 11,92252501 | 0,050519174 | NA |
| CCNA_00289 | hypothetical protein | 9 | 0,038135593 | 15,59099425 | 0,066063535 | NA |
| CCNA_00704 | hypothetical protein | 6 | 0,038216561 | 22,01081541 | 0,140196276 | NA |
| CCNA_00693 | heat shock protein 15 | 9 | 0,038297872 | 17,42522886 | 0,07414991 | J |
| CCNA_03145 | hypothetical protein | 11 | 0,038327526 | 23,84505002 | 0,083083798 | NA |
| CCNA_01190 | hypothetical protein | 47 | 0,03843009 | 99,96578664 | 0,081738174 | NA |
| CCNA_00016 | molybdopterin converting factor, large subunit | 14 | 0,038674033 | 22,92793271 | 0,063336831 | H |
| CCNA_00175 | type II secretion pathway protein G | 14 | 0,038674033 | 23,84505002 | 0,065870304 | NU |
| CCNA_01778 | penicillin-binding protein | 40 | 0,03868472 | 83,45767508 | 0,080713419 | V |
| CCNA_02208 | thymidylate synthase | 26 | 0,038748137 | 1,834234617 | 0,002733584 | F |
| CCNA_00657 | type I restriction-modification system specificity subunit | 42 | 0,038817006 | 62,36397698 | 0,057637687 | V |
| CCNA_02091 | hypothetical protein | 29 | 0,039030956 | 42,18739619 | 0,056779806 | NA |
| CCNA_00076 | hypothetical protein | 12 | 0,039087948 | 16,50811155 | 0,05377235 | NA |

| | | | | | | |
|---|---|---|---|---|---|---|
| CCNA_02098 | molybdopterin-guanine dinucleotide biosynthesis protein A | 28 | 0,039106145 | 56,86127313 | 0,079415186 | NA |
| CCNA_00576 | oligopeptide-binding protein oppA | 39 | 0,039156627 | 84,37479239 | 0,084713647 | NA |
| CCNA_03623 | transcriptional regulator | 38 | 0,039378238 | 56,86127313 | 0,058923599 | KT |
| CCNA_R0099 | small non-coding RNA | 8 | 0,039408867 | 17,42522886 | 0,085838566 | NA |
| CCNA_01119 | hypothetical protein | 12 | 0,039473684 | 31,18198849 | 0,102572331 | NA |
| CCNA_02650 | anhydro-N-acetylmuramyl-tripeptide amidase | 23 | 0,0395189 | 25,67928464 | 0,044122482 | V |
| CCNA_03661 | hypothetical protein | 4 | 0,03960396 | 6,41982116 | 0,063562586 | NA |
| CCNA_01656 | endonuclease/exonuclease/phosphatase family protein | 31 | 0,039692702 | 48,60721735 | 0,062237154 | S |
| CCNA_02038 | hypothetical protein | 85 | 0,039831303 | 48,60721735 | 0,022777515 | NA |
| CCNA_02679 | hypothetical protein | 24 | 0,03986711 | 33,01622311 | 0,054844224 | NA |
| CCNA_01491 | cystathionine beta-lyase | 37 | 0,03987069 | 55,02703851 | 0,059296378 | E |
| CCNA_01757 | dimethyladenosine transferase | 27 | 0,039881832 | 18,34234617 | 0,027093569 | J |
| CCNA_R0102 | small non-coding RNA | 2 | 0,04 | 0 | 0 | J |
| CCNA_R0105 | small non-coding RNA | 10 | 0,04 | 7,336938469 | 0,029347754 | NA |
| CCNA_00518 | LSU ribosomal protein L25P | 19 | 0,04 | 18,34234617 | 0,038615466 | NA |
| CCNA_03185 | glutathione S-transferase | 20 | 0,04 | 28,43063657 | 0,056861273 | NA |
| CCNA_03576 | hypothetical protein | 9 | 0,04 | 26,59640195 | 0,118206231 | O |
| CCNA_03513 | cytochrome c oxidase subunit III coxC | 28 | 0,040114613 | 33,93334042 | 0,048615101 | C |
| CCNA_01929 | hypothetical protein | 13 | 0,040123457 | 12,83964232 | 0,039628526 | NA |
| CCNA_03525 | hypothetical protein | 11 | 0,04029304 | 18,34234617 | 0,067188081 | NA |
| CCNA_01885 | short chain dehydrogenase | 25 | 0,040322581 | 36,68469234 | 0,059168859 | IQR |
| CCNA_R0142 | small non-coding RNA | 5 | 0,040322581 | 9,171173086 | 0,073961073 | NA |
| CCNA_01806 | ferritin-like domain protein | 47 | 0,040343348 | 79,78920585 | 0,068488589 | NA |
| CCNA_02273 | glutathione peroxidase | 8 | 0,04040404 | 28,43063657 | 0,143589074 | NA |
| CCNA_03895 | hypothetical protein | 9 | 0,040540541 | 7,336938469 | 0,033049272 | NA |
| CCNA_03826 | hypothetical protein | 18 | 0,040540541 | 41,27027889 | 0,092951079 | R |
| CCNA_00153 | GrpE protein | 21 | 0,042 | 9,171173086 | 0,018342346 | O |
| CCNA_R0005 | 4,5S RNA | 2 | 0,042553191 | 0 | 0 | NA |
| CCNA_00195 | 3-isopropylmalate dehydratase small subunit | 21 | 0,043209877 | 9,171173086 | 0,018870727 | E |
| CCNA_01739 | LSU ribosomal protein L9P | 21 | 0,044871795 | 1,834234617 | 0,003919305 | J |
| *CCNA_01084* | *hypothetical protein* | *45* | 0,045592705 | 15,59099425 | 0,015796347 | NA |
| CCNA_02855 | hypothetical protein | 12 | 0,04743083 | 2,751351926 | 0,010874909 | S |
| CCNA_03150 | hypothetical protein | 6 | 0,047619048 | 1,834234617 | 0,014557418 | NA |
| CCNA_02705 | cold shock protein | 8 | 0,047904192 | 2,751351926 | 0,016475161 | K |

| | | | | | | |
|---|---|---|---|---|---|---|
| CCNA_00932 | 6,7-dimethyl-8-ribityllumazine synthase | 18 | 0,048780488 | 2,751351926 | 0,007456238 | H |
| CCNA_02088 | HesB protein family | 13 | 0,04887218 | 0,917117309 | 0,003447809 | S |
| CCNA_01433 | integration host factor alpha-subunit | 12 | 0,049586777 | 4,585586543 | 0,018948705 | L |
| CCNA_01989 | (3R)-hydroxymyristoyl-(acyl-carrier-protein) dehydratase | 19 | 0,049608355 | 5,502703851 | 0,014367373 | I |
| CCNA_R0037 | tRNA Glu | 3 | 0,05 | 0,917117309 | 0,015285288 | NA |
| *CCNA_03517* | *cytochrome c oxidase polypeptide I coxA* | *67* | 0,050527903 | 24,76216733 | 0,018674334 | C |
| CCNA_02521 | endonuclease involved in recombination | 19 | 0,050531915 | 4,585586543 | 0,012195709 | L |
| CCNA_03748 | dTDP-4-dehydrorhamnose 3,5-epimerase | 23 | 0,051685393 | 4,585586543 | 0,010304689 | M |
| CCNA_03107 | cytidine deaminase | 21 | 0,052109181 | 8,254055777 | 0,020481528 | F |
| CCNA_02064 | UDP-3-O-(3-hydroxymyristoyl) N-acetylglucosamine deacetylase LpxC | 38 | 0,053072626 | 11,92252501 | 0,016651571 | M |
| CCNA_00758 | protein translation Elongation factor P EF-P | 24 | 0,053097345 | 7,336938469 | 0,016232165 | J |
| CCNA_01630 | ribonuclease III | 30 | 0,054054054 | 1,834234617 | 0,003304927 | K |
| *CCNA_01953* | *hypothetical protein* | *4* | 0,054054054 | 0,917117309 | 0,012393477 | NA |
| CCNA_00382 | adenine-specific methyltransferase ccrM | 47 | 0,054651163 | 15,59099425 | 0,018129063 | L |
| CCNA_03078 | hypothetical protein | 5 | 0,054945055 | 0,917117309 | 0,010078212 | K |
| CCNA_03961 | hypothetical protein | 10 | 0,055555556 | 1,834234617 | 0,010190192 | NA |
| CCNA_01819 | RNA-binding protein, Hfq family | 11 | 0,055555556 | 3,668469234 | 0,018527622 | R |
| CCNA_01979 | LexA repressor | 33 | 0,056027165 | 7,336938469 | 0,012456602 | KT |
| CCNA_01544 | hypothetical protein | 10 | 0,056497175 | 3,668469234 | 0,020725815 | NA |
| CCNA_01079 | hypothetical protein | 8 | 0,057142857 | 2,751351926 | 0,019652514 | NA |
| CCNA_02021 | hypothetical protein | 11 | 0,057591623 | 2,751351926 | 0,014404984 | NA |
| CCNA_01521 | hypothetical protein | 38 | 0,058371736 | 10,08829039 | 0,015496606 | NA |
| CCNA_00735 | hypothetical protein BamF | 27 | 0,059340659 | 7,336938469 | 0,016125139 | NA |
| CCNA_03931 | hypothetical protein | 14 | 0,059574468 | 4,585586543 | 0,019513134 | NA |
| CCNA_01671 | diguanylate receptor protein dgrA | 17 | 0,059859155 | 4,585586543 | 0,016146431 | NA |
| CCNA_02242 | PHB granule-associated protein, phasin2 | 21 | 0,060344828 | 5,502703851 | 0,015812367 | S |
| CCNA_02962 | hypothetical protein | 11 | 0,060773481 | 2,751351926 | 0,015200839 | S |
| CCNA_02987 | hypothetical protein | 12 | 0,062827225 | 2,751351926 | 0,014404984 | NA |
| CCNA_01628 | hypothetical protein | 15 | 0,063829787 | 4,585586543 | 0,019513134 | NA |
| CCNA_03030 | cytochrome c | 36 | 0,065693431 | 4,585586543 | 0,008367859 | C |
| CCNA_00239 | HPR(SER) kinase/phosphatase HprK | 23 | 0,066091954 | 5,502703851 | 0,015812367 | T |
| CCNA_R0135 | small non-coding RNA | 4 | 0,066666667 | 0,917117309 | 0,015285288 | NA |
| CCNA_03264 | hypothetical protein | 11 | 0,067484663 | 2,751351926 | 0,01687946 | NA |
| ***CCNA_01667*** | ***two-component response regulator kdpE*** | ***38*** | 0,068100358 | 1,834234617 | 0,003287159 | TK |
| CCNA_R004*9* | *nan* | *12* | 0,070175439 | 0,917117309 | 0,005363259 | NA |

| | | | | | | |
|---|---|---|---|---|---|---|
| CCNA_00771 | hypothetical protein | 7 | 0,071428571 | 1,834234617 | 0,01871668 | NA |
| ***CCNA_01664*** | ***potassium-transporting ATPase B chain kdpB*** | ***119*** | 0,072208738 | 2,751351926 | 0,00166951 | P |
| CCNA_R0181 | small non-coding RNA | 5 | 0,073529412 | 0,917117309 | 0,013487019 | NA |
| CCNA_03115 | cobaltochelatase cobS subunit | 62 | 0,076732673 | 14,67387694 | 0,018160739 | R |
| CCNA_00852 | transcriptional regulator | 38 | 0,077235772 | 0 | 0 | K |
| *CCNA_01427* | *lipoprotein, SmpA/OmlA family BamE* | *30* | 0,077720207 | 5,502703851 | 0,014255709 | J |
| CCNA_02463 | UDP-N-acetylglucosamine 4-epimerase | 61 | 0,078205128 | 11,92252501 | 0,015285288 | MG |
| CCNA_03650 | mannose-1-phosphate guanyltransferase | 46 | 0,079037801 | 9,171173086 | 0,015758029 | MJ |
| CCNA_01724 | hypothetical protein | 48 | 0,079734219 | 1,834234617 | 0,003046901 | S |
| CCNA_02663 | DnaK suppressor protein dksA | 28 | 0,081871345 | 5,502703851 | 0,016089777 | T |
| CCNA_00080 | LexA-related transcriptional repressor | 43 | 0,083820663 | 0,917117309 | 0,001787753 | K |
| ***CCNA_01663*** | ***potassium-transporting ATPase A chain kdpA*** | ***115*** | 0,083941606 | 2,751351926 | 0,002008286 | P |
| CCNA_03978 | hypothetical protein | 10 | 0,086206897 | 0 | 0 | NA |
| CCNA_R0049 | nan | 20 | 0,088495575 | 3,668469234 | 0,016232165 | NA |
| ***CCNA_01666*** | ***osmosensitive K+ channel histidine kinase kdpD*** | ***205*** | 0,095704949 | 42,18739619 | 0,01969533 | T |
| CCNA_03954 | hypothetical protein | 22 | 0,096069869 | 3,668469234 | 0,016019516 | NA |
| CCNA_00708 | cobaltochelatase cobT subunit | 152 | 0,098573281 | 29,34775387 | 0,019032266 | H |
| CCNA_R0059 | tRNA Thr | 6 | 0,1 | 0 | 0 | NA |
| CCNA_01525 | flagellar biosynthesis repressor flbT | 38 | 0,11143695 | 4,585586543 | 0,013447468 | N |
| *CCNA_00919* | *hypothetical protein* | *21* | 0,116666667 | 2,751351926 | 0,015285288 | NA |
| CCNA_R0100 | small non-coding RNA chvR | 8 | 0,119402985 | 0 | 0 | NA |
| CCNA_R0143 | small non-coding RNA | 8 | 0,126984127 | 0,917117309 | 0,014557418 | NA |
| CCNA_01436 | hypothetical protein | 14 | 0,142857143 | 1,834234617 | 0,01871668 | NA |
| CCNA_01756 | hypothetical protein | 14 | 0,147368421 | 0,917117309 | 0,009653866 | NA |
| CCNA_R0166 | small non-coding RNA | 17 | 0,354166667 | 0,917117309 | 0,019106611 | NA |

**Table S3. Transcriptome in low $K^+$ concentrations determined by RNA-seq.** (A) genes with expression upregulated in the WT cells grown in M2G-K 0.025 mM $K^+$ compared to WT cells grown in M2G-K 0.5 mM $K^+$.

Genes found in Tn-Seq_WT (Table S2) are highlighted in **bold**.
Genes found in RNA-Seq_Δ*kdpE* (Table S4) are highlighted in yellow.
Genes found in ChIP-Seq_WT (Table S5 or S6) are highlighted in *italics*.

| Top hit | Gene ID | Description | $log_2$ (FC WT Low $K^+$ / WT High $K^+$) | P-value | P-adj |
|---|---|---|---|---|---|

| | | | | | |
|---|---|---|---|---|---|
| ***1*** | ***CCNA_01664*** | ***potassium-transporting ATPase B chain*** | ***7,037843115*** | ***4,64E-180*** | ***1,88E-176*** |
| ***2*** | ***CCNA_01663*** | ***potassium-transporting ATPase A chain*** | ***6,690420041*** | ***6,70E-115*** | ***6,77E-112*** |
| ***3*** | ***CCNA_01665*** | ***potassium-transporting ATPase C chain*** | ***5,685459296*** | ***6,88E-168*** | ***1,39E-164*** |
| ***4*** | ***CCNA_01666*** | ***osmosensitive K+ channel histidine kinase kdpD*** | ***4,621986028*** | ***2,11E-160*** | ***2,84E-157*** |
| *5* | *CCNA_02727* | *PhoH protein* | *4,136136312* | *9,20E-58* | *6,21E-55* |
| 6 | CCNA_03313 | hypothetical protein | 4,106537473 | 4,57E-17 | 1,53E-15 |
| 7 | CCNA_03970 | hypothetical protein | 3,486079541 | 9,49E-34 | 2,02E-31 |
| 8 | CCNA_01701 | xylose isomerase family protein | 3,334117778 | 6,77E-32 | 1,14E-29 |
| 9 | CCNA_01859 | TonB-dependent receptor | 3,247710055 | 1,54E-51 | 8,91E-49 |
| 10 | CCNA_01858 | transcriptional regulator, AraC family | 3,226194837 | 9,16E-35 | 2,42E-32 |
| *11* | *CCNA_03312* | *hypothetical protein* | *3,201915294* | *2,84E-24* | *2,50E-22* |
| 12 | CCNA_03689 | alanine dehydrogenase | 3,18829282 | 4,06E-34 | 9,25E-32 |
| 13 | CCNA_02433 | hypothetical protein | 3,06124785 | 1,56E-29 | 2,25E-27 |
| 14 | CCNA_03973 | hypothetical protein | 3,048041159 | 2,55E-29 | 3,56E-27 |
| 15 | CCNA_02491 | protocatechuate 3,4-dioxygenase subunit beta | 3,03171835 | 2,15E-23 | 1,64E-21 |
| 16 | CCNA_01706 | dehydrogluconate dehydrogenase | 2,953314068 | 4,89E-26 | 4,94E-24 |
| 17 | CCNA_03487 | hypothetical protein | 2,934633544 | 5,09E-19 | 2,34E-17 |
| 18 | CCNA_03660 | Usg protein | 2,931396301 | 3,40E-21 | 1,99E-19 |
| 19 | CCNA_01702 | NAD-dependent oxidoreductase | 2,874386106 | 4,48E-27 | 5,18E-25 |
| 20 | CCNA_02489 | acetate/3-ketoacid CoA transferase, subunit B | 2,840499457 | 1,84E-18 | 8,02E-17 |
| 21 | CCNA_02490 | 3-ketoacyl-CoA thiolase | 2,823008911 | 3,35E-09 | 3,24E-08 |
| 22 | CCNA_01855 | iron/manganese superoxide dismutase | 2,789284439 | 4,11E-34 | 9,25E-32 |
| 23 | CCNA_03906 | conserved hypothetical protein | 2,774144712 | 2,93E-07 | 1,81E-06 |
| 24 | CCNA_03593 | integral membrane protein | 2,753947897 | 8,68E-38 | 3,19E-35 |
| 25 | CCNA_00754 | hypothetical protein | 2,753336422 | 8,23E-20 | 3,96E-18 |
| 26 | CCNA_03945 | hypothetical protein | 2,751746652 | 2,20E-33 | 4,44E-31 |
| 27 | CCNA_02493 | 3-carboxy-cis,cis-muconate cycloisomerase | 2,725171928 | 2,72E-25 | 2,50E-23 |
| 28 | CCNA_01860 | rare lipoprotein B precursor | 2,695388167 | 1,95E-32 | 3,43E-30 |
| 29 | CCNA_02492 | protocatechuate 3,4-dioxygenase subunit alpha | 2,680510219 | 1,33E-21 | 8,02E-20 |
| 30 | CCNA_02604 | host cell attachment protein | 2,639574744 | 2,11E-06 | 1,07E-05 |
| 31 | CCNA_03096 | TonB-dependent receptor | 2,593632516 | 6,52E-41 | 2,93E-38 |
| 32 | CCNA_00882 | hypothetical protein | 2,577288273 | 1,37E-26 | 1,46E-24 |
| 33 | CCNA_03400 | hypothetical protein | 2,566325916 | 1,22E-22 | 8,21E-21 |
| **34** | **CCNA_01667** | **two-component response regulator kdpE** | **2,556970902** | **8,30E-64** | **6,72E-61** |
| 35 | CCNA_03138 | peroxidase/catalase katG | 2,534302299 | 2,50E-31 | 4,05E-29 |
| 36 | CCNA_03269 | glucokinase | 2,499496439 | 4,41E-31 | 6,87E-29 |

| | | | | | |
|---|---|---|---|---|---|
| 37 | CCNA_01519 | universal stress protein family | 2,49858221 | 2,55E-07 | 1,62E-06 |
| 38 | CCNA_01700 | nucleoside permease | 2,481469667 | 8,92E-26 | 8,80E-24 |
| 39 | CCNA_03926 | hypothetical protein | 2,470222203 | 1,27E-17 | 4,72E-16 |
| 40 | CCNA_01705 | hypothetical protein | 2,444574931 | 1,07E-20 | 5,77E-19 |
| 41 | CCNA_01049 | hypothetical protein | 2,427171619 | 5,86E-43 | 2,96E-40 |
| 42 | CCNA_01236 | cytosolic protein | 2,404331998 | 1,94E-32 | 3,43E-30 |
| 43 | CCNA_02872 | phage prohead protease | 2,395216009 | 2,90E-38 | 1,17E-35 |
| 44 | CCNA_02871 | gene transfer agent (GTA)-related protein | 2,38810748 | 2,80E-16 | 8,46E-15 |
| 45 | CCNA_01707 | hypothetical protein | 2,378383889 | 8,45E-07 | 4,74E-06 |
| 46 | CCNA_01269 | GcrB protein | 2,348238869 | 3,04E-23 | 2,26E-21 |
| 47 | CCNA_02869 | gene transfer agent (GTA) protein | 2,340542131 | 4,59E-14 | 1,07E-12 |
| 48 | CCNA_03695 | aldehyde dehydrogenase | 2,339355855 | 3,81E-18 | 1,56E-16 |
| 49 | CCNA_R0147 | small non-coding RNA | 2,336475826 | 8,79E-09 | 7,81E-08 |
| 50 | CCNA_00801 | cytochrome bd-type quinol oxidase, subunit 2 cydB | 2,31843058 | 8,51E-16 | 2,44E-14 |
| 51 | CCNA_01520 | hypothetical protein | 2,300690604 | 8,18E-07 | 4,60E-06 |
| 52 | CCNA_03298 | TonB-dependent outer membrane receptor | 2,293944605 | 6,22E-35 | 1,80E-32 |
| *53* | *CCNA_01093* | *hypothetical protein* | *2,29010297* | *8,29E-15* | *2,18E-13* |
| 54 | CCNA_03624 | sodium bicarbonate cotransporter | 2,2759875 | 2,87E-35 | 8,94E-33 |
| 55 | CCNA_00800 | cytochrome bd-type quinol oxidase, subunit 1 cydA | 2,257772954 | 5,12E-06 | 2,32E-05 |
| 56 | CCNA_01857 | transcriptional regulator, AraC family | 2,253369114 | 6,09E-21 | 3,42E-19 |
| 57 | CCNA_01021 | TonB-dependent receptor | 2,236283168 | 2,83E-25 | 2,54E-23 |
| 58 | CCNA_02432 | cytosine/uracil/thiamine/allantoin permease family protein | 2,225040404 | 1,66E-35 | 5,59E-33 |
| 59 | CCNA_01237 | hypothetical protein | 2,222147589 | 2,64E-30 | 3,95E-28 |
| 60 | CCNA_03447 | hypothetical protein | 2,219728775 | 3,86E-13 | 7,57E-12 |
| 61 | CCNA_03882 | phage gp6-like head-tail connector protein | 2,216001075 | 9,22E-18 | 3,52E-16 |
| 62 | CCNA_02715 | MASE1-family sensor histidine kinase | 2,194203742 | 7,20E-23 | 5,11E-21 |
| 63 | CCNA_01703 | IolE-related protein | 2,166360699 | 1,14E-20 | 6,05E-19 |
| 64 | CCNA_01270 | hypothetical protein | 2,159902647 | 1,24E-20 | 6,54E-19 |
| 65 | CCNA_03444 | TonB-dependent receptor | 2,155830395 | 6,78E-33 | 1,31E-30 |
| 66 | CCNA_01601 | hypothetical protein | 2,15310564 | 2,62E-16 | 7,96E-15 |
| 67 | CCNA_00994 | oxidoreductase, GMC family | 2,144454098 | 1,25E-21 | 7,80E-20 |
| 68 | CCNA_02494 | 3-oxoadipate enol-lactonase/4-carboxymuconolactone decarboxylase | 2,120527106 | 1,27E-21 | 7,81E-20 |
| 69 | CCNA_03267 | transcriptional regulatory protein | 2,116371563 | 2,54E-15 | 7,03E-14 |
| 70 | CCNA_00551 | hypothetical protein | 2,110190263 | 1,72E-26 | 1,79E-24 |
| 71 | CCNA_00279 | NAD(P)H dehydrogenase (quinone) | 2,10318198 | 2,55E-05 | 0,000100553 |
| 72 | CCNA_03694 | transcriptional regulator | 2,075627428 | 8,26E-18 | 3,22E-16 |
| 73 | CCNA_01588 | ABC transporter permease protein | 2,066794794 | 9,63E-05 | 0,000337429 |

| | | | | | |
|---|---|---|---|---|---|
| 74 | CCNA_03263 | TonB-dependent receptor | 2,066228854 | 9,58E-35 | 2,42E-32 |
| 75 | CCNA_01017 | acyl-CoA synthetase | 2,061852423 | 4,66E-21 | 2,65E-19 |
| 76 | CCNA_02880 | phage DNA packaging protein | 2,060587864 | 3,36E-24 | 2,83E-22 |
| 77 | CCNA_02868 | hypothetical protein | 2,059759675 | 3,14E-07 | 1,93E-06 |
| 78 | CCNA_02603 | phosphatidylserine decarboxylase | 2,035963159 | 8,41E-12 | 1,31E-10 |
| 79 | CCNA_01822 | melibiose carrier protein | 2,015984284 | 4,28E-19 | 1,99E-17 |
| 80 | CCNA_00127 | hypothetical protein | 2,012212854 | 1,82E-11 | 2,68E-10 |
| 81 | CCNA_00709 | hypothetical protein | 2,011987156 | 3,93E-17 | 1,33E-15 |
| 82 | CCNA_02283 | HNH endonuclease family protein | 1,979705138 | 3,39E-27 | 4,15E-25 |
| 83 | CCNA_01120 | hypothetical protein | 1,974997276 | 4,09E-14 | 9,56E-13 |
| 84 | CCNA_02251 | methylglutaconyl-CoA hydratase | 1,965268645 | 4,07E-12 | 6,74E-11 |
| 85 | CCNA_03327 | hypothetical protein | 1,949305952 | 5,25E-16 | 1,53E-14 |
| 86 | CCNA_02862 | phage host specificity protein | 1,943408153 | 1,50E-13 | 3,23E-12 |
| *87* | *CCNA_01092* | *hypothetical protein* | *1,938618638* | *2,61E-12* | *4,49E-11* |
| 88 | CCNA_03242 | succinate-semialdehyde dehydrogenase (NADP+) | 1,928983757 | 9,32E-19 | 4,15E-17 |
| 89 | CCNA_01523 | acetyltransferase flmH | 1,926114231 | 0,000821079 | 0,002295845 |
| 90 | CCNA_00802 | protein ybgT | 1,920840782 | 1,86E-11 | 2,72E-10 |
| 91 | CCNA_01874 | transposase | 1,918810382 | 4,17E-10 | 4,71E-09 |
| 92 | CCNA_02606 | hybrid sensor histidine kinase/receiver domain protein | 1,916433145 | 8,46E-22 | 5,44E-20 |
| 93 | CCNA_02570 | transporter | 1,911152261 | 7,56E-21 | 4,19E-19 |
| 94 | CCNA_03579 | hypothetical protein | 1,906819483 | 3,14E-14 | 7,46E-13 |
| 95 | CCNA_00969 | two component sensor histidine kinase | 1,906316416 | 2,48E-13 | 5,06E-12 |
| 96 | CCNA_00437 | methyl-accepting chemotaxis protein | 1,90068667 | 4,35E-24 | 3,59E-22 |
| 97 | CCNA_02709 | CBS domain containing protein | 1,899475944 | 1,80E-17 | 6,46E-16 |
| 98 | CCNA_01606 | Soj/ParA-related ATPase protein | 1,898735176 | 3,59E-20 | 1,77E-18 |
| 99 | CCNA_02488 | acetate/3-ketoacid CoA transferase, subunit A | 1,897458153 | 0,000555824 | 0,001620218 |
| 100 | CCNA_02487 | 4-hydroxybenzoate 3-monooxygenase | 1,894314567 | 1,66E-10 | 2,02E-09 |
| 101 | CCNA_01499 | acetyl-CoA acetyltransferase | 1,893067998 | 8,89E-18 | 3,42E-16 |
| 102 | CCNA_02870 | phage protein | 1,885199149 | 1,88E-15 | 5,28E-14 |
| 103 | CCNA_03573 | D-aminoacylase | 1,881772367 | 1,67E-23 | 1,33E-21 |
| 104 | CCNA_01476 | CRP-family transcription regulator ftrB | 1,879753477 | 0,000645928 | 0,001850097 |
| 105 | CCNA_03446 | feruloyl-CoA synthetase | 1,875277476 | 2,83E-27 | 3,58E-25 |
| 106 | CCNA_00416 | hypothetical protein | 1,865240123 | 5,33E-10 | 5,90E-09 |
| 107 | CCNA_03488 | hypothetical protein | 1,857851946 | 2,70E-18 | 1,14E-16 |
| 108 | CCNA_03169 | hypothetical protein | 1,846578923 | 5,67E-15 | 1,53E-13 |
| 109 | CCNA_03936 | hypothetical protein | 1,843675924 | 1,03E-07 | 7,18E-07 |
| 110 | CCNA_02684 | cyclopropane fatty acid synthase-like methyltransferase | 1,82496084 | 2,21E-25 | 2,08E-23 |
| 111 | CCNA_02110 | hypothetical protein | 1,812184555 | 2,83E-20 | 1,45E-18 |

| | | | | | |
|---|---|---|---|---|---|
| 112 | CCNA_03188 | epoxide hydrolase | 1,787406155 | 1,38E-11 | 2,08E-10 |
| 113 | CCNA_02660 | pyridoxamine 5 -phosphate oxidase family protein | 1,769467959 | 4,66E-14 | 1,07E-12 |
| 114 | CCNA_02703 | urate oxidase | 1,761679867 | 1,02E-14 | 2,61E-13 |
| 115 | CCNA_00987 | hypothetical protein | 1,754644673 | 1,09E-22 | 7,50E-21 |
| 116 | CCNA_02632 | SOUL domain heme-binding protein | 1,751644606 | 6,84E-15 | 1,82E-13 |
| *117* | *CCNA_02728* | *hypothetical protein* | *1,744722285* | *9,18E-23* | *6,41E-21* |
| 118 | CCNA_03696 | acetyl-coenzyme A synthetase | 1,739306617 | 3,51E-17 | 1,21E-15 |
| 119 | CCNA_02111 | pilF-related tetratricopeptide repeat protein | 1,738676337 | 1,71E-17 | 6,16E-16 |
| 120 | CCNA_01477 | oxygen-independent coproporphyrinogen-III oxidase hemN | 1,735205781 | 0,000935145 | 0,002570378 |
| 121 | CCNA_02885 | isochorismatase family protein | 1,724704331 | 1,15E-13 | 2,51E-12 |
| 122 | CCNA_02659 | two-component receiver domain protein | 1,719636096 | 1,03E-13 | 2,27E-12 |
| *123* | *CCNA_01536* | *hypothetical protein* | *1,712548378* | *3,70E-17* | *1,27E-15* |
| 124 | CCNA_01890 | enoyl-CoA hydratase/isomerase family protein | 1,70885786 | 1,26E-16 | 3,94E-15 |
| 125 | CCNA_02877 | portal protein | 1,707339983 | 1,36E-14 | 3,39E-13 |
| 126 | CCNA_00854 | metallo-beta-lactamase protein | 1,703723359 | 0,000481589 | 0,001427478 |
| 127 | CCNA_00799 | ABC transporter, ATP-binding protein cydD | 1,702065697 | 1,02E-08 | 8,94E-08 |
| 128 | CCNA_02200 | cytochrome c-family protein | 1,671353766 | 0,000811752 | 0,002274479 |
| 129 | CCNA_02865 | phage protein | 1,66885283 | 5,94E-10 | 6,52E-09 |
| *130* | *CCNA_03091* | *hypothetical protein NstA* | *1,649558462* | *7,38E-14* | *1,68E-12* |
| 131 | CCNA_03124 | NAD/mycothiol-dependent formaldehyde dehydrogenase | 1,647037747 | 1,54E-14 | 3,81E-13 |
| *132* | *CCNA_03711* | *ribosome-associated factor Y* | *1,644362024* | *1,51E-12* | *2,69E-11* |
| 133 | CCNA_03616 | sulfite reductase (NADPH) flavoprotein alpha-component | 1,643824808 | 3,07E-23 | 2,26E-21 |
| 134 | CCNA_03572 | cytosolic protein | 1,635547954 | 1,49E-18 | 6,54E-17 |
| 135 | CCNA_03535 | fatty acid desaturase | 1,634713505 | 8,47E-11 | 1,11E-09 |
| 136 | CCNA_02796 | hypothetical protein | 1,633811112 | 2,46E-08 | 1,98E-07 |
| *137* | *CCNA_02517* | *hypothetical protein* | *1,630509135* | *1,55E-13* | *3,29E-12* |
| 138 | CCNA_01290 | hypothetical protein | 1,624229317 | 1,16E-26 | 1,27E-24 |
| *139* | *CCNA_01091* | *peptide methionine sulfoxide reductase* | *1,622088886* | *9,31E-10* | *9,87E-09* |
| 140 | CCNA_02895 | TonB-dependent receptor | 1,616658252 | 3,18E-24 | 2,74E-22 |
| 141 | CCNA_01852 | major facilitator superfamily transporter | 1,614440704 | 3,77E-27 | 4,49E-25 |
| 142 | CCNA_03605 | hypothetical protein | 1,612686853 | 2,12E-16 | 6,49E-15 |
| 143 | CCNA_02286 | hypothetical protein | 1,596930102 | 0,003429563 | 0,008076842 |
| 144 | CCNA_03928 | hypothetical protein | 1,59498858 | 2,63E-22 | 1,72E-20 |
| 145 | CCNA_03245 | omega-amino acid-pyruvate aminotransferase | 1,592577223 | 1,01E-12 | 1,84E-11 |
| 146 | CCNA_03567 | acyl-CoA dehydrogenase | 1,591897632 | 4,62E-14 | 1,07E-12 |
| 147 | CCNA_03617 | hypothetical protein | 1,579472028 | 1,52E-13 | 3,26E-12 |

| | | | | | |
|---|---|---|---|---|---|
| 148 | CCNA_02977 | RNA polymerase ECF-type sigma factor sigU | 1,577870422 | 1,21E-11 | 1,84E-10 |
| 149 | CCNA_00813 | dimethylaniline monooxygenase (N-oxide forming) | 1,576166774 | 2,95E-20 | 1,49E-18 |
| 150 | CCNA_01467 | cytochrome cbb3 oxidase subunit I ccoN | 1,568167742 | 0,001134555 | 0,003040863 |
| 151 | CCNA_01498 | DNA-dependent DNA polymerase III subunit alpha | 1,566237114 | 2,20E-12 | 3,81E-11 |
| 152 | CCNA_01260 | hypothetical protein | 1,56563376 | 2,00E-25 | 1,93E-23 |
| 153 | CCNA_02146 | hypothetical protein | 1,565409853 | 2,67E-12 | 4,58E-11 |
| 154 | CCNA_01468 | cytochrome cbb3 oxidase, cytochrome c subunit ccoO | 1,564431828 | 2,09E-08 | 1,71E-07 |
| 155 | CCNA_01466 | hypothetical protein | 1,557860971 | 0,002923104 | 0,00705689 |
| 156 | CCNA_00817 | L-sorbosone dehydrogenase | 1,533274243 | 2,13E-14 | 5,16E-13 |
| 157 | CCNA_02861 | phage host specificity protein | 1,527085686 | 1,73E-16 | 5,34E-15 |
| 158 | CCNA_03369 | hypothetical protein | 1,523489267 | 9,94E-13 | 1,84E-11 |
| 159 | CCNA_00061 | cytochrome p450 | 1,522171024 | 8,85E-13 | 1,66E-11 |
| 160 | CCNA_00687 | hypothetical protein | 1,510668445 | 1,91E-10 | 2,30E-09 |
| 161 | CCNA_03669 | outer membrane protein | 1,508447745 | 6,08E-17 | 1,98E-15 |
| 162 | CCNA_00838 | alpha-glucosidase | 1,494783417 | 1,46E-10 | 1,81E-09 |
| 163 | CCNA_01072 | hypothetical protein | 1,494163805 | 2,94E-08 | 2,33E-07 |
| 164 | CCNA_01194 | TonB-dependent receptor | 1,492712519 | 2,62E-22 | 1,72E-20 |
| 165 | CCNA_02797 | hypothetical protein | 1,490802531 | 2,71E-08 | 2,16E-07 |
| 166 | CCNA_00993 | long-chain-fatty-acid--CoA ligase | 1,486784969 | 3,42E-20 | 1,71E-18 |
| 167 | CCNA_02685 | hypothetical protein | 1,486737483 | 2,84E-14 | 6,80E-13 |
| 168 | CCNA_01287 | epoxide hydrolase | 1,484893636 | 1,43E-16 | 4,44E-15 |
| 169 | CCNA_02875 | hypothetical protein | 1,483632587 | 3,23E-06 | 1,55E-05 |
| 170 | CCNA_01854 | transcriptional regulator, GntR family | 1,482659382 | 7,14E-14 | 1,63E-12 |
| 171 | CCNA_03618 | SCO1/SenC family protein | 1,480914173 | 6,33E-15 | 1,70E-13 |
| 172 | CCNA_00996 | enoyl-CoA hydratase | 1,477435234 | 2,41E-11 | 3,45E-10 |
| 173 | CCNA_00798 | ABC transporter, ATP-binding protein cydC | 1,46699211 | 7,70E-09 | 6,97E-08 |
| 174 | CCNA_03374 | hypothetical protein | 1,46629447 | 2,16E-17 | 7,65E-16 |
| 175 | CCNA_03065 | cytochrome c family protein | 1,465913437 | 1,29E-11 | 1,96E-10 |
| 176 | CCNA_02250 | methylcrotonyl-CoA carboxylase biotin-containing subunit | 1,463517181 | 6,84E-08 | 4,97E-07 |
| 177 | CCNA_00064 | methyl-accepting chemotaxis protein | 1,46256757 | 9,27E-10 | 9,84E-09 |
| 178 | CCNA_02495 | 4-hydroxybenzoate transporter | 1,459937843 | 1,27E-12 | 2,30E-11 |
| 179 | CCNA_01382 | long-chain-fatty-acid--CoA ligase | 1,457672524 | 3,04E-16 | 9,13E-15 |
| 180 | CCNA_00263 | hypothetical protein | 1,455183112 | 9,37E-17 | 3,01E-15 |
| 181 | CCNA_01259 | class II aldolase and Adducin protein | 1,453014595 | 8,41E-13 | 1,59E-11 |
| *182* | *CCNA_00627* | *hypothetical protein* | *1,437585321* | *7,28E-08* | *5,25E-07* |
| 183 | CCNA_02702 | xanthine dehydrogenase small subunit | 1,433871649 | 6,04E-12 | 9,73E-11 |
| 184 | CCNA_01291 | phosphotransferase family protein | 1,433332713 | 1,48E-17 | 5,39E-16 |

| | | | | | |
|---|---|---|---|---|---|
| 185 | CCNA_02686 | N-carbamoyl-L-amino acid hydrolase | 1,431647756 | 1,22E-10 | 1,54E-09 |
| 186 | CCNA_02863 | Spr-family cell wall-associated hydrolase | 1,429382636 | 1,13E-05 | 4,80E-05 |
| 187 | CCNA_00820 | 3-ketoacyl-CoA thiolase | 1,427959457 | 7,57E-13 | 1,44E-11 |
| 188 | CCNA_00716 | hypothetical protein | 1,424196672 | 1,15E-16 | 3,63E-15 |
| 189 | CCNA_01830 | TonB-dependent receptor | 1,423181201 | 1,59E-11 | 2,38E-10 |
| 190 | CCNA_01185 | hypothetical protein | 1,419834434 | 0,005928509 | 0,013186776 |
| 191 | CCNA_03615 | TonB dependent receptor | 1,41954622 | 6,38E-18 | 2,56E-16 |
| 192 | CCNA_01410 | cyclohexanone monooxygenase | 1,41798581 | 7,32E-12 | 1,17E-10 |
| 193 | CCNA_00815 | lignostilbene-alpha,beta-dioxygenase | 1,413245162 | 9,17E-17 | 2,97E-15 |
| 194 | CCNA_01704 | cytochrome c | 1,412654574 | 1,70E-12 | 2,97E-11 |
| 195 | CCNA_01303 | hypothetical protein | 1,411887819 | 0,002572706 | 0,006297138 |
| 196 | CCNA_00124 | hypothetical protein | 1,411410627 | 3,04E-12 | 5,14E-11 |
| 197 | CCNA_01503 | hypothetical protein | 1,404898833 | 4,76E-06 | 2,18E-05 |
| 198 | CCNA_02866 | hypothetical protein | 1,404055383 | 3,15E-08 | 2,48E-07 |
| 199 | CCNA_01368 | acyl-CoA dehydrogenase | 1,399514325 | 2,24E-14 | 5,39E-13 |
| 200 | CCNA_02867 | phage tail length tape measure-related protein | 1,394808601 | 5,88E-08 | 4,38E-07 |
| 201 | CCNA_02926 | hypothetical protein | 1,394323198 | 3,26E-10 | 3,71E-09 |
| 202 | CCNA_00726 | acyl-CoA dehydrogenase | 1,392053032 | 4,82E-09 | 4,55E-08 |
| 203 | CCNA_00248 | sensor histidine protein kinase | 1,391982398 | 1,76E-12 | 3,05E-11 |
| 204 | CCNA_01595 | hypothetical protein | 1,390138919 | 6,32E-07 | 3,64E-06 |
| 205 | CCNA_03154 | PAS-family sensor histidine kinase/receiver protein | 1,386287666 | 9,00E-08 | 6,32E-07 |
| 206 | CCNA_02864 | phage protein | 1,386178273 | 6,80E-08 | 4,95E-07 |
| 207 | CCNA_02319 | hypothetical protein | 1,380521327 | 9,13E-09 | 8,09E-08 |
| 208 | CCNA_01470 | cytochrome cbb3 oxidase, diheme subunit, membrane-bound ccoP | 1,378990091 | 1,99E-08 | 1,64E-07 |
| 209 | CCNA_02701 | xanthine dehydrogenase large subunit | 1,376745831 | 7,65E-12 | 1,21E-10 |
| 210 | CCNA_01007 | hypothetical protein | 1,375798694 | 3,01E-06 | 1,46E-05 |
| 211 | CCNA_02762 | DNA repair protein radC | 1,375632807 | 2,30E-11 | 3,32E-10 |
| 212 | CCNA_00789 | hypoxia transcriptional regulator FixK | 1,37371203 | 4,68E-06 | 2,14E-05 |
| 213 | CCNA_03349 | hypothetical protein | 1,370256295 | 5,46E-10 | 6,03E-09 |
| 214 | CCNA_02967 | PAS-family sensor histidine kinase | 1,369559029 | 8,53E-14 | 1,92E-12 |
| 215 | CCNA_01925 | coniferyl aldehyde dehydrogenase | 1,368659114 | 5,15E-18 | 2,08E-16 |
| 216 | CCNA_01856 | ferrichrome-iron receptor | 1,36668478 | 3,82E-17 | 1,30E-15 |
| 217 | CCNA_02238 | hypothetical protein | 1,366124731 | 1,53E-06 | 8,05E-06 |
| ***218*** | ***CCNA_00628*** | ***chemotaxis protein cheY*** | ***1,366076615*** | ***3,84E-11*** | ***5,29E-10*** |
| 219 | CCNA_02652 | cyclohexanone monooxygenase | 1,360002697 | 8,68E-14 | 1,94E-12 |
| 220 | CCNA_03450 | hypothetical protein | 1,358948068 | 2,50E-11 | 3,56E-10 |
| 221 | CCNA_02674 | transposase | 1,352444488 | 0,018815889 | 0,036884247 |
| *222* | *CCNA_03348* | *hypothetical protein* | *1,350912962* | *2,10E-10* | *2,50E-09* |

| | | | | | |
|---|---|---|---|---|---|
| 223 | CCNA_00403 | short chain dehydrogenase | 1,349374714 | 3,32E-14 | 7,85E-13 |
| 224 | CCNA_00433 | type 1 capsular polysaccharide biosynthesis protein J | 1,347019778 | 0,000813168 | 0,002276869 |
| 225 | CCNA_01585 | TonB-dependent receptor | 1,345544605 | 3,96E-06 | 1,85E-05 |
| 226 | CCNA_01051 | TonB-dependent receptor | 1,337923873 | 7,33E-20 | 3,57E-18 |
| 227 | CCNA_03668 | hypothetical protein | 1,334221966 | 7,39E-08 | 5,32E-07 |
| 228 | CCNA_03125 | melibiose carrier protein | 1,330193597 | 4,34E-12 | 7,17E-11 |
| 229 | CCNA_02571 | transporter | 1,327729379 | 1,60E-11 | 2,39E-10 |
| 230 | CCNA_00712 | hypothetical protein | 1,324328379 | 3,41E-09 | 3,29E-08 |
| 231 | CCNA_01235 | transcriptional regulatory protein | 1,323469359 | 3,40E-12 | 5,71E-11 |
| 232 | CCNA_03574 | TonB-dependent receptor | 1,316188395 | 7,29E-18 | 2,89E-16 |
| *233* | *CCNA_00247* | *two-component receiver protein SpdR* | *1,31572471* | *5,40E-15* | *1,47E-13* |
| 234 | CCNA_02840 | transcriptional regulator MecR-family | 1,314834865 | 3,34E-13 | 6,66E-12 |
| 235 | CCNA_00714 | hypothetical protein | 1,313979676 | 1,40E-14 | 3,49E-13 |
| 236 | CCNA_00417 | hypothetical protein | 1,304657403 | 8,93E-06 | 3,89E-05 |
| 237 | CCNA_03192 | transposase | 1,303314545 | 7,59E-11 | 1,00E-09 |
| 238 | CCNA_01384 | amidase family protein | 1,30027161 | 9,71E-15 | 2,50E-13 |
| 239 | CCNA_00405 | short chain dehydrogenase | 1,295635606 | 3,26E-10 | 3,71E-09 |
| 240 | CCNA_00022 | putative aerobic-type carbon monoxide dehydrogenase, small subunit CoxS | 1,292936372 | 2,68E-09 | 2,64E-08 |
| *241* | *CCNA_00783* | *secreted protease precursor sapA* | *1,291648348* | *1,18E-10* | *1,50E-09* |
| 242 | CCNA_02696 | glutamyl-tRNA(Gln) amidotransferase subunit A | 1,28238783 | 4,92E-10 | 5,47E-09 |
| 243 | CCNA_03762 | hypothetical protein | 1,282173732 | 1,11E-05 | 4,74E-05 |
| 244 | CCNA_03172 | 3-oxoacyl-(acyl-carrier protein) reductase | 1,278233171 | 8,51E-09 | 7,60E-08 |
| 245 | CCNA_02978 | two-component sensor histidine kinase | 1,277175179 | 2,10E-10 | 2,50E-09 |
| 246 | CCNA_02205 | hypothetical protein | 1,276867424 | 3,00E-09 | 2,93E-08 |
| 247 | CCNA_01587 | ABC transporter permease protein | 1,276707061 | 0,02136131 | 0,041087979 |
| 248 | CCNA_03293 | enoyl-CoA hydratase/delta(3)-cis-delta(2)-trans-enoyl-CoA isomerase/3-hydroxyacyl-CoA dehydrogenase/ | 1,275028837 | 1,97E-13 | 4,07E-12 |
| *249* | *CCNA_03092* | *cytochrome P450 IVA5* | *1,272346697* | *1,55E-10* | *1,92E-09* |
| 250 | CCNA_03459 | methyl-accepting chemotaxis protein | 1,268896459 | 7,94E-08 | 5,68E-07 |
| 251 | CCNA_02542 | hypothetical protein | 1,266622797 | 9,69E-10 | 1,02E-08 |
| 252 | CCNA_02212 | antibiotic induced protein, Drp35 | 1,26573802 | 1,64E-10 | 2,01E-09 |
| *253* | *CCNA_01094* | *hypothetical protein* | *1,263346738* | *6,76E-06* | *3,00E-05* |
| 254 | CCNA_02471 | cobalt-zinc-cadmium resistance protein czcC | 1,26276974 | 5,99E-09 | 5,56E-08 |
| *255* | *CCNA_00021* | *putative aerobic-type carbon monoxide dehydrogenase, large subunit CoxL* | *1,260223069* | *2,01E-10* | *2,40E-09* |
| 256 | CCNA_00873 | amidophosphoribosyltransferase family protein | 1,251008593 | 4,14E-08 | 3,20E-07 |

| | | | | | |
|---|---|---|---|---|---|
| 257 | CCNA_01472 | cation pump-linked membrane protein fixH | 1,250937309 | 3,33E-09 | 3,22E-08 |
| 258 | CCNA_03087 | H-NOX family heme binding protein | 1,250419524 | 3,01E-08 | 2,39E-07 |
| 259 | CCNA_01511 | sterol desaturase family protein | 1,243383486 | 2,34E-10 | 2,74E-09 |
| 260 | CCNA_03193 | lactoylglutathione lyase | 1,238848888 | 1,63E-12 | 2,88E-11 |
| 261 | CCNA_01469 | cytochrome cbb3 oxidase, subunit ccoQ | 1,23688235 | 8,19E-06 | 3,59E-05 |
| 262 | CCNA_02553 | ATP-dependent clp protease ATP-binding subunit ClpA | 1,235230782 | 1,47E-08 | 1,25E-07 |
| 263 | CCNA_03966 | hypothetical protein | 1,233017721 | 1,16E-09 | 1,21E-08 |
| 264 | CCNA_02697 | hypothetical protein | 1,23162994 | 1,49E-08 | 1,26E-07 |
| 265 | CCNA_00282 | conserved cell surface protein | 1,231147326 | 1,68E-12 | 2,94E-11 |
| 266 | CCNA_01471 | polyferredoxin protein fixG | 1,220721348 | 1,46E-07 | 9,82E-07 |
| 267 | CCNA_02995 | glutathione S-transferase | 1,219404038 | 8,33E-12 | 1,31E-10 |
| 268 | CCNA_02695 | hypothetical protein | 1,215292525 | 5,66E-06 | 2,55E-05 |
| 269 | CCNA_02552 | ATP-dependent Clp protease adaptor protein ClpS | 1,214470602 | 5,01E-08 | 3,77E-07 |
| 270 | CCNA_00424 | aldehyde dehydrogenase | 1,213964901 | 5,94E-09 | 5,53E-08 |
| 271 | CCNA_03244 | transcriptional regulator | 1,207000492 | 3,53E-07 | 2,12E-06 |
| 272 | CCNA_02616 | 4-hydroxyphenylpyruvate dioxygenase | 1,203938052 | 2,16E-06 | 1,09E-05 |
| 273 | CCNA_02805 | hypothetical protein | 1,200784274 | 3,81E-18 | 1,56E-16 |
| 274 | CCNA_03368 | CheE protein | 1,200391655 | 3,84E-11 | 5,29E-10 |
| 275 | CCNA_00623 | PAS-family sensor histidine kinase | 1,197483971 | 6,37E-08 | 4,68E-07 |
| 276 | CCNA_02653 | sensory transduction protein kinase | 1,194927752 | 8,66E-08 | 6,12E-07 |
| 277 | CCNA_00662 | hypothetical protein | 1,192490885 | 1,17E-09 | 1,22E-08 |
| 278 | CCNA_01022 | tryptophan halogenase superfamily protein | 1,190508788 | 4,85E-12 | 7,94E-11 |
| 279 | CCNA_00275 | transposase | 1,189785901 | 1,45E-06 | 7,69E-06 |
| 280 | CCNA_03603 | beta-lactamase family protein | 1,186852511 | 1,20E-14 | 3,01E-13 |
| 281 | CCNA_R0176 | small non-coding RNA | 1,181396208 | 3,44E-05 | 0,000133079 |
| 282 | CCNA_00339 | transporter | 1,176606242 | 1,61E-14 | 3,95E-13 |
| 283 | CCNA_03484 | hypothetical protein | 1,168811497 | 9,47E-11 | 1,22E-09 |
| 284 | CCNA_02700 | XdhC protein (assists in molybdopterin insertion into xanthine dehydrogenase) | 1,168336842 | 1,02E-10 | 1,31E-09 |
| 285 | CCNA_01369 | acyl-CoA dehydrogenase, short-chain specific | 1,163537538 | 1,28E-12 | 2,30E-11 |
| 286 | CCNA_03927 | hypothetical protein | 1,163439808 | 6,24E-12 | 1,00E-10 |
| 287 | CCNA_03445 | aminopeptidase | 1,162924194 | 1,18E-14 | 2,99E-13 |
| 288 | CCNA_03085 | nitropropane dioxygenase/trans-enoyl-CoA reductase family | 1,162028812 | 2,13E-09 | 2,13E-08 |
| 289 | CCNA_03568 | LemA protein | 1,159205281 | 2,61E-06 | 1,29E-05 |
| 290 | CCNA_00853 | 3-isopropylmalate dehydrogenase | 1,158160194 | 0,019729994 | 0,038397093 |
| *291* | *CCNA_00338* | *TonB-dependent receptor* | *1,156407908* | *1,09E-11* | *1,67E-10* |
| 292 | CCNA_00570 | transcriptional regulator, GntR family | 1,156296891 | 1,19E-06 | 6,47E-06 |

| | | | | | |
|---|---|---|---|---|---|
| 293 | CCNA_03675 | PAS-family sensor histidine kinase | 1,153065245 | 7,69E-09 | 6,97E-08 |
| 294 | CCNA_01052 | SapC family protein | 1,15283234 | 1,30E-08 | 1,11E-07 |
| 295 | CCNA_02105 | hypothetical protein | 1,146463374 | 5,04E-09 | 4,75E-08 |
| 296 | CCNA_02886 | translation initiation inhibitor | 1,144170694 | 5,76E-11 | 7,74E-10 |
| 297 | CCNA_03403 | hypothetical protein | 1,143848047 | 7,26E-07 | 4,11E-06 |
| 298 | CCNA_00710 | hypothetical protein | 1,142735419 | 1,53E-13 | 3,26E-12 |
| 299 | CCNA_00434 | hypothetical protein | 1,142420359 | 0,000718952 | 0,002039886 |
| 300 | CCNA_03243 | aldehyde dehydrogenase | 1,140487288 | 1,85E-07 | 1,21E-06 |
| 301 | CCNA_02410 | alpha-methylacyl-CoA racemase | 1,140281323 | 4,62E-08 | 3,50E-07 |
| 302 | CCNA_01853 | hypothetical protein | 1,139868058 | 8,53E-08 | 6,05E-07 |
| 303 | CCNA_02216 | ABM family monooxygenase | 1,13751218 | 3,09E-12 | 5,20E-11 |
| 304 | CCNA_02215 | acyl-CoA dehydrogenase | 1,137139438 | 7,72E-14 | 1,75E-12 |
| 305 | CCNA_03437 | hypothetical protein | 1,135586573 | 1,07E-08 | 9,25E-08 |
| 306 | CCNA_00585 | hypothetical protein | 1,134892733 | 1,10E-12 | 2,01E-11 |
| 307 | CCNA_01844 | DNA integration/recombination/inversion protein | 1,134561438 | 9,56E-05 | 0,00033505 |
| 308 | CCNA_00665 | GAF-family sensor histidine kinase | 1,132763918 | 7,99E-11 | 1,05E-09 |
| 309 | CCNA_02791 | FecR family protein | 1,132424549 | 1,29E-10 | 1,62E-09 |
| 310 | CCNA_03241 | glutamate-1-semialdehyde 2,1-aminomutase | 1,13027181 | 1,27E-09 | 1,32E-08 |
| 311 | CCNA_03227 | TonB-dependent receptor | 1,129344654 | 1,68E-14 | 4,10E-13 |
| *312* | *CCNA_01767* | *cytochrome C family protein* | *1,127732737* | *2,08E-06* | *1,06E-05* |
| 313 | CCNA_01754 | transglycosylase associated protein | 1,125716222 | 1,41E-05 | 5,86E-05 |
| 314 | CCNA_02128 | acyl-CoA dehydrogenase | 1,12435322 | 8,51E-10 | 9,08E-09 |
| 315 | CCNA_00839 | glucan 1,4-beta-glucosidase | 1,122176667 | 2,78E-12 | 4,74E-11 |
| *316* | *CCNA_03795* | *tellurium resistance protein terB* | *1,120825394* | *8,77E-06* | *3,83E-05* |
| 317 | CCNA_00480 | bacteriophage P4 integrase | 1,120688718 | 2,48E-08 | 1,99E-07 |
| 318 | CCNA_03997 | amelogenin/CpxP-related protein | 1,119100457 | 1,33E-06 | 7,14E-06 |
| 319 | CCNA_02472 | cobalt-zinc-cadmium resistance protein czcB | 1,116886615 | 2,06E-08 | 1,70E-07 |
| 320 | CCNA_03485 | membrane protease family, stomatin/prohibitin-like protein | 1,115629003 | 5,82E-08 | 4,35E-07 |
| 321 | CCNA_01383 | hypothetical protein | 1,115241809 | 7,11E-10 | 7,72E-09 |
| *322* | *CCNA_03710* | *nitrogen regulatory EIIA_Ntr protein* | *1,114673013* | *5,69E-10* | *6,27E-09* |
| 323 | CCNA_01985 | ABC transporter ATP-binding protein | 1,11171358 | 1,42E-10 | 1,78E-09 |
| 324 | CCNA_03011 | outer membrane lipoprotein Blc | 1,110121091 | 3,15E-11 | 4,46E-10 |
| 325 | CCNA_00842 | hypothetical protein | 1,104348146 | 6,22E-09 | 5,73E-08 |
| ***326*** | ***CCNA_02816*** | ***hypothetical protein*** | ***1,102074315*** | ***8,64E-12*** | ***1,34E-10*** |
| 327 | CCNA_01891 | enoyl-CoA hydratase/carnithine racemase | 1,100347608 | 4,46E-06 | 2,06E-05 |
| *328* | *CCNA_01766* | *hypothetical protein* | *1,099103606* | *9,87E-11* | *1,27E-09* |
| 329 | CCNA_01927 | enoyl-CoA hydratase | 1,090909324 | 1,57E-12 | 2,79E-11 |
| 330 | CCNA_00818 | glutathione S-transferase | 1,087501421 | 7,34E-07 | 4,15E-06 |

| | | | | | |
|---|---|---|---|---|---|
| ***331*** | ***CCNA_03911*** | ***hypothetical protein*** | ***1,085780867*** | ***3,51E-07*** | ***2,11E-06*** |
| 332 | CCNA_02581 | hypothetical protein | 1,084372835 | 1,08E-07 | 7,52E-07 |
| 333 | CCNA_02931 | flgE-related flagellar hook protein | 1,080712397 | 3,56E-06 | 1,68E-05 |
| 334 | CCNA_00991 | nodulation protein N | 1,080638395 | 1,02E-08 | 8,94E-08 |
| 335 | CCNA_01462 | lactate 2-monooxygenase | 1,079012952 | 1,06E-08 | 9,22E-08 |
| 336 | CCNA_03280 | pyruvate ferredoxin/flavodoxin oxidoreductase family protein | 1,077987102 | 2,63E-06 | 1,30E-05 |
| 337 | CCNA_03532 | transporter | 1,077897887 | 2,53E-09 | 2,50E-08 |
| 338 | CCNA_02189 | peptidase, C13 family | 1,074267315 | 6,54E-11 | 8,74E-10 |
| 339 | CCNA_03271 | methionine gamma-lyase | 1,07159329 | 1,84E-06 | 9,49E-06 |
| 340 | CCNA_02564 | acyl-CoA dehydrogenase, short-chain specific | 1,070587009 | 3,19E-10 | 3,65E-09 |
| 341 | CCNA_02813 | transposase | 1,069788725 | 0,005637873 | 0,012595711 |
| 342 | CCNA_02456 | ribonuclease III | 1,067391382 | 1,11E-08 | 9,53E-08 |
| 343 | CCNA_03181 | alcohol dehydrogenase | 1,062860926 | 9,48E-15 | 2,46E-13 |
| 344 | CCNA_03709 | transposase | 1,062058619 | 0,007568706 | 0,016411032 |
| 345 | CCNA_00734 | GumN superfamily protein | 1,060324288 | 1,28E-08 | 1,09E-07 |
| 346 | CCNA_01230 | sulfatase family protein | 1,058552567 | 6,94E-12 | 1,11E-10 |
| 347 | CCNA_00822 | hypothetical protein | 1,05550664 | 6,95E-09 | 6,35E-08 |
| 348 | CCNA_02246 | division plane positioning ATPase mipZ | 1,055245769 | 1,31E-06 | 7,04E-06 |
| 349 | CCNA_02563 | acyl-CoA dehydrogenase | 1,054674235 | 3,80E-11 | 5,26E-10 |
| 350 | CCNA_02806 | nickel-cobalt-cadmium resistance protein nccC | 1,054350344 | 1,72E-06 | 8,93E-06 |
| 351 | CCNA_00788 | transporter, major facilitator superfamily | 1,053695098 | 3,61E-08 | 2,83E-07 |
| 352 | CCNA_02687 | serine-pyruvate aminotransferase | 1,049380365 | 6,08E-08 | 4,52E-07 |
| 353 | CCNA_01510 | hypothetical protein | 1,047985365 | 5,26E-07 | 3,08E-06 |
| 354 | CCNA_R0143 | small non-coding RNA | 1,047323852 | 0,003103207 | 0,007411792 |
| 355 | CCNA_01038 | glyoxalase family protein | 1,040662252 | 2,51E-09 | 2,48E-08 |
| *356* | *CCNA_02815* | *ice nucleation protein* | *1,039856696* | *1,22E-10* | *1,54E-09* |
| 357 | CCNA_02310 | cellulose 1,4-beta-cellobiosidase | 1,039165991 | 1,56E-05 | 6,41E-05 |
| 358 | CCNA_02614 | fumarylacetoacetase | 1,035515735 | 1,54E-10 | 1,91E-09 |
| 359 | CCNA_00790 | hypoxia negative feedback regulator FixT | 1,03521265 | 0,00096903 | 0,002650911 |
| 360 | CCNA_00079 | transcriptional regulator, MerR family | 1,033351644 | 1,48E-08 | 1,25E-07 |
| 361 | CCNA_01425 | H+ translocating pyrophosphatase | 1,031268451 | 2,06E-13 | 4,22E-12 |
| 362 | CCNA_03456 | lipase | 1,029583281 | 9,80E-06 | 4,23E-05 |
| 363 | CCNA_01458 | hypothetical protein | 1,02761446 | 3,54E-05 | 0,000136381 |
| 364 | CCNA_00983 | hybrid sensor histidine kinase/receiver protein | 1,026256388 | 8,70E-09 | 7,76E-08 |
| 365 | CCNA_00816 | hypothetical protein | 1,025364234 | 2,51E-07 | 1,60E-06 |
| 366 | CCNA_03580 | hypothetical protein | 1,025066318 | 5,88E-08 | 4,38E-07 |
| 367 | CCNA_02936 | glutathione S-transferase | 1,02237773 | 1,99E-11 | 2,89E-10 |
| 368 | CCNA_02924 | cytosolic protein | 1,020890426 | 4,49E-05 | 0,000169883 |

| | | | | | |
|---|---|---|---|---|---|
| 369 | CCNA_03743 | beta-lactamase family protein | 1,020557074 | 3,36E-11 | 4,71E-10 |
| 370 | CCNA_03418 | thioesterase superfamily protein | 1,019424914 | 3,20E-06 | 1,54E-05 |
| 371 | CCNA_02065 | acylamino-acid-releasing enzyme | 1,017841314 | 1,64E-12 | 2,89E-11 |
| 372 | CCNA_02927 | short chain dehydrogenase | 1,017701103 | 9,92E-07 | 5,51E-06 |
| 373 | CCNA_00076 | hypothetical protein | 1,017439679 | 1,98E-10 | 2,38E-09 |
| 374 | CCNA_02615 | homogentisate 1,2-dioxygenase | 1,016385339 | 4,12E-06 | 1,92E-05 |
| 375 | CCNA_00358 | delta 3,5-delta2,4-dienoyl-CoA isomerase precursor | 1,014229427 | 3,21E-08 | 2,52E-07 |
| *376* | *CCNA_03396* | *trypsin-like serine protease, typically periplasmic, contains C-terminal PDZ domain* | *1,012081704* | *4,99E-07* | *2,93E-06* |
| 377 | CCNA_03295 | PAS-family sensor histidine kinase | 1,011657456 | 9,06E-06 | 3,94E-05 |
| 378 | CCNA_03489 | nitropropane dioxygenase/trans-enoyl-CoA reductase family | 1,00987616 | 2,54E-06 | 1,26E-05 |
| 379 | CCNA_00484 | hypothetical protein | 1,009459635 | 1,21E-07 | 8,32E-07 |
| 380 | CCNA_01386 | hypothetical protein | 1,009056475 | 7,07E-07 | 4,02E-06 |
| 381 | CCNA_02725 | choline dehydrogenase | 1,008620229 | 5,96E-11 | 7,99E-10 |
| 382 | CCNA_02116 | putative capsule polysaccharide biosynthesis protein | 1,007752063 | 1,98E-07 | 1,29E-06 |
| *383* | *CCNA_02790* | *RNA polymerase ECF-type sigma factor* | *1,005700557* | *7,82E-07* | *4,42E-06* |
| 384 | CCNA_03569 | hypothetical protein | 1,004128441 | 1,59E-05 | 6,52E-05 |
| 385 | CCNA_01568 | Na(+)/H(+) antiporter | 1,001185833 | 1,24E-10 | 1,57E-09 |

Genes highlighted in **bold** were found in RNA-Seq_WT (Table S3). Genes highlighted in yellow were found in RNA-Seq_Δ*kdpE* (Table S4). Genes highlighted in *italics* were found in ChIP-Seq_WT (Table S5 or S6).

**Table S3. Transcriptome in low $K^+$ concentrations determined by RNA-seq.** (B) genes with expression downregulated in the WT cells grown in M2G-K 0.025 mM $K^+$ compared to WT cells grown in M2G-K 0.5 mM $K^+$.

| Top hit | Gene ID | Description | $log_2$ (FC WT Low K/ WT High K) | P-value | P-adj |
|---|---|---|---|---|---|
| 1 | CCNA_R0158 | small non-coding RNA | -2,266613584 | 6,00E-05 | 0,000220547 |
| 2 | CCNA_02277 | hemin receptor | -2,121748086 | 9,72E-22 | 6,15E-20 |
| 3 | CCNA_03022 | transporter | -2,09040474 | 5,93E-17 | 1,95E-15 |
| 4 | CCNA_02274 | EF-Hand domain protein | -2,063516921 | 2,78E-19 | 1,31E-17 |
| 5 | CCNA_00722 | chaperonin GroES | -2,044054673 | 4,55E-23 | 3,28E-21 |
| 6 | CCNA_03159 | sulfite reductase (NADPH) flavoprotein alpha-component | -1,972808766 | 3,72E-16 | 1,09E-14 |
| 7 | CCNA_03158 | iron-sulfur cluster assembly/repair protein ApbE | -1,950789808 | 3,76E-14 | 8,84E-13 |
| 8 | CCNA_02275 | hypothetical protein | -1,903004921 | 8,08E-15 | 2,14E-13 |
| 9 | CCNA_01353 | 3-phytase/6-phytase | -1,893289575 | 3,17E-13 | 6,39E-12 |
| 10 | CCNA_00721 | chaperonin GroEL | -1,859769013 | 9,87E-24 | 7,99E-22 |
| 11 | CCNA_R0211 | small non-coding RNA | -1,812733571 | 8,31E-11 | 1,09E-09 |
| ***12*** | ***CCNA_01304*** | ***hypothetical protein*** | ***-1,783427154*** | ***3,64E-29*** | ***4,91E-27*** |

| | | | | | |
|---|---|---|---|---|---|
| 13 | CCNA_03157 | hypothetical protein | -1,779345454 | 7,73E-12 | 1,22E-10 |
| 14 | CCNA_00027 | 2OG-Fe(II) oxygenase | -1,775445379 | 2,93E-18 | 1,22E-16 |
| *15* | *CCNA_00028* | *TonB-dependent receptor* | *-1,766133042* | *6,29E-13* | *1,22E-11* |
| 16 | CCNA_01516 | NAD(P)H-dependent quinone reductase | -1,757540016 | 2,25E-17 | 7,92E-16 |
| 17 | CCNA_03021 | hypothetical protein | -1,753123402 | 7,43E-13 | 1,42E-11 |
| 18 | CCNA_03023 | TonB-dependent receptor | -1,749446031 | 9,72E-13 | 1,80E-11 |
| 19 | CCNA_01310 | SSU ribosomal protein S19P | -1,723126984 | 1,06E-20 | 5,77E-19 |
| 20 | CCNA_01309 | LSU ribosomal protein L2P | -1,696268906 | 1,39E-20 | 7,21E-19 |
| 21 | CCNA_R0083 | 5S RNA | -1,683622733 | 7,43E-19 | 3,38E-17 |
| 22 | CCNA_02273 | glutathione peroxidase | -1,682694342 | 1,19E-15 | 3,38E-14 |
| ***23*** | ***CCNA_01308*** | ***LSU ribosomal protein L23P*** | ***-1,656676537*** | ***2,68E-19*** | ***1,27E-17*** |
| 24 | CCNA_01311 | LSU ribosomal protein L22P | -1,609201754 | 2,34E-17 | 8,15E-16 |
| 25 | CCNA_00084 | phosphoribosylaminoimidazolecarboxamide formyltransferase/IMP cyclohydrolase | -1,605639172 | 4,86E-17 | 1,61E-15 |
| 26 | CCNA_R0082 | tRNA Met | -1,593788998 | 3,60E-15 | 9,91E-14 |
| 27 | CCNA_R0068 | tRNA Ile | -1,59155819 | 3,00E-13 | 6,06E-12 |
| 28 | CCNA_02179 | long-chain fatty acid transport protein precursor | -1,589897209 | 1,34E-17 | 4,92E-16 |
| 29 | CCNA_R0132 | small non-coding RNA | -1,558045729 | 0,000125059 | 0,000426994 |
| 30 | CCNA_03322 | D-3-phosphoglycerate dehydrogenase | -1,55755912 | 9,96E-17 | 3,17E-15 |
| ***31*** | ***CCNA_00808*** | ***LSU ribosomal protein L34P*** | ***-1,545856364*** | ***8,29E-27*** | ***9,31E-25*** |
| ***32*** | ***CCNA_00320*** | ***LSU ribosomal protein L27P*** | ***-1,539918441*** | ***1,83E-27*** | ***2,39E-25*** |
| ***33*** | ***CCNA_R0048*** | ***tRNA Leu*** | ***-1,538133798*** | ***9,38E-18*** | ***3,55E-16*** |
| 34 | CCNA_01312 | SSU ribosomal protein S3P | -1,529651858 | 7,69E-18 | 3,02E-16 |
| 35 | CCNA_R0012 | small non-coding RNA | -1,526636243 | 0,00244696 | 0,006007525 |
| 36 | CCNA_03156 | hypothetical protein | -1,508585446 | 7,08E-09 | 6,46E-08 |
| 37 | CCNA_01792 | LSU ribosomal protein L32P | -1,50663602 | 1,84E-23 | 1,43E-21 |
| ***38*** | ***CCNA_01307*** | ***LSU ribosomal protein L1E-L4P*** | ***-1,501742185*** | ***2,91E-12*** | ***4,95E-11*** |
| 39 | CCNA_02272 | hypothetical protein | -1,473999915 | 8,64E-13 | 1,63E-11 |
| 40 | CCNA_00409 | hypothetical protein | -1,453052929 | 1,13E-09 | 1,19E-08 |
| 41 | CCNA_R0097 | small non-coding RNA | -1,450927201 | 0,0010325 | 0,002796181 |
| ***42*** | ***CCNA_01306*** | ***LSU ribosomal protein L3P*** | ***-1,450007734*** | ***4,91E-13*** | ***9,56E-12*** |
| 43 | CCNA_03307 | hypothetical protein | -1,448604402 | 3,42E-16 | 1,01E-14 |
| 44 | CCNA_01741 | SSU ribosomal protein S6P | -1,446386473 | 9,98E-13 | 1,84E-11 |
| 45 | CCNA_03196 | hypothetical protein | -1,437674855 | 0,00072422 | 0,002051958 |
| 46 | CCNA_02693 | iron-regulated protein frpC | -1,434631195 | 2,03E-21 | 1,21E-19 |
| 47 | CCNA_03767 | SSU ribosomal protein S16P | -1,431651002 | 3,22E-13 | 6,45E-12 |
| 48 | CCNA_03321 | hypothetical protein | -1,422461086 | 6,89E-16 | 1,99E-14 |
| 49 | CCNA_01739 | LSU ribosomal protein L9P | -1,40673832 | 1,12E-13 | 2,46E-12 |
| 50 | CCNA_01313 | LSU ribosomal protein L16P | -1,406490315 | 2,53E-18 | 1,08E-16 |
| ***51*** | ***CCNA_01305*** | ***SSU ribosomal protein S10P*** | ***-1,392512189*** | ***2,68E-13*** | ***5,45E-12*** |
| 52 | CCNA_02596 | SSU ribosomal protein S4P | -1,390324916 | 1,19E-14 | 3,01E-13 |

| | | | | | |
|---|---|---|---|---|---|
| 53 | CCNA_01350 | transposase | -1,389661875 | 6,36E-08 | 4,68E-07 |
| ***54*** | ***CCNA_03558*** | ***ATP synthase subunit epsilon*** | ***-1,385639994*** | ***9,62E-16*** | ***2,74E-14*** |
| 55 | CCNA_03904 | hypothetical protein | -1,381595611 | 6,89E-09 | 6,31E-08 |
| 56 | CCNA_03559 | hypothetical protein | -1,367783501 | 1,30E-13 | 2,82E-12 |
| 57 | CCNA_R0067 | tRNA Ala | -1,361395621 | 3,30E-06 | 1,58E-05 |
| ***58*** | ***CCNA_R0008*** | ***tRNA-Arg*** | ***-1,358985178*** | ***9,04E-10*** | ***9,62E-09*** |
| 59 | CCNA_R0065 | 5S RNA | -1,354830008 | 1,27E-17 | 4,72E-16 |
| *60* | *CCNA_03430* | *LSU ribosomal protein L36P* | *-1,350112858* | *3,80E-21* | *2,20E-19* |
| 61 | CCNA_R0024 | tRNA Ser | -1,343668691 | 3,65E-11 | 5,10E-10 |
| 62 | CCNA_01440 | LSU ribosomal protein L13P | -1,336494039 | 2,68E-11 | 3,80E-10 |
| 63 | CCNA_03968 | hypothetical protein | -1,332661483 | 0,019788387 | 0,038473721 |
| 64 | CCNA_01314 | LSU ribosomal protein L29P | -1,326700893 | 9,14E-19 | 4,11E-17 |
| **65** | **CCNA_R0090** | **tRNA Leu** | **-1,325694799** | **3,10E-09** | **3,02E-08** |
| 66 | CCNA_00530 | LSU ribosomal protein L10P | -1,320883079 | 3,24E-11 | 4,56E-10 |
| 67 | CCNA_03155 | transporter | -1,318773867 | 1,29E-07 | 8,84E-07 |
| *68* | *CCNA_03384* | *LSU ribosomal protein L31P* | *-1,317417219* | *2,35E-18* | *1,01E-16* |
| 69 | CCNA_03303 | protein translation elongation factor Tu (EF-TU) | -1,314010055 | 8,90E-15 | 2,32E-13 |
| 70 | CCNA_03766 | 16S rRNA processing protein rimM | -1,309170006 | 1,70E-13 | 3,56E-12 |
| 71 | CCNA_00532 | hypothetical protein | -1,301753238 | 2,12E-15 | 5,92E-14 |
| 72 | CCNA_R0026 | tRNA Gly | -1,283186615 | 1,53E-08 | 1,29E-07 |
| 73 | CCNA_R0030 | tRNA Tyr | -1,27995233 | 3,28E-16 | 9,75E-15 |
| 74 | CCNA_01332 | LSU ribosomal protein L17P | -1,265181575 | 3,07E-10 | 3,53E-09 |
| 75 | CCNA_03305 | SSU ribosomal protein S7P | -1,262447243 | 3,58E-13 | 7,06E-12 |
| 76 | CCNA_00289 | hypothetical protein | -1,258868389 | 1,30E-09 | 1,35E-08 |
| ***77*** | ***CCNA_01330*** | ***DNA-directed RNA polymerase subunit alpha*** | ***-1,254221452*** | ***1,26E-12*** | ***2,29E-11*** |
| 78 | CCNA_R0162 | small non-coding RNA | -1,253752698 | 2,69E-10 | 3,12E-09 |
| 79 | CCNA_01740 | SSU ribosomal protein S18P | -1,246563564 | 1,74E-09 | 1,76E-08 |
| 80 | CCNA_01770 | nucleoside diphosphate kinase | -1,24614693 | 1,22E-11 | 1,86E-10 |
| 81 | CCNA_03428 | cytosolic protein | -1,244560566 | 3,88E-10 | 4,40E-09 |
| 82 | CCNA_01543 | chaperone protein dnaK | -1,244460752 | 4,73E-11 | 6,43E-10 |
| 83 | CCNA_03768 | signal recognition particle GTPase, SRP | -1,236760211 | 1,62E-13 | 3,41E-12 |
| 84 | CCNA_00749 | ferrous iron transport protein B | -1,235633187 | 8,97E-07 | 5,01E-06 |
| 85 | CCNA_01939 | ADP-ribosylglycohydrolase | -1,234190835 | 7,70E-09 | 6,97E-08 |
| 86 | CCNA_00887 | hypothetical protein | -1,232687774 | 4,09E-13 | 8,00E-12 |
| 87 | CCNA_03851 | imidazole glycerol phosphate synthase, glutamine amidotransferase subunit | -1,232146312 | 1,82E-11 | 2,68E-10 |
| 88 | CCNA_R0022 | tRNA Gln | -1,231606349 | 8,62E-12 | 1,34E-10 |
| 89 | CCNA_R0058 | tRNA Phe | -1,231231887 | 4,00E-15 | 1,09E-13 |
| 90 | CCNA_03901 | conserved hypothetical protein | -1,22712028 | 8,56E-11 | 1,11E-09 |
| 91 | CCNA_03807 | organic solvent resistance transport system Ttg2D protein | -1,225924378 | 8,91E-08 | 6,27E-07 |

| | | | | | |
|---|---|---|---|---|---|
| 92 | CCNA_03323 | phosphoserine aminotransferase | -1,225920478 | 3,32E-11 | 4,67E-10 |
| 93 | CCNA_03561 | ATP synthase subunit gamma | -1,225666127 | 2,47E-11 | 3,54E-10 |
| 94 | CCNA_01938 | ATP-dependent transporter sufC | -1,220529929 | 7,91E-11 | 1,04E-09 |
| 95 | CCNA_00678 | LSU ribosomal protein L11P | -1,217495323 | 1,31E-11 | 1,98E-10 |
| 96 | CCNA_03509 | zinc protease | -1,216558718 | 4,39E-10 | 4,91E-09 |
| 97 | CCNA_03560 | ATP synthase subunit beta | -1,212809158 | 5,28E-11 | 7,12E-10 |
| 98 | CCNA_00805 | GTP-binding protein YihA | -1,21029823 | 5,24E-12 | 8,54E-11 |
| 99 | CCNA_R0052 | tRNA Leu | -1,21004549 | 2,14E-10 | 2,54E-09 |
| 100 | CCNA_01802 | hypothetical protein | -1,208222265 | 5,13E-06 | 2,33E-05 |
| 101 | CCNA_02307 | arogenate dehydrogenase/Prephenate dehydrogenase | -1,199690955 | 1,04E-11 | 1,60E-10 |
| 102 | CCNA_01441 | SSU ribosomal protein S9P | -1,195661344 | 1,13E-09 | 1,19E-08 |
| 103 | CCNA_03723 | PaaI thioesterase family protein | -1,191926314 | 1,61E-07 | 1,07E-06 |
| 104 | CCNA_00637 | tetratricopeptide repeat family protein | -1,191625311 | 8,76E-14 | 1,95E-12 |
| 105 | CCNA_R0007 | tRNA-Ser | -1,186936736 | 1,94E-11 | 2,84E-10 |
| 106 | CCNA_01560 | phosphoenolpyruvate carboxylase | -1,176991781 | 2,48E-09 | 2,46E-08 |
| 107 | CCNA_R0160 | small non-coding RNA | -1,175139646 | 0,001158339 | 0,003093492 |
| *108* | *CCNA_00034* | *SSU ribosomal protein S15P* | *-1,170154689* | *4,27E-11* | *5,84E-10* |
| 109 | CCNA_01827 | adenosylcobyric acid synthase (glutamine-hydrolysing) | -1,169261562 | 5,15E-11 | 6,97E-10 |
| *110* | *CCNA_01920* | *transcription-repair coupling factor* | *-1,165293873* | *5,43E-12* | *8,82E-11* |
| 111 | CCNA_02181 | phosphatase/phosphohexomutase family protein | -1,164999483 | 2,07E-11 | 3,00E-10 |
| 112 | CCNA_01561 | hypothetical protein | -1,15996932 | 1,50E-05 | 6,21E-05 |
| 113 | CCNA_01098 | LSU ribosomal protein L35P | -1,158741619 | 2,66E-10 | 3,09E-09 |
| 114 | CCNA_00531 | LSU ribosomal protein L12P (L7/L12) | -1,158576088 | 1,34E-08 | 1,14E-07 |
| 115 | CCNA_00138 | TonB-dependent receptor | -1,156776146 | 3,39E-06 | 1,61E-05 |
| 116 | CCNA_02574 | cytosolic protein | -1,153504566 | 6,56E-08 | 4,79E-07 |
| 117 | CCNA_03304 | protein translation elongation factor G (EF-G) | -1,153495243 | 1,80E-10 | 2,18E-09 |
| 118 | CCNA_R0021 | tRNA Pro | -1,151821424 | 7,97E-09 | 7,18E-08 |
| 119 | CCNA_R0085 | tRNA Ala | -1,14998249 | 0,02652878 | 0,049715351 |
| 120 | CCNA_02324 | heat shock protein 33 hsp33 | -1,149235322 | 1,74E-11 | 2,57E-10 |
| 121 | CCNA_03352 | cytosolic protein | -1,148224232 | 1,06E-09 | 1,12E-08 |
| 122 | CCNA_00807 | ribonuclease P protein component | -1,148164023 | 1,68E-11 | 2,50E-10 |
| 123 | CCNA_R0027 | tRNA Gly | -1,146666468 | 3,33E-07 | 2,03E-06 |
| *124* | *CCNA_03406* | *SSU ribosomal protein S21P* | *-1,141407492* | *9,92E-07* | *5,51E-06* |
| *125* | *CCNA_03372* | *bacterioferritin-associated ferredoxin* | *-1,140794536* | *2,48E-10* | *2,89E-09* |
| 126 | CCNA_00085 | dienelactone hydrolase-related protein | -1,137725663 | 4,57E-11 | 6,23E-10 |
| 127 | CCNA_00677 | LSU ribosomal protein L1P | -1,136501061 | 1,34E-09 | 1,38E-08 |
| 128 | CCNA_01563 | 2-dehydro-3-deoxygluconokinase | -1,133658967 | 1,59E-09 | 1,62E-08 |
| *129* | *CCNA_01328* | *SSU ribosomal protein S13P* | *-1,130535094* | *2,26E-10* | *2,65E-09* |
| 130 | CCNA_01315 | SSU ribosomal protein S17P | -1,129807092 | 7,13E-13 | 1,37E-11 |
| 131 | CCNA_01329 | SSU ribosomal protein S11P | -1,129354409 | 3,99E-11 | 5,48E-10 |

| | | | | | |
|---|---|---|---|---|---|
| 132 | CCNA_00372 | ATP synthase C chain | -1,128492511 | 8,14E-12 | 1,28E-10 |
| 133 | CCNA_01937 | SufD protein | -1,12591442 | 4,26E-10 | 4,80E-09 |
| ***134*** | ***CCNA_R0028*** | ***tRNA Lys*** | ***-1,125723092*** | ***1,57E-10*** | ***1,94E-09*** |
| *135* | *CCNA_01921* | *TPR repeat containing protein* | *-1,122721008* | *6,79E-09* | *6,23E-08* |
| *136* | *CCNA_00042* | *cytosolic protein/LSU ribosomal protein L7AE* | *-1,120879099* | *1,71E-13* | *3,56E-12* |
| 137 | CCNA_01097 | LSU ribosomal protein L20P | -1,120525525 | 8,82E-11 | 1,14E-09 |
| 138 | CCNA_R0144 | small non-coding RNA | -1,114030841 | 0,001098232 | 0,002966253 |
| 139 | CCNA_00284 | hydrolase (HAD superfamily) | -1,113531614 | 1,15E-08 | 9,86E-08 |
| **140** | **CCNA_03306** | **SSU ribosomal protein S12P** | **-1,1116061** | **7,96E-09** | **7,18E-08** |
| 141 | CCNA_R0031 | tRNA Gly | -1,11061932 | 3,25E-07 | 1,98E-06 |
| 142 | CCNA_01316 | LSU ribosomal protein L14P | -1,109764013 | 1,86E-13 | 3,86E-12 |
| 143 | CCNA_01363 | two-component sensor histidine kinase, UrpS | -1,109526931 | 1,68E-09 | 1,71E-08 |
| 144 | CCNA_02999 | hypothetical protein | -1,107455544 | 1,80E-07 | 1,19E-06 |
| 145 | CCNA_01962 | leucyl/phenylalanyl-tRNA--protein transferase | -1,10500018 | 2,45E-08 | 1,97E-07 |
| *146* | *CCNA_00035* | *tRNA pseudouridine synthase B* | *-1,104807872* | *4,44E-12* | *7,30E-11* |
| 147 | CCNA_00011 | chaperone protein DnaJ | -1,103254627 | 2,37E-11 | 3,42E-10 |
| *148* | *CCNA_00044* | *cytosolic protein* | *-1,102563685* | *5,46E-12* | *8,84E-11* |
| *149* | *CCNA_00699* | *LSU ribosomal protein L28P* | *-1,099537861* | *7,14E-10* | *7,72E-09* |
| 150 | CCNA_01999 | protein translation Elongation factor Ts (EF-Ts) | -1,098779112 | 6,33E-09 | 5,83E-08 |
| *151* | *CCNA_R0076* | *tRNA Trp* | *-1,097946662* | *1,36E-13* | *2,94E-12* |
| 152 | CCNA_01936 | cysteine desulfurase/Selenocysteine lyase | -1,097183947 | 2,19E-10 | 2,58E-09 |
| 153 | CCNA_03476 | ribonuclease HI | -1,096753863 | 3,55E-07 | 2,13E-06 |
| 154 | CCNA_03028 | prephenate dehydratase | -1,094043915 | 2,96E-10 | 3,42E-09 |
| 155 | CCNA_01691 | 16S rRNA m(5)C 967 methyltransferase | -1,089807434 | 9,68E-13 | 1,80E-11 |
| 156 | CCNA_01546 | DEAD-box RNA helicase-like protein | -1,087752953 | 6,69E-13 | 1,29E-11 |
| 157 | CCNA_03806 | outer membrane lipoprotein | -1,087692644 | 1,60E-08 | 1,34E-07 |
| 158 | CCNA_00215 | cytosolic protein | -1,08496814 | 8,44E-10 | 9,04E-09 |
| 159 | CCNA_02000 | SSU ribosomal protein S2P | -1,084301547 | 1,77E-09 | 1,78E-08 |
| 160 | CCNA_01361 | hypothetical protein | -1,082905501 | 7,51E-10 | 8,10E-09 |
| 161 | CCNA_02325 | ornithine carbamoyltransferase | -1,082465239 | 2,29E-09 | 2,28E-08 |
| 162 | CCNA_00374 | putative ATP synthase protein I | -1,08218505 | 3,48E-13 | 6,90E-12 |
| 163 | CCNA_02042 | trigger factor, ppiase | -1,078687818 | 9,55E-09 | 8,44E-08 |
| 164 | CCNA_03164 | protein translocase subunit secA | -1,078567312 | 2,42E-10 | 2,83E-09 |
| *165* | *CCNA_R0001* | *tRNA-thr* | *-1,078382591* | *3,73E-11* | *5,18E-10* |
| 166 | CCNA_00518 | LSU ribosomal protein L25P | -1,076918617 | 1,04E-08 | 9,11E-08 |
| 167 | CCNA_01774 | phosphoribosylglycinamide formyltransferase | -1,068469973 | 3,97E-12 | 6,61E-11 |
| 168 | CCNA_01795 | enolase | -1,067095685 | 4,32E-10 | 4,86E-09 |
| 169 | CCNA_03810 | organic solvent resistance transport system ATP-binding protein | -1,064928189 | 2,52E-06 | 1,25E-05 |

| | | | | | |
|---|---|---|---|---|---|
| *170* | *CCNA_02543* | *LSU ribosomal protein L33P* | *-1,064297608* | *1,69E-10* | *2,06E-09* |
| 171 | CCNA_01325 | LSU ribosomal protein L15P | -1,063883259 | 4,28E-08 | 3,29E-07 |
| 172 | CCNA_03583 | hypothetical protein | -1,062490644 | 2,31E-07 | 1,48E-06 |
| 173 | CCNA_01801 | hypothetical protein | -1,057767464 | 6,20E-08 | 4,59E-07 |
| 174 | CCNA_00850 | cation/multidrug efflux pump acrB2 | -1,056776196 | 1,39E-12 | 2,49E-11 |
| 175 | CCNA_01254 | SsrA-binding protein | -1,048221709 | 5,73E-10 | 6,30E-09 |
| 176 | CCNA_R0003 | tRNA-Arg | -1,047930317 | 5,64E-07 | 3,28E-06 |
| 177 | CCNA_03153 | hypothetical protein | -1,046975345 | 3,95E-08 | 3,06E-07 |
| 178 | CCNA_01362 | two-component response regulator, UrpR | -1,046961368 | 3,36E-07 | 2,04E-06 |
| 179 | CCNA_01298 | protein translation Elongation factor Tu (EF-TU) | -1,04524205 | 3,66E-07 | 2,18E-06 |
| *180* | *CCNA_R0091* | *tRNA Ala* | *-1,044378004* | *8,91E-09* | *7,91E-08* |
| 181 | CCNA_00638 | heme:hemopexin-binding protein | -1,043666401 | 2,31E-07 | 1,48E-06 |
| 182 | CCNA_01562 | 4-hydroxy-2-oxoglutarate aldolase/2-dehydro-3-deoxyphosphogluconate aldolase | -1,042397206 | 9,65E-09 | 8,51E-08 |
| 183 | CCNA_00410 | acetyltransferase | -1,042002782 | 1,69E-06 | 8,78E-06 |
| 184 | CCNA_01162 | beta-glucosidase | -1,039701844 | 8,25E-08 | 5,86E-07 |
| 185 | CCNA_02572 | adenylosuccinate lyase | -1,038035854 | 1,28E-09 | 1,33E-08 |
| 186 | CCNA_03809 | organic solvent resistance transport system permease | -1,037251266 | 1,11E-07 | 7,71E-07 |
| 187 | CCNA_03168 | hypothetical protein | -1,03681032 | 5,61E-09 | 5,26E-08 |
| 188 | CCNA_00184 | hypothetical protein | -1,026963456 | 0,001690335 | 0,004314887 |
| 189 | CCNA_03907 | conserved hypothetical protein | -1,026429721 | 4,38E-10 | 4,91E-09 |
| 190 | CCNA_03108 | ChvT TonB-dependent outer membrane receptor | -1,025609737 | 2,03E-07 | 1,32E-06 |
| 191 | CCNA_03301 | cation/multidrug efflux pump acrB | -1,024462465 | 1,42E-07 | 9,59E-07 |
| 192 | CCNA_03995 | hypothetical protein | -1,024185464 | 8,78E-12 | 1,36E-10 |
| 193 | CCNA_00154 | hypothetical protein | -1,023221642 | 2,88E-06 | 1,41E-05 |
| 194 | CCNA_03152 | hypothetical protein | -1,021247763 | 2,80E-05 | 0,000109813 |
| 195 | CCNA_03300 | multidrug resistance efflux pump, periplasmic component | -1,021099783 | 1,41E-08 | 1,20E-07 |
| 196 | CCNA_01095 | phenylalanyl-tRNA synthetase subunit beta | -1,02068867 | 1,64E-09 | 1,67E-08 |
| 197 | CCNA_01969 | aspartyl-tRNA synthetase | -1,017890038 | 3,99E-09 | 3,82E-08 |
| 198 | CCNA_03481 | DNA integration/recombination/inversion protein | -1,013846903 | 1,19E-07 | 8,18E-07 |
| 199 | CCNA_02573 | proline hydroxylase | -1,012693543 | 1,64E-10 | 2,02E-09 |
| 200 | CCNA_00373 | ATP synthase A chain | -1,009779271 | 1,70E-08 | 1,41E-07 |
| 201 | CCNA_03378 | myo-inositol-1(or 4)-monophosphatase | -1,009071794 | 6,52E-10 | 7,12E-09 |
| 202 | CCNA_R0182 | small non-coding RNA | -1,00723354 | 3,90E-06 | 1,83E-05 |
| 203 | CCNA_00212 | N-carbamoylputrescine amidase | -1,002924601 | 1,90E-09 | 1,91E-08 |
| 204 | CCNA_03012 | peroxiredoxin | -1,002818642 | 1,38E-07 | 9,33E-07 |
| 205 | CCNA_01950 | bacterial peptide chain Release factor 2 (RF-2) | -1,002517566 | 1,17E-07 | 8,09E-07 |
| *206* | *CCNA_00198* | *tRNA (guanine-N1) -methyltransferase* | *-1,001445129* | *1,09E-08* | *9,44E-08* |

| | | | | | |
|---|---|---|---|---|---|
| 207 | CCNA_03167 | hypothetical protein | -1,001258472 | 2,19E-10 | 2,58E-09 |
| 208 | CCNA_02942 | hypothetical protein | -1,000364621 | 2,80E-10 | 3,24E-09 |
| 209 | CCNA_01648 | adenosylmethionine-8-amino-7-oxononanoate | -1,000025645 | 1,02E-08 | 8,94E-08 |

Genes highlighted in **bold** were found in Tn-Seq (Table S2). Genes highlighted in yellow were found in RNA-Seq_Δ*kdpE* (Table S4). Genes highlighted in *italics* were found in ChIP-Seq_WT (Table S5 or S6).

**Table S4. KdpE transcriptome in low $K^+$ concentrations determined by RNA-seq.** (A) genes with expression upregulated in the Δ*kdpE* cells grown in M2G-K 0.025 mM $K^+$ compared to WT cells grown in the same medium.

| Top hit | Gene ID | Description | $log_2$ (FC Low $K^+$ Δ*kdpE* / WT) | P-value | P-adj |
|---|---|---|---|---|---|
| *1* | *CCNA_00224* | *hypothetical protein* | *1,491748502* | *4,63E-27* | *2,52E-24* |
| *2* | *CCNA_02186* | *acetolactate synthase small subunit* | *1,306474726* | *9,05E-12* | *1,57E-09* |
| 3 | CCNA_00882 | hypothetical protein | 1,279600876 | 4,83E-10 | 5,26E-08 |
| 4 | CCNA_03098 | hypothetical protein | 1,244016525 | 1,61E-16 | 6,80E-14 |
| 5 | CCNA_00721 | chaperonin GroEL | 1,242237545 | 6,23E-13 | 1,32E-10 |
| 6 | CCNA_00913 | hypothetical protein | 1,237346479 | 1,10E-13 | 2,81E-11 |
| 7 | CCNA_00768 | hypothetical protein | 1,197172624 | 1,02E-11 | 1,69E-09 |
| 8 | CCNA_02723 | hypothetical protein | 1,166203023 | 1,53E-10 | 2,01E-08 |
| 9 | CCNA_00858 | TonB-dependent receptor | 1,158842343 | 9,20E-11 | 1,30E-08 |
| 10 | CCNA_00722 | chaperonin GroES | 1,148844671 | 8,85E-11 | 1,30E-08 |
| 11 | CCNA_02255 | lysozyme-family localization factor spmX | 1,142950391 | 3,65E-12 | 6,95E-10 |
| 12 | CCNA_00769 | transcriptional regulator, TetR family | 1,136883719 | 5,58E-11 | 8,50E-09 |
| 13 | CCNA_01551 | hypothetical protein | 1,129900675 | 1,52E-10 | 2,01E-08 |
| 14 | CCNA_02758 | hypothetical protein | 1,128675814 | 2,91E-10 | 3,58E-08 |
| 15 | CCNA_02172 | transporter | 1,120401271 | 8,21E-14 | 2,40E-11 |
| 16 | CCNA_03456 | lipase | 1,0618296 | 2,60E-08 | 2,36E-06 |
| 17 | CCNA_03271 | methionine gamma-lyase | 1,061312939 | 4,47E-09 | 4,60E-07 |
| 18 | CCNA_01700 | nucleoside permease | 1,048870804 | 7,14E-08 | 5,92E-06 |
| *19* | *CCNA_03396* | *trypsin-like serine protease, typically periplasmic, contains C-terminal PDZ domain* | *1,021382926* | *7,42E-09* | *7,24E-07* |
| 20 | CCNA_00912 | 6-aminohexanoate-dimer hydrolase | 1,002520812 | 5,28E-13 | 1,18E-10 |
| 21 | CCNA_03721 | glutamate synthase (NADPH) small chain | 0,951759426 | 6,17E-08 | 5,22E-06 |
| 22 | CCNA_03097 | aldo/keto reductase family protein | 0,928909429 | 1,69E-07 | 1,29E-05 |
| *23* | *CCNA_00338* | *TonB-dependent receptor* | *0,903361432* | *1,13E-08* | *1,08E-06* |
| 24 | CCNA_00732 | GumN superfamily protein | 0,901757647 | 7,22E-07 | 4,74E-05 |
| 25 | CCNA_02173 | ABC transporter ATP-binding protein | 0,90166574 | 3,60E-10 | 4,16E-08 |
| ***26*** | ***CCNA_02185*** | ***acetolactate synthase large subunit*** | ***0,851684778*** | ***3,21E-06*** | ***0,000174899*** |
| 27 | CCNA_00339 | transporter | 0,846247777 | 3,04E-09 | 3,22E-07 |
| 28 | CCNA_03805 | transcriptional regulator | 0,837923125 | 1,40E-07 | 1,09E-05 |

| | | | | | |
|---|---|---|---|---|---|
| 29 | CCNA_03936 | hypothetical protein | 0,833715833 | 5,40E-05 | 0,002100432 |
| 30 | CCNA_03138 | peroxidase/catalase katG | 0,822720034 | 1,89E-05 | 0,000793458 |
| ***31*** | ***CCNA_03911*** | ***hypothetical protein*** | ***0,814625888*** | ***1,55E-06*** | ***9,38E-05g*** |
| 32 | CCNA_03301 | cation/multidrug efflux pump acrB | 0,813937972 | 4,34E-06 | 0,000226565 |
| 33 | CCNA_01078 | methyltransferase | 0,79752365 | 4,58E-05 | 0,001799629 |
| 34 | CCNA_02174 | multidrug resistance efflux pump | 0,790309972 | 1,89E-07 | 1,41E-05 |
| 35 | CCNA_02341 | small heat shock protein | 0,787815331 | 5,50E-08 | 4,76E-06 |
| 36 | CCNA_00022 | putative aerobic-type carbon monoxide dehydrogenase, small subunit CoxS | 0,785655946 | 1,92E-05 | 0,000793458 |
| 37 | CCNA_01179 | phosphoadenosine phosphosulfate reductase | 0,782410072 | 2,57E-05 | 0,001040559 |
| 38 | CCNA_01312 | SSU ribosomal protein S3P | 0,77976128 | 5,26E-06 | 0,000263512 |
| 39 | CCNA_01379 | type I secretion outer membrane protein rsaFb | 0,769374448 | 1,37E-08 | 1,27E-06 |
| 40 | CCNA_03117 | cytosolic protein | 0,7672322 | 4,87E-07 | 3,32E-05 |
| 41 | CCNA_00148 | hypothetical protein | 0,763059403 | 2,75E-07 | 1,98E-05 |
| 42 | CCNA_01311 | LSU ribosomal protein L22P | 0,76068326 | 1,59E-05 | 0,000700388 |
| *43* | *CCNA_00021* | *putative aerobic-type carbon monoxide dehydrogenase, large subunit CoxL* | *0,752909155* | *1,79E-05* | *0,000774757* |
| 44 | CCNA_02856 | hypothetical protein | 0,746175383 | 2,96E-06 | 0,000163232 |
| ***45*** | ***CCNA_03558*** | ***ATP synthase subunit epsilon*** | ***0,733212235*** | ***7,36E-06*** | ***0,000346247*** |
| 46 | CCNA_01376 | glutathione S-transferase | 0,728099678 | 5,90E-07 | 3,94E-05 |
| 47 | CCNA_00064 | methyl-accepting chemotaxis protein | 0,718067175 | 0,0003104 | 0,009771188 |
| 48 | CCNA_01139 | conserved cytosolic protein | 0,716288841 | 9,60E-08 | 7,78E-06 |
| 49 | CCNA_01313 | LSU ribosomal protein L16P | 0,708753046 | 4,33E-06 | 0,000226565 |
| 50 | CCNA_01178 | oxidoreductase | 0,706030875 | 0,000275913 | 0,008831525 |
| 51 | CCNA_01314 | LSU ribosomal protein L29P | 0,702803242 | 1,32E-06 | 8,38E-05 |
| 52 | CCNA_01702 | NAD-dependent oxidoreductase | 0,701785684 | 0,000705233 | 0,018525754 |
| 53 | CCNA_03034 | hypothetical protein | 0,69619191 | 1,64E-06 | 9,74E-05 |
| 54 | CCNA_01968 | hypothetical protein | 0,695825204 | 0,000698181 | 0,018525754 |
| 55 | CCNA_03569 | hypothetical protein | 0,695499006 | 0,000317818 | 0,009922683 |
| 56 | CCNA_00077 | acyl-CoA dehydrogenase, long-chain specific | 0,679872581 | 3,10E-05 | 0,001244389 |
| 57 | CCNA_02196 | hypothetical protein | 0,679682776 | 0,000701194 | 0,018525754 |
| 58 | CCNA_00849 | outer membrane efflux protein | 0,676479274 | 3,71E-06 | 0,000198892 |
| 59 | CCNA_00711 | hypothetical protein | 0,669575122 | 1,40E-05 | 0,000641142 |
| 60 | CCNA_00076 | hypothetical protein | 0,660685727 | 5,06E-06 | 0,000257207 |
| 61 | CCNA_03810 | organic solvent resistance transport system ATP-binding protein | 0,651282884 | 0,000758549 | 0,019425723 |
| 62 | CCNA_03559 | hypothetical protein | 0,648636275 | 0,00012381 | 0,004326529 |
| 63 | CCNA_02199 | methyltransferase | 0,647582529 | 0,000883854 | 0,021908821 |
| 64 | CCNA_01172 | hypothetical protein | 0,638680329 | 1,56E-05 | 0,000700388 |
| 65 | CCNA_03570 | hypothetical protein | 0,634003049 | 0,001271768 | 0,02918682 |
| 66 | CCNA_01315 | SSU ribosomal protein S17P | 0,633173229 | 2,08E-05 | 0,000853519 |
| 67 | CCNA_00538 | methyl-accepting chemotaxis protein | 0,631547909 | 3,81E-05 | 0,001512033 |

| | | | | | |
|---|---|---|---|---|---|
| 68 | CCNA_02366 | cytoplasmic membrane maltose transporter malY | 0,630376313 | 8,82E-06 | 0,000409787 |
| 69 | CCNA_02621 | CAAX amino terminal protease family | 0,629271021 | 0,000118547 | 0,004180964 |
| 70 | CCNA_01589 | acyltransferase 3 | 0,628363282 | 0,000504374 | 0,014215232 |
| 71 | CCNA_01712 | beta-lactamase repressor | 0,627724975 | 0,000101004 | 0,00361659 |
| 72 | CCNA_00437 | methyl-accepting chemotaxis protein | 0,624720037 | 0,000446439 | 0,01328506 |
| 73 | CCNA_03049 | transporter, MFS superfamily | 0,613383199 | 6,37E-05 | 0,002452672 |
| 74 | CCNA_02845 | two-component response regulator | 0,613259443 | 5,83E-06 | 0,000284738 |
| 75 | CCNA_03568 | LemA protein | 0,609263935 | 0,002189214 | 0,045566752 |
| 76 | CCNA_01310 | SSU ribosomal protein S19P | 0,607493214 | 0,000759893 | 0,019425723 |
| 77 | CCNA_02677 | glutathione S-transferase | 0,60114495 | 0,000138797 | 0,004720325 |
| 78 | CCNA_02361 | polysaccharide biosynthesis protein celD | 0,596610061 | 1,60E-05 | 0,000700388 |
| 79 | CCNA_02987 | hypothetical protein | 0,591666252 | 0,001390836 | 0,031453898 |
| 80 | CCNA_03699 | peptidase, M16 family | 0,590103116 | 7,34E-05 | 0,002768562 |
| 81 | CCNA_01335 | transporter | 0,589771406 | 0,000483941 | 0,013756201 |
| 82 | CCNA_00459 | Na+/H+ antiporter nhaA | 0,588818046 | 0,000786237 | 0,019965183 |
| 83 | CCNA_03804 | cytosolic protein | 0,5861044 | 0,000539262 | 0,014993064 |

Rows in grey indicate genes with log$_2$ FC > 1. Genes highlighted in **bold** were found in Tn-Seq (Table S2). Genes highlighted in yellow were found in RNA-Seq_WT (Table S3). Genes highlighted in *italics* were found in ChIP-Seq_WT (Table S5 or S6).

**Table S4. KdpE transcriptome in low $K^+$ concentrations determined by RNA-seq.** (B) genes with expression downregulated in the $\Delta$*kdpE* cells grown in M2G-K 0.025 mM $K^+$ compared to WT cells grown in the same medium.

| Top hit | Gene ID | Description | log2 (FC WT Low K/ WT High K) | P-value | P-adj |
|---|---|---|---|---|---|
| 1 | CCNA_01667 | two-component response regulator kdpE | -5,900050777 | 6,15E-215 | 2,34E-211 |
| ***2*** | ***CCNA_01664*** | ***potassium-transporting ATPase B chain*** | ***-5,437104702*** | ***1,20E-162*** | ***2,28E-159*** |
| ***3*** | ***CCNA_01663*** | ***potassium-transporting ATPase A chain*** | ***-4,587220099*** | ***4,25E-97*** | ***3,24E-94*** |
| ***4*** | ***CCNA_01665*** | ***potassium-transporting ATPase C chain*** | ***-4,340624976*** | ***2,17E-135*** | ***2,07E-132*** |
| ***5*** | ***CCNA_01666*** | ***osmosensitive K+ channel histidine kinase kdpD*** | ***-3,866286502*** | ***1,48E-143*** | ***1,87E-140*** |
| 6 | CCNA_02277 | hemin receptor | -1,822485608 | 1,46E-19 | 6,97E-17 |
| 7 | CCNA_03624 | sodium bicarbonate cotransporter | -1,738888284 | 3,15E-30 | 2,00E-27 |
| 8 | CCNA_01859 | TonB-dependent receptor | -1,73850716 | 1,62E-15 | 6,17E-13 |
| 9 | CCNA_02274 | EF-Hand domain protein | -1,639766842 | 3,34E-15 | 1,16E-12 |
| 10 | CCNA_02275 | hypothetical protein | -1,600005957 | 3,90E-15 | 1,24E-12 |
| 11 | CCNA_01860 | rare lipoprotein B precursor | -1,541891697 | 4,09E-13 | 9,74E-11 |
| 12 | CCNA_03904 | hypothetical protein | -1,511387377 | 9,65E-14 | 2,63E-11 |
| 13 | CCNA_01353 | 3-phytase/6-phytase | -1,488911056 | 1,47E-12 | 2,94E-10 |
| 14 | CCNA_01858 | transcriptional regulator, AraC family | -1,367325688 | 3,09E-10 | 3,68E-08 |

| | | | | | |
|---|---|---|---|---|---|
| *15* | *CCNA_00028* | *TonB-dependent receptor* | *-1,280932291* | *2,80E-10* | *3,56E-08* |
| 16 | CCNA_R0143 | small non-coding RNA | -1,071255395 | 5,40E-06 | 0,000266891 |
| 17 | CCNA_00930 | riboflavin synthase subunit alpha | -1,047615917 | 6,10E-12 | 1,11E-09 |
| *18* | *CCNA_03372* | *bacterioferritin-associated ferredoxin* | *-1,046973822* | *4,64E-10* | *5,20E-08* |
| 19 | CCNA_01855 | iron/manganese superoxide dismutase | -1,006894907 | 2,52E-06 | 0,000143207z |
| 20 | CCNA_03155 | transporter | -0,990933278 | 1,67E-06 | 9,79E-05 |
| 21 | CCNA_02999 | hypothetical protein | -0,953064967 | 3,45E-07 | 2,43E-05 |
| 22 | CCNA_00931 | GTP cyclohydrolase II/3,4-dihydroxy-2-butanone-4-phosphate synthase | -0,94878507 | 3,38E-11 | 5,36E-09 |
| 23 | CCNA_01853 | hypothetical protein | -0,939666602 | 4,88E-07 | 3,32E-05 |
| 24 | CCNA_02472 | cobalt-zinc-cadmium resistance protein czcB | -0,93862543 | 1,31E-07 | 1,04E-05 |
| 25 | CCNA_02471 | cobalt-zinc-cadmium resistance protein czcC | -0,937075698 | 2,11E-07 | 1,55E-05 |
| 26 | CCNA_02452 | hypothetical protein | -0,935024241 | 2,43E-06 | 0,000140408 |
| 27 | CCNA_00929 | diaminohydroxyphosphoribosylaminopyrimidine deaminase/5-amino-6-(5-phosphoribosylamino)uracil reduct | -0,928472758 | 6,42E-09 | 6,43E-07 |
| 28 | CCNA_02743 | TonB dependent receptor | -0,914436815 | 1,37E-06 | 8,58E-05 |
| 29 | CCNA_03157 | hypothetical protein | -0,901721215 | 1,90E-05 | 0,000793458 |
| 30 | CCNA_02493 | 3-carboxy-cis,cis-muconate cycloisomerase | -0,886076335 | 1,91E-05 | 0,000793458 |
| 31 | CCNA_02273 | glutathione peroxidase | -0,881452184 | 2,76E-06 | 0,000154841 |
| 32 | CCNA_02494 | 3-oxoadipate enol-lactonase/4-carboxymuconolactone decarboxylase | -0,8500248 | 4,78E-06 | 0,000245929 |
| 33 | CCNA_03156 | hypothetical protein | -0,844762764 | 9,65E-05 | 0,003533781 |
| 34 | CCNA_01361 | hypothetical protein | -0,817658395 | 6,62E-06 | 0,000315296 |
| 35 | CCNA_03158 | iron-sulfur cluster assembly/repair protein ApbE | -0,811475272 | 0,000101595 | 0,00361659 |
| 36 | CCNA_00932 | 6,7-dimethyl-8-ribityllumazine synthase | -0,808751533 | 5,45E-08 | 4,76E-06 |
| 37 | CCNA_01362 | two-component response regulator, UrpR | -0,803688315 | 1,44E-05 | 0,000651024 |
| 38 | CCNA_03023 | TonB-dependent receptor | -0,742348942 | 0,000333024 | 0,010312929 |
| 39 | CCNA_02272 | hypothetical protein | -0,737986183 | 7,53E-05 | 0,002811464 |
| 40 | CCNA_01585 | TonB-dependent receptor | -0,728996795 | 0,001099798 | 0,026336972 |
| *41* | *CCNA_02727* | *PhoH protein* | *-0,722558742* | *0,000434815* | *0,013041024* |
| 42 | CCNA_02690 | hypothetical protein | -0,718862968 | 7,74E-07 | 5,00E-05 |
| 43 | CCNA_02699 | guanine deaminase | -0,69110443 | 5,95E-06 | 0,000287021 |
| 44 | CCNA_R0041 | tRNA Lys | -0,663738585 | 0,000286803 | 0,009103601 |
| 45 | CCNA_03239 | spermidine/putrescine-binding protein | -0,653249387 | 7,33E-05 | 0,002768562 |
| 46 | CCNA_01400 | nitrogen regulatory protein GlnK | -0,65021694 | 9,97E-05 | 0,00361659 |
| ***47*** | ***CCNA_03703*** | ***cytidylate kinase*** | ***-0,647363891*** | ***1,54E-06*** | ***9,38E-05*** |
| 48 | CCNA_02806 | nickel-cobalt-cadmium resistance protein nccC | -0,611848204 | 0,000461825 | 0,013548718 |
| 49 | CCNA_01856 | ferrichrome-iron receptor | -0,599934241 | 0,001643947 | 0,0361762 |
| 50 | CCNA_03697 | cyclase/dehydrase | -0,585902267 | 0,000467166 | 0,013583485 |

Rows in grey indicate genes with $\log_2$ FC < -1. Rows in grey indicate genes with $\log_2$ FC > 1. Genes highlighted in **bold** were found in Tn-Seq (Table S2). Genes highlighted in yellow were found in RNA-Seq_WT (Table S3). Genes highlighted in *italics* were found in ChIP-Seq_WT (Table S5 or S6).

**Table S5. ChIP-seq hits with KdpE antibodies in Δ*kdpE* Δ*xylX* $P_{xyl}$::*kdpE*.**

| Top hit | Peak coordinates | | Reads | Gene(s) upstream | Description gene upstream | Gene(s) down-stream | Description gene downstream |
|---|---|---|---|---|---|---|---|
| | Start | End | | | | | |
| 1 | 1782145 | 1782805 | 0,0404 | CCNA_01661 | hypothetical protein | **CCNA_01663-67** | **potassium-transporting ATPase A chain; potassium-transporting ATPase B chain; potassium-transporting ATPase C chain; potassium-transporting ATPase D chain; potassium-transporting ATPase E chain** |
| 2 | 947332 | 948171 | 0,03514 | CCNA_00869 | DNA modification methyltransferase-related protein | NA | NA |
| 3 | 1159186 | 1159948 | 0,03275 | CCNA_01058 | transcriptional regulator | CCNA_01059 | S-layer protein rsaA |
| 4 | 3944386 | 3945087 | 0,03088 | CCNA_03779 | cytosolic protein | CCNA_03778 | integral membrane protein |
| 5 | 3483833 | 3484620 | 0,02577 | **CCNA_03306** | **SSU ribosomal protein S12P** | CCNA_03308 | hypothetical protein |
| 6 | 4034558 | 4035564 | 0,02439 | CCNA_03872 | tRNA (5-carboxymethylamino methyl-2-thiouridylate) synthase | NA | NA |
| 7 | 3214639 | 3215202 | 0,02346 | CCNA_03063 | hypothetical protein | NA | NA |
| 8 | 2293937 | 2294452 | 0,02334 | CCNA_02137 | endoglucanase H | NA | NA |
| 9 | 2589539 | 2590102 | 0,02322 | CCNA_02448 | hypothetical protein | CCNA_02449 | hypothetical protein |
| 10 | 3882478 | 3883020 | 0,02303 | **CCNA_R0090** | **tRNA Leu** | CCNA_03718 | NADH-ubiquinone oxidoreductase subunit |
| 11 | 382496 | 383029 | 0,02238 | NA | NA | CCNA_00366 | phosphonates transport ATP-binding protein phnC |
| 12 | 3946088 | 3946602 | 0,02175 | CCNA_03780 | hypothetical protein | CCNA_R0092 | Minimal medium sRNA |
| 13 | 1360966 | 1361449 | 0,02148 | CCNA_01233 | haloalkane dehalogenase | CCNA_01234 | metal-dependent hydrolase |
| 14 | 2656060 | 2656622 | 0,02101 | CCNA_R0062 | tRNA Ser | CCNA_02507 | polyisoprenylphosphate hexose-1-phosphotransferase hfsE |

| | | | | | | | |
|---|---|---|---|---|---|---|---|
| 15 | 271358 | 271833 | 0,01815 | CCNA_00259 | hypothetical protein | NA | NA |
| 16 | 1231248 | 1231721 | 0,01712 | CCNA_01127 | hypothetical protein | NA | NA |
| 17 | 51962 | 52413 | 0,01659 | **CCNA_00049** | **hipB HTH transcriptional regulator** | NA | NA |
| 18 | 2867876 | 2868424 | 0,01546 | CCNA_02707 | hypothetical protein | CCNA_02708 | Rrf2 family protein |
| 19 | 370230 | 370667 | 0,01508 | NA | NA | CCNA_00354 | hypothetical protein |
| 20 | 3830048 | 3830494 | 0,01443 | CCNA_03669 | outer membrane protein | CCNA_R0081 | Cell cycle sRNA |
| 21 | 528731 | 529182 | 0,01412 | NA | NA | CCNA_00515 | 5-methyltetrahydropteroyltriglutamate--homocysteine methyltransferase |
| 22 | 2374465 | 2374877 | 0,01349 | CCNA_02226 | glyoxalase family protein | NA | NA |
| 23 | 48733 | 49160 | 0,0124 | **CCNA_00044** | **cytosolic protein** | NA | NA |
| 24 | 1443150 | 1443536 | 0,01237 | NA | NA | CCNA_01328-29 | SSU ribosomal protein S13P; SSU ribosomal protein S11P |
| 25 | 2097129 | 2097566 | 0,01237 | **CCNA_01953** | **hypothetical protein** | CCNA_01954 | ribonuclease E |
| 26 | 3903963 | 3904375 | 0,01221 | NA | NA | CCNA_03735 | transketolase |
| 27 | 1430987 | 1431392 | 0,01191 | NA | NA | **CCNA_01304-08** | **hypothetical protein; SSU and LSU ribosomal proteins S10P, L3P, L1E-L4P, L23P** |
| 28 | 4738 | 5158 | 0,01164 | NA | NA | CCNA_00007 | SSU ribosomal protein S20P |
| 29 | 259263 | 259653 | 0,01071 | CCNA_00247 | Two-component system response regulator SpdR | NA | NA |
| 30 | 3905945 | 3906336 | 0,01016 | CCNA_R0091 | tRNA Ala | **CCNA_03736** | **YGGT family protein** |
| 31 | 211854 | 212213 | 0,00901 | CCNA_00198 | tRNA (guanine-N1) -methyltransferase | CCNA_00199 | hypothetical protein |
| 32 | 898340 | 898686 | 0,00899 | CCNA_00832 | transcriptional regulator, LysR family | CCNA_00833 | Trp repressor binding protein |
| 33 | 564584 | 564941 | 0,00882 | NA | NA | CCNA_00546 | hypothetical protein |
| 34 | 3943983 | 3944347 | 0,00879 | CCNA_03777 | Tas protein | CCNA_03778 | integral membrane protein (2nd peak) |
| 35 | 170660 | 171169 | 0,00858 | NA | NA | **CCNA_00161** | **hypothetical protein** |
| 36 | 3713677 | 3714067 | 0,00857 | **CCNA_03558** | **ATP synthase subunit epsilon** | NA | NA |

| | | | | | | | |
|---|---|---|---|---|---|---|---|
| 37 | 3871400 | 3871739 | 0,00834 | **CCNA_03703** | **cytidylate kinase** | NA | NA |
| 38 | 673942 | 674279 | 0,00826 | **CCNA_00628-27** | **chemotaxis protein cheY; hypothetical protein** | CCNA_00629-32 | methyl-accepting chemotaxis protein; chemotaxis histidine kinase protein cheAII; chemotaxis protein cheW; chemotaxis receiver domain protein cheYII |
| 39 | 75971 | 76311 | 0,00823 | CCNA_00074 | multifunctional fatty acid oxidation complex subunit alpha FadJ | CCNA_R0001 | tRNA-thr |
| 40 | 451606 | 451941 | 0,00821 | NA | NA | **CCNA_00445-50** | **receiver domain-glutamate methylesterase cheBI; chemotaxis receiver domain protein cheYII; chemotaxis protein cheD; cheU protein** |
| 41 | 3701934 | 3702278 | 0,00812 | NA | NA | **CCNA_03545** | **iojap protein family** |
| 42 | 3240296 | 3240644 | 0,00795 | CCNA_03089-90 | hypothetical protein; acetyl-coenzyme A carboxylase carboxyl transferase subunit alpha | CCNA_03091-92 | hypothetical protein NstA; cytochrome P450 IVA5 |
| 43 | 4025020 | 4025362 | 0,00793 | CCNA_03863 | hypothetical protein | CCNA_03865-67 | leucyl-tRNA synthetase; hypothetical periplasmic protein; DNA polymerase III, delta subunit |
| 44 | 3505401 | 3505747 | 0,00791 | CCNA_03324 | hypothetical protein | NA | NA |
| 45 | 1491482 | 1491850 | 0,00786 | CCNA_01377 | hypothetical protein | **CCNA_R0032** | **cell cycle sRNA** |
| 46 | 4035833 | 4036190 | 0,00758 | **CCNA_03873** | **hypothetical protein** | CCNA_03874-75 | Carboxymethylenebutenolidase; quinone oxidoreductase |
| 47 | 3506266 | 3506598 | 0,0075 | **CCNA_03325** | **hypothetical protein** | CCNA_03326 | two-component sensor histidine kinase |
| 48 | 547480 | 547805 | 0,00725 | NA | NA | **CCNA_00536** | **DNA-directed RNA polymerase subunit beta RpoB** |
| 49 | 3487417 | 3487744 | 0,00706 | CCNA_R0076 | tRNA Trp | CCNA_03312 | hypothetical protein |
| 50 | 3505884 | 3506195 | 0,00679 | **CCNA_03325** | **hypothetical protein** | CCNA_03326 | two-component sensor histidine kinase |
| **51** | **2974124** | **2974434** | **0,00668** | **CCNA_02815-16** | **Ice nucleation protein; hypothetical protein** | NA | NA |
| 52 | 792888 | 793205 | 0,00656 | CCNA_00736 | lipoprotein signal peptidase | NA | NA |
| 53 | 338272 | 338581 | 0,00652 | CCNA_00325 | hypothetical protein | CCNA_00326 | vegetatible incompatibility protein HET-E-1 |
| 54 | 45774 | 46091 | 0,00649 | CCNA_00042 | cytosolic protein/LSU ribosomal protein L7AE | NA | NA |

| | | | | | | | |
|---|---|---|---|---|---|---|---|
| 55 | 3836219 | 3836529 | 0,00628 | CCNA_03677 | CoA-transferase family III protein | CCNA_03676 | GNAT acetyltranferase protein |
| 56 | 872140 | 872444 | 0,00605 | **CCNA_00808** | **LSU ribosomal protein L34P** | NA | NA |
| 57 | 844196 | 844494 | 0,006 | CCNA_00783 | secreted protease precursor sapA | CCNA_R0016 | sRNA |
| 58 | 38569 | 38872 | 0,00587 | CCNA_00035-34 | tRNA pseudouridine synthase B; SSU ribosomal protein S15P | NA | NA |
| 59 | 3589483 | 3589783 | 0,00583 | CCNA_03423 | hypothetical protein | CCNA_03424 | AAA-family response regulator tacA |
| 60 | 842145 | 842431 | 0,00522 | CCNA_00782 | aminobenzoyl-glutamate transport protein | **CCNA_R0015** | **tRNA-Val** |
| 61 | 3650680 | 3650964 | 0,00512 | CCNA_03491 | 3-oxoacyl-(acyl-carrier protein) reductase | NA | NA |
| 62 | 299146 | 299438 | 0,00497 | NA | NA | CCNA_00287 | photosensory histidine protein kinase LovK |
| 63 | 3592240 | 3592520 | 0,00474 | CCNA_03426 | hypothetical protein | **CCNA_03427** | **hypothetical protein** |
| 64 | 171437 | 171822 | 0,00445 | NA | NA | CCNA_00162 | hypothetical protein |
| 65 | 2936566 | 2936834 | 0,00422 | CCNA_02777 | vitamin B12 receptor | CCNA_02779 | hypothetical protein |
| 66 | 5419 | 5686 | 0,00417 | NA | NA | CCNA_00008 | chromosomal replication initiator protein DnaA |
| 67 | 3101357 | 3101622 | 0,00407 | NA | NA | CCNA_02941 | transcription elongation factor greA |
| 68 | 3798160 | 3798426 | 0,00407 | **CCNA_03639** | **ferredoxin, 2Fe-2s** | NA | NA |
| 69 | 748304 | 748568 | 0,00398 | CCNA_00691 | ferredoxin | CCNA_00692 | PAS-family GGDEF/EAL protein |
| 70 | 335706 | 335967 | 0,00388 | **CCNA_00320-21** | **LSU ribosomal proteins L27P; L21P** | CCNA_00322 | hypothetical protein |
| 71 | 3527935 | 3528198 | 0,00379 | NA | NA | CCNA_03348 | hypothetical protein |
| 72 | 242803 | 243063 | 0,00374 | CCNA_00224-23 | esterase lipase Family protein; TonB-dependent receptor | CCNA_00225 | hypothetical protein |
| 73 | 2744295 | 2744552 | 0,00369 | CCNA_02597 | hypothetical protein | NA | NA |
| 74 | 3704113 | 3704374 | 0,00351 | NA | NA | CCNA_03548 | carboxy-terminal processing protease precursor |
| 75 | 1637627 | 1637878 | 0,00341 | NA | NA | CCNA_01530 | flagellin |
| 76 | 507753 | 508001 | 0,00337 | CCNA_00489 | phosphoribosyl-AMP cyclohydrolase | CCNA_00490 | hypothetical protein |
| 77 | 3594033 | 3594286 | 0,00332 | CCNA_03430 | LSU ribosomal protein L36P | CCNA_03431 | membrane-bound lytic murein transglycosylase B |

| | | | | | | | |
|---|---|---|---|---|---|---|---|
| 78 | 3958555 | 3958804 | 0,00332 | NA | NA | CCNA_03795 | tellurium resistance protein terB |
| 79 | 101671 | 101918 | 0,00313 | CCNA_00089 | MHYT/PAS-family GGDEF/EAL protein | CCNA_00090 | UDP-glucose 4-epimerase |
| 80 | 3547354 | 3547599 | 0,00313 | CCNA_03372 | bacterioferritin-associated ferredoxin | **CCNA_03373** | **thiamin-phosphate pyrophosphorylase** |
| 81 | 2335782 | 2336026 | 0,00308 | NA | NA | **CCNA_02185-86** | **acetolactate synthase large subunit; acetolactate synthase small subunit** |
| 82 | 2666049 | 2666300 | 0,00303 | CCNA_02516 | glutathione S-transferase | CCNA_02517 | hypothetical protein |
| 83 | 3557028 | 3557275 | 0,00284 | CCNA_03385 | cytosolic protein | CCNA_03384 | LSU ribosomal protein L31P |
| 84 | 3198415 | 3198652 | 0,00283 | CCNA_03043 | type IV pilin protein pilA | CCNA_03044 | CpaC-related secretion pathway protein |
| 85 | 32075 | 32313 | 0,00279 | CCNA_00028 | TonB-dependent receptor | CCNA_00029 | lysine exporter protein |
| 86 | 1951484 | 1951722 | 0,00279 | CCNA_01825 | hypothetical protein | CCNA_01826 | vitamin B12 receptor |
| 87 | 1233328 | 1233564 | 0,00261 | NA | NA | CCNA_01129 | flagellar biosynthesis protein FliQ |
| 88 | 3803931 | 3804165 | 0,00261 | CCNA_03645 | phosphohydrolase (MutT/nudix family protein) | NA | NA |
| 89 | 2889646 | 2889874 | 0,00227 | CCNA_02726 | putative acetyltransferase/acyltransferase | CCNA_02727 | PhoH protein |
| 90 | 1026769 | 1026994 | 0,00218 | NA | NA | CCNA_00950 | flagellar M-ring protein FliF |
| 91 | 4042586 | 4042812 | 0,00218 | NA | NA | NA | NA |
| 92 | 2068013 | 2068237 | 0,00208 | CCNA_01922-20 | SAM-dependent methyltransferase; TPR repeat containing protein; transcription-repair coupling factor | CCNA_01923 | ATP-dependent RNA helicase |
| 93 | 757893 | 758110 | 0,0018 | CCNA_00698 | hypothetical protein | CCNA_00699 | LSU ribosomal protein L28P |
| 94 | 242528 | 242743 | 0,00171 | CCNA_00224-23 | esterase lipase Family protein; TonB-dependent receptor | CCNA_00225 | hypothetical protein |
| 95 | 348344 | 348562 | 0,00156 | NA | NA | CCNA_00338 | TonB-dependent receptor |
| 96 | 659895 | 660107 | 0,00156 | CCNA_00616 | transcriptional regulator | CCNA_00617 | arginine N-succinyltransferase, subunit beta |
| 97 | 285309 | 285578 | 0,00137 | NA | NA | **CCNA_00272-73** | **RmuC family protein; peptide deformylase** |
| 98 | 3876823 | 3877030 | 0,00133 | CCNA_03711-10 | ribosome-associated factor Y; nitrogen regulatory EIIA_Ntr protein | NA | NA |
| 99 | 3415806 | 3416011 | 0,00123 | NA | NA | CCNA_03248 | TonB-dependent receptor |

| | | | | | | | |
|---|---|---|---|---|---|---|---|
| 100 | 4036227 | 4036433 | 0,00123 | **CCNA_03873** | **hypothetical protein** | CCNA_03874-75 | Carboxymethylenebutenolidase; quinone oxidoreductase |
| 101 | 3965156 | 3965367 | 0,00114 | NA | NA | NA | NA |
| 102 | 4034269 | 4034471 | 0,00109 | CCNA_03872 | tRNA (5-carboxymethylaminomethyl-2-thiouridylate) synthase | NA | NA |
| 103 | 247877 | 248075 | 0,0009 | CCNA_00232 | hypothetical protein | CCNA_00233-34 | UDP-N-acetylglucosamine 4,6-dehydratase FlaA1; WecE-family cell wall biogenesis enzyme |
| 104 | 1544848 | 1545045 | 0,00066 | **CCNA_01427** | **lipoprotein, SmpA/OmlA family** | CCNA_01428 | hypothetical protein |
| 105 | 3573867 | 3574059 | 0,00057 | CCNA_03407 | sodium export permease protein | CCNA_03406 | SSU ribosomal protein S21P |
| 106 | 4033069 | 4033260 | 0,00052 | **CCNA_03871-70** | **glucose inhibited division protein A; methyltransferase gidB** | NA | NA |
| 107 | 2775578 | 2775766 | 0,00043 | CCNA_02626 | D-alanine--D-alanine ligase | NA | NA |
| 108 | 220766 | 220949 | 0,00019 | CCNA_00205 | NAD(P)H oxidoreductase yheR | CCNA_00206 | succinyl-CoA:3-ketoacid-coenzyme A transferase subunit A |
| 109 | 840089 | 840271 | 0,00014 | CCNA_00780 | N-formylglutamate deformylase | CCNA_R0014 | Stat phase sRNA |
| 110 | 996211 | 996394 | 0,00014 | NA | NA | **CCNA_00919** | **hypothetical protein** |

Genes highlighted in **bold** were found in Tn-Seq (Table S2). Genes highlighted in green were identified also in RNA-seq experiments (Tables S3-4). Rows in grey indicate genes only found in the Δ*kdpE* Δ*xylX* $P_{xyl}$::*kdpE*, but not in Δ*kdpDE* Δ*xylX* $P_{xyl}$::*kdpE* (Table S6).

**Table S6. ChIP-seq hits with KdpE antibodies in Δ*kdpDE* Δ*xylX* $P_{xyl}$::*kdpE*.**

| Top hit | Peaks coordinates | | Reads | Gene(s) upstream | Description gene upstream | Gene(s) downstream | Description gene downstream |
|---|---|---|---|---|---|---|---|
| | Start | End | | | | | |
| 1 | 1781634 | 1786927 | 0,2096 | NA | NA | **CCNA_01664-67** | **potassium-transporting ATPase B chain, potassium-transporting ATPase C chain, potassium-transporting ATPase D chain, potassium-transporting ATPase E chain** |
| 2 | 1158766 | 1160726 | 0,0446 | CCNA_01058 | transcriptional regulator | CCNA_01059 | S-layer protein rsaA |
| 3 | 271156 | 272038 | 0,0232 | CCNA_00259 | hypothetical protein | NA | NA |
| 4 | 51815 | 52713 | 0,0222 | CCNA_00049 | hipB HTH transcriptional regulator | NA | NA |

| | | | | | | | |
|---|---|---|---|---|---|---|---|
| 5 | 3214428 | 3215249 | 0,021 | CCNA_03063 | hypothetical protein | NA | NA |
| 6 | 528561 | 529343 | 0,0202 | NA | NA | CCNA_00515 | 5-methyltetrahydropteroyltriglutamate--homocysteine methyltransferase |
| 7 | 2374352 | 2374999 | 0,0201 | CCNA_02226 | glyoxalase family protein | NA | NA |
| 8 | 3483614 | 3484591 | 0,017 | **CCNA_03306** | **SSU ribosomal protein S12P** | CCNA_03307 | hypothetical protein |
| 9 | 3505836 | 3506665 | 0,0159 | CCNA_03325 | hypothetical protein | CCNA_03326 | two-component sensor histidine kinase |
| 10 | 2867774 | 2868483 | 0,0153 | CCNA_02707 | hypothetical protein | CCNA_02708 | Rrf2 family protein |
| 11 | 2293858 | 2294484 | 0,0151 | CCNA_02137 | endoglucanase H | NA | NA |
| 12 | 1430933 | 1431696 | 0,015 | NA | NA | **CCNA_01304-08** | **hypothetical protein; SSU and LSU ribosomal proteins S10P, L3P, L1E-L4P, L23P** |
| 13 | 382442 | 383068 | 0,015 | NA | NA | CCNA_00366 | phosphonates transport ATP-binding protein phnC |
| 14 | 1443086 | 1443866 | 0,0139 | NA | NA | CCNA_01328-29 | SSU ribosomal protein S13P; SSU ribosomal protein S11P |
| 15 | 3240242 | 3240999 | 0,0133 | NA | NA | CCNA_03091-92 | hypothetical protein NstA; cytochrome P450 IVA5 |
| 16 | 4034530 | 4035369 | 0,013 | CCNA_03872 | tRNA (5-carboxymethylaminomethyl-2-thiouridylate) synthase | NA | NA |
| 17 | 171271 | 172104 | 0,0129 | NA | NA | CCNA_00162 | hypothetical protein |
| 18 | 2589515 | 2590107 | 0,0129 | CCNA_02448 | hypothetical protein | CCNA_02449 | hypothetical protein |
| 19 | 3946063 | 3946655 | 0,0122 | CCNA_03780 | hypothetical protein | CCNA_R0092 | Minimal medium sRNA |
| 20 | 2656025 | 2656662 | 0,0122 | CCNA_R0062 | tRNA Ser | CCNA_02507 | polyisoprenylphosphate hexose-1-phosphotransferase hfsE |
| 21 | 3882449 | 3883038 | 0,0117 | **CCNA_R0090** | **tRNA Leu** | CCNA_03718 | NADH-ubiquinone oxidoreductase subunit |
| 22 | 3944388 | 3944948 | 0,0108 | CCNA_03779 | cytosolic protein | CCNA_03778 | integral membrane protein |
| 23 | 2097018 | 2097634 | 0,0104 | **CCNA_01953** | **hypothetical protein** | CCNA_01954 | ribonuclease E |
| 24 | 170625 | 171253 | 0,0103 | NA | NA | **CCNA_00161** | **hypothetical protein** |
| 25 | 3763427 | 3764187 | 0,0091 | *CCNA_03609-07* | *cell surface antigen Sca2* | NA | NA |

| | | | | | | | |
|---|---|---|---|---|---|---|---|
| 26 | 484160 | 484722 | 0,0087 | CCNA_00470 | O-antigen polymerase | NA | NA |
| 27 | 748153 | 748640 | 0,0077 | CCNA_00691 | ferredoxin | CCNA_00692 | PAS-family GGDEF/EAL protein |
| 28 | 370229 | 370723 | 0,0077 | NA | NA | CCNA_00354 | hypothetical protein |
| 29 | 4024910 | 4025427 | 0,0077 | CCNA_03863 | hypothetical protein | CCNA_03864 | cytosolic protein |
| 30 | 2945607 | 2946164 | 0,0076 | **CCNA_02789** | **transcriptional regulator, Xre family** | CCNA_02790 | RNA polymerase ECF-type sigma factor |
| 31 | 3701913 | 3702411 | 0,0074 | NA | NA | **CCNA_03545** | **iojap protein family** |
| 32 | 947622 | 948166 | 0,0073 | CCNA_00869 | DNA modification methyltransferase-related protein | NA | NA |
| 33 | 673754 | 674368 | 0,0073 | **CCNA_00628-27** | **chemotaxis protein cheY; hypothetical protein** | CCNA_00629-32 | methyl-accepting chemotaxis protein; chemotaxis histidine kinase protein cheAII; chemotaxis protein cheW; chemotaxis receiver domain protein cheYII |
| 34 | 3797884 | 3798428 | 0,0072 | **CCNA_03639** | **ferredoxin, 2Fe-2s** | NA | NA |
| 35 | 48702 | 49188 | 0,0071 | **CCNA_00044** | **cytosolic protein** | NA | NA |
| 36 | 5346 | 5833 | 0,007 | NA | NA | CCNA_00008 | chromosomal replication initiator protein DnaA |
| 37 | 3713607 | 3714072 | 0,006 | **CCNA_03558** | **ATP synthase subunit epsilon** | NA | NA |
| 38 | 455229 | 455667 | 0,0059 | NA | NA | CCNA_00451 | TonB-dependent receptor |
| 39 | 45627 | 46104 | 0,0059 | CCNA_00042 | cytosolic protein/LSU ribosomal protein L7AE | NA | NA |
| 40 | 348251 | 348691 | 0,0057 | NA | NA | CCNA_00338 | TonB-dependent receptor |
| 41 | 564557 | 564988 | 0,0057 | NA | NA | CCNA_00546 | hypothetical protein |
| 42 | 547457 | 547923 | 0,0054 | NA | NA | **CCNA_00536** | **DNA-directed RNA polymerase subunit beta RpoB** |
| 43 | 839993 | 840384 | 0,0052 | CCNA_00780 | N-formylglutamate deformylase | CCNA_R0014 | Stat phase sRNA |
| 44 | 1231262 | 1231682 | 0,005 | CCNA_01127 | hypothetical protein | NA | NA |
| 45 | 211816 | 212222 | 0,0048 | CCNA_00198 | tRNA (guanine-N1) - methyltransferase | CCNA_00199 | hypothetical protein |
| 46 | 4756 | 5173 | 0,0048 | NA | NA | CCNA_00007 | SSU ribosomal protein S20P |
| 47 | 3546391 | 3546811 | 0,0047 | CCNA_03371 | bacterioferritin | CCNA_03370 | acetyltransferase |

| | | | | | | | |
|---|---|---|---|---|---|---|---|
| 48 | 259251 | 259649 | 0,0045 | CCNA_00247 | Two-component system response regulator SpdR | NA | NA |
| 49 | 1790492 | 1790885 | 0,0043 | NA | NA | **CCNA_01668-70** | **sulfate transport system permease protein cysT; cysW; sulfate transport ATP binding protein cysA** |
| 50 | 2936524 | 2936898 | 0,0043 | CCNA_02777 | vitamin B12 receptor | CCNA_02779 | hypothetical protein |
| 51 | 335602 | 335998 | 0,0043 | **CCNA_00320-21** | **LSU ribosomal proteins L27P; L21P** | CCNA_00322 | hypothetical protein |
| 52 | 2974061 | 2974430 | 0,004 | CCNA_02817 | retrotransposon-related protein | CCNA_02818 | hypothetical protein |
| 53 | 3903982 | 3904357 | 0,004 | NA | NA | CCNA_03735 | transketolase |
| 54 | 1187408 | 1187814 | 0,004 | CCNA_01082 | hypothetical protein | **CCNA_01084** | **hypothetical protein** |
| 55 | 3505400 | 3505781 | 0,0039 | CCNA_03324 | hypothetical protein | NA | NA |
| 56 | 1258471 | 1258839 | 0,0039 | NA | NA | CCNA_01155 | TonB-dependent outer membrane receptor |
| 57 | 2890974 | 2891373 | 0,0039 | NA | NA | CCNA_02728 | hypothetical protein |
| 58 | 483445 | 483815 | 0,0038 | NA | NA | NA | NA |
| 59 | 1361047 | 1361412 | 0,0037 | CCNA_01233 | haloalkane dehalogenase | CCNA_01234 | metal-dependent hydrolase |
| 60 | 451604 | 451958 | 0,0037 | NA | NA | **CCNA_00445-50** | **receiver domain-glutamate methylesterase cheBI; chemotaxis receiver domain protein cheYII; chemotaxis protein cheD; cheU; cheYIIII; cheE proteins** |
| 61 | 3871377 | 3871728 | 0,0037 | **CCNA_03703** | **cytidylate kinase** | NA | NA |
| 62 | 3527895 | 3528260 | 0,0037 | NA | NA | CCNA_03348 | hypothetical protein |
| 63 | 338221 | 338584 | 0,0037 | CCNA_00325 | hypothetical protein | CCNA_00326 | vegetatible incompatibility protein HET-E-1 |
| 64 | 2923091 | 2923459 | 0,0036 | NA | NA | CCNA_02760 | hypothetical protein |
| 65 | 872095 | 872450 | 0,0035 | **CCNA_00808** | **LSU ribosomal protein L34P** | NA | NA |
| 66 | 75973 | 76320 | 0,0035 | CCNA_00074 | multifunctional fatty acid oxidation complex subunit alpha FadJ | CCNA_R0001 | tRNA-thr |
| 67 | 792857 | 793211 | 0,0035 | CCNA_00736 | lipoprotein signal peptidase | NA | NA |
| 68 | 1491511 | 1491867 | 0,0033 | CCNA_01377 | hypothetical protein | **CCNA_R0032** | **cell cycle sRNA** |

| | | | | | | | |
|---|---|---|---|---|---|---|---|
| 69 | 659814 | 660161 | 0,0033 | CCNA_00616 | transcriptional regulator | CCNA_00617 | arginine N-succinyltransferase, subunit beta |
| 70 | 3101337 | 3101667 | 0,0032 | NA | NA | CCNA_02941 | transcription elongation factor greA |
| 71 | 1199039 | 1199369 | 0,0032 | CCNA_01093-91 | hypothetical protein; hypothetical protein; peptide methionine sulfoxide reductase | CCNA_01094 | hypothetical protein |
| 72 | 954916 | 955252 | 0,003 | CCNA_00877 | hypothetical protein | CCNA_00878 | ATP-dependent RNA helicase |
| 73 | 2338437 | 2338776 | 0,003 | NA | NA | **CCNA_R0049** | **tmRNA ssrA acceptor RNA 5' fragment** |
| 74 | 179187 | 179525 | 0,003 | CCNA_00166 | HvyA | CCNA_00167 | bis(5 -nucleosyl)-tetraphosphatase (symmetrical) |
| 75 | 1571743 | 1572068 | 0,003 | CCNA_01463 | 3-deoxy-7-phosphoheptulonate synthase | CCNA_01464 | hypothetical protein |
| 76 | 2666012 | 2666340 | 0,0029 | CCNA_02516 | glutathione S-transferase | CCNA_02517 | hypothetical protein |
| 77 | 3487401 | 3487741 | 0,0029 | CCNA_R0076 | tRNA Trp | CCNA_03312 | hypothetical protein |
| 78 | 3676331 | 3676675 | 0,0029 | **CCNA_03518-17** | **cytochrome c oxidase polypeptide II coxB; cytochrome c oxidase polypeptide I coxA** | CCNA_03519-20 | transcriptional regulator, PadR family; hypothetical protein |
| 79 | 2188294 | 2188620 | 0,0029 | **CCNA_R0048** | **tRNA Leu** | NA | NA |
| 80 | 3589470 | 3589792 | 0,0028 | CCNA_03423 | hypothetical protein | CCNA_03424 | AAA-family response regulator tacA |
| 81 | 2628943 | 2629266 | 0,0028 | CCNA_02482 | transcriptional regulator, AraC family | NA | NA |
| 82 | 2090119 | 2090438 | 0,0027 | NA | NA | CCNA_01948 | peptidoglycan-specific endopeptidase, M23 family LdpA |
| 83 | 1644243 | 1644560 | 0,0027 | CCNA_01537 | acetyltransferase | CCNA_01536 | hypothetical protein |
| 84 | 2465130 | 2465450 | 0,0027 | CCNA_02322 | ApaG protein | CCNA_02323 | hypothetical protein |
| 85 | 479468 | 479780 | 0,0026 | NA | NA | CCNA_00466 | glycosyltransferase |
| 86 | 581935 | 582253 | 0,0025 | CCNA_00566 | starvation sensing protein rspA | CCNA_R0008 | tRNA-Arg |
| 87 | 38521 | 38834 | 0,0024 | CCNA_00035-34 | tRNA pseudouridine synthase B; SSU ribosomal protein S15P | NA | NA |
| 88 | 3198389 | 3198691 | 0,0024 | CCNA_03043 | type IV pilin protein pilA | CCNA_03044 | CpaC-related secretion pathway protein |
| 89 | 996159 | 996458 | 0,0023 | NA | NA | **CCNA_00919** | **hypothetical protein** |
| 90 | 1673358 | 1673661 | 0,0023 | NA | NA | CCNA_01557 | uronate isomerase |

| | | | | | | | |
|---|---|---|---|---|---|---|---|
| 91 | 2744263 | 2744559 | 0,0023 | CCNA_02597 | hypothetical protein | NA | NA |
| 92 | 773135 | 773477 | 0,0021 | NA | NA | NA | NA |
| 93 | 481563 | 481861 | 0,002 | **CCNA_00468** | **hypothetical protein** | CCNA_00467 | oligosaccharide translocase/flippase |
| 94 | 3671724 | 3672009 | 0,002 | CCNA_03514 | cytochrome c oxidase assembly protein ctaG | NA | NA |
| 95 | 1524193 | 1524477 | 0,002 | **CCNA_01405** | **cytosolic protein** | NA | NA |
| 96 | 938582 | 938860 | 0,0019 | **CCNA_00860** | **holdfast inhibitor HfiA;** | NA | NA |
| 97 | 1051337 | 1051620 | 0,0019 | NA | NA | CCNA_00974 | OAR protein precursor |
| 98 | 20519 | 20832 | 0,0019 | **CCNA_03911;** CCNA_00021 | **hypothetical protein; putative aerobic-type carbon monoxide dehydrogenase, large subunit CoxL** | CCNA_00019 | molybdopterin-guanine dinucleotide biosynthesis protein A |
| 99 | 626365 | 626644 | 0,0019 | CCNA_00594-93 | hypothetical protein; hypothetical protein | CCNA_00595 | FecR family protein |
| 100 | 1512258 | 1512537 | 0,0019 | CCNA_01394 | uracil DNA glycosylase superfamily protein | NA | NA |
| 101 | 3704098 | 3704376 | 0,0019 | NA | NA | CCNA_03548 | carboxy-terminal processing protease precursor |
| 102 | 3069632 | 3069909 | 0,0019 | CCNA_02911 | hypothetical protein | CCNA_02912 | prolyl 4-hydroxylase, alpha subunit |
| 103 | 844218 | 844494 | 0,0018 | CCNA_00783 | secreted protease precursor sapA | CCNA_R0016 | sRNA |
| 104 | 507750 | 508024 | 0,0018 | CCNA_00489 | phosphoribosyl-AMP cyclohydrolase | CCNA_00490 | hypothetical protein |
| 105 | 2974952 | 2975226 | 0,0018 | CCNA_02819 | hypothetical protein | CCNA_02818 | hypothetical protein |
| 106 | 477414 | 477685 | 0,0017 | NA | NA | CCNA_00464 | hypothetical protein |
| 107 | 1544819 | 1545089 | 0,0017 | CCNA_01427 | lipoprotein, SmpA/OmlA family | CCNA_01428 | hypothetical protein |
| 108 | 2775537 | 2775803 | 0,0017 | CCNA_02626 | D-alanine--D-alanine ligase | NA | NA |
| 109 | 2232305 | 2232570 | 0,0016 | CCNA_02082 | Sec-independent protein translocase protein tatA | NA | NA |
| 110 | 4032557 | 4032820 | 0,0016 | CCNA_03871 | glucose inhibited division protein A | NA | NA |
| 111 | 1191014 | 1191276 | 0,0016 | CCNA_01086 | GTP-binding protein lepA | NA | NA |
| 112 | 3905969 | 3906326 | 0,0015 | CCNA_R0091 | tRNA Ala | CCNA_03736 | YGGT family protein |
| 113 | 898393 | 898650 | 0,0015 | CCNA_00832 | transcriptional regulator, LysR family | CCNA_00833 | Trp repressor binding protein |

| | | | | | | | |
|---|---|---|---|---|---|---|---|
| 114 | 3594015 | 3594271 | 0,0014 | CCNA_03430 | LSU ribosomal protein L36P | CCNA_03431 | membrane-bound lytic murein transglycosylase B |
| 115 | 771357 | 771604 | 0,0013 | CCNA_00716 | hypothetical protein | CCNA_00717 | hypothetical protein |
| 116 | 1188608 | 1188856 | 0,0013 | CCNA_01082 | hypothetical protein | **CCNA_01084** | **hypothetical protein** |
| 117 | 2727710 | 2727954 | 0,0012 | CCNA_02575 | cytosolic protein | CCNA_02576 | phosphoribosylamidoimidazole-succinocarboxamide synthase |
| 118 | 4042519 | 4042799 | 0,0011 | NA | NA | NA | NA |
| 119 | 3624423 | 3624656 | 0,001 | **CCNA_03462** | **glycine dehydrogenase (decarboxylating)** | NA | NA |
| 120 | 3830117 | 3830348 | 0,001 | CCNA_03669 | outer membrane protein | CCNA_R0081 | Cell cycle sRNA |
| 121 | 1026769 | 1026996 | 0,0009 | NA | NA | CCNA_00950 | flagellar M-ring protein FliF |
| 122 | 3573846 | 3574070 | 0,0008 | CCNA_03407 | sodium export permease protein | CCNA_03406 | SSU ribosomal protein S21P |
| 123 | 1951486 | 1951708 | 0,0008 | CCNA_01825 | hypothetical protein | CCNA_01826 | vitamin B12 receptor |
| 124 | 3836259 | 3836480 | 0,0008 | CCNA_03677 | CoA-transferase family III protein | CCNA_03676 | GNAT acetyltranferase protein |
| 125 | 534860 | 535078 | 0,0007 | CCNA_00520 | ribose-phosphate pyrophosphokinase | NA | NA |
| 126 | 480898 | 481119 | 0,0007 | **CCNA_00468** | **hypothetical protein** | CCNA_00467 | oligosaccharide translocase/flippase |
| 127 | 757105 | 757322 | 0,0007 | **CCNA_R0116** ; CCNA_00698 | **small non-coding RNA**;hypothetical protein | NA | NA |
| 128 | 1162236 | 1162453 | 0,0007 | NA | NA | **CCNA_01060** | **type I protein secretion ATP-binding protein RsaD** |
| 129 | 1291555 | 1291774 | 0,0007 | **CCNA_R0028** | **tRNA Lys** | CCNA_01171 | histidine phosphotransferase ShpA |
| 130 | 3882125 | 3882343 | 0,0007 | CCNA_03717 | ribonuclease D | CCNA_03718 | NADH-ubiquinone oxidoreductase subunit |
| 131 | 3566540 | 3566755 | 0,0007 | NA | NA | CCNA_03396 | trypsin-like serine protease, typically periplasmic, contains C-terminal PDZ domain |
| 132 | 757896 | 758109 | 0,0006 | CCNA_00698 | hypothetical protein | CCNA_00699 | LSU ribosomal protein L28P |
| 133 | 2692075 | 2692286 | 0,0006 | **CCNA_02544** | **hypothetical protein** | CCNA_02543 | LSU ribosomal protein L33P |
| 134 | 1894566 | 1894774 | 0,0005 | CCNA_01766 | hypothetical protein | CCNA_01767 | cytochrome C family protein |
| 135 | 1162558 | 1162765 | 0,0005 | NA | NA | **CCNA_01060** | **type I protein secretion ATP-binding protein RsaD** |
| 136 | 436028 | 436231 | 0,0005 | CCNA_00425 | hypothetical protein | CCNA_00426 | very-short-patch-repair endonuclease |

| | | | | | | | |
|---|---|---|---|---|---|---|---|
| 137 | 1161429 | 1161656 | 0,0005 | NA | NA | NA | NA |
| 138 | 842187 | 842391 | 0,0004 | CCNA_00782 | aminobenzoyl-glutamate transport protein | **CCNA_R0015** | **tRNA-Val** |
| 139 | 1161801 | 1162002 | 0,0004 | NA | NA | NA | NA |
| 140 | 2068008 | 2068238 | 0,0003 | CCNA_01922-20 | SAM-dependent methyltransferase; TPR repeat containing protein; transcription-repair coupling factor | CCNA_01923 | ATP-dependent RNA helicase |
| 141 | 3046614 | 3046812 | 0,0003 | NA | NA | CCNA_02896 | tryptophan halogenase |
| 142 | 1444180 | 1444375 | 0,0002 | CCNA_01331 | hypothetical protein | **CCNA_01330** | **DNA-directed RNA polymerase subunit alpha** |
| 143 | 3547350 | 3547549 | 0,0002 | CCNA_03372 | bacterioferritin-associated ferredoxin | **CCNA_03373** | **thiamin-phosphate pyrophosphorylase** |
| 144 | 2889666 | 2889854 | 0,0002 | CCNA_02726 | putative acetyltransferase/acyltransferase | CCNA_02727 | PhoH protein |
| 145 | 4034294 | 4034486 | $3x10^{-5}$ | CCNA_03872 | tRNA (5-carboxymethylaminomethyl-2-thiouridylate) synthase | NA | NA |
| 146 | 1160778 | 1160958 | $2x10^{-5}$ | NA | NA | NA | NA |

Genes highlighted in **bold** were found in Tn-Seq (Table S2). Genes highlighted in green were identified also in RNA-seq experiments (Tables S3-4). Rows in grey indicate genes only found in the Δ*kdpDE* Δ*xylX* $P_{xyl}$::*kdpE*, but not in Δ*kdpE* Δ*xylX* $P_{xyl}$::*kdpE* (Table S5).

### Supplementary methods

#### Construction of plasmids

pNPTS138-Δ*kdpD* (pHR1121)
Upstream and downstream regions of *C. crescentus kdpD* (*CCNA_01666*) were amplified from NA1000 gDNA by PCR respectively with primers 2002/2003 and 2004/2005. The PCR were then respectively digested with *Hind* III/*Eco* RI and *Eco* RI/*Bam* HI; and ligated into the pNPTS138 vector cut with *Hind* III and *Bam* HI.

pNPTS138-Δ*kdpABC* (pHR1127)
Upstream region of *C. crescentus kdpA* (*CCNA_01663*) and downstream region of *C. crescentus kdpC* (*CCNA_01665*) were amplified from NA1000 gDNA by PCR respectively with primers 1990/1991 and 2000/2001. The PCR were then respectively digested with *Bam* HI/*Eco* RI and *Eco* RI/*Hind* III; and ligated into the pHR253 pNPTS138 cut with *Hind* III and *Bam* HI.

pNPTS138-Δ*kdpE* (pHR1125)
Upstream and downstream regions of *C. crescentus kdpE* (*CCNA_01667*) were amplified from NA1000 gDNA by PCR respectively with primers 2072/2074 and 2073/2075. The PCR were then respectively digested with *Bam* HI/*Eco* RI and *Eco* RI/*Hind* III; and ligated into the pNPTS138 vector cut with *Hind* III and *Bam* HI.

pNPTS138-Δ*kdpDE* (pHR1128)
Upstream region of *C. crescentus kdpC* (*CCNA_01665*) and downstream region of *C. crescentus kdpE* (*CCNA_01667*) were amplified from NA1000 gDNA by PCR respectively with primers 2002/2003 and 2073/2113. The PCR were then respectively digested with *Hind* III/*Eco* RI and *Eco* RI/*Bam* HI; and ligated into the pNPTS138 vector cut with *Hind* III and *Bam* HI.

pNPTS138-Δ*kdpABCDE* (pHR1126)
Upstream region of *C. crescentus kdpA* (*CCNA_01663*) and downstream region of *C. crescentus kdpE* (*CCNA_01667*) were amplified from NA1000 gDNA by PCR respectively with primers 1990/1991 and **2073**/2075. The PCR were then respectively digested with *Bam* HI/*Eco* RI and *Eco* RI/*Hind* III; and ligated into the pHR253 pNPTS138 cut with *Hind* III and *Bam* HI.

pNPTS138-Δ*kefB* (pHR1136)
Upstream and downstream regions of *C. crescentus kefB* (*CCNA_00204*) were amplified from NA1000 gDNA by PCR respectively with primers 2006/2007 and 2008/2009. The PCR were then respectively digested with *Bam* HI/*Eco* RI and *Eco* RI/*Hind* III; and ligated into the pNPTS138 vector cut with *Hind* III and *Bam* HI.

pNPTS138-Δ*kefC* (pHR1141)
Upstream and downstream regions of *C. crescentus kefC* (*CCNA_03611*) were amplified from NA1000 gDNA by PCR respectively with primers 2010/2011 and 2012/2013. The PCR were then respectively digested with *Bam* HI/*Eco* RI and *Eco* RI/*Hind* III; and ligated into the pNPTS138 vector cut with *Hind* III and *Bam* HI.

pNPTS138-Δ*CCNA_01688* (pHR1139)
Upstream and downstream regions of *C. crescentus CCNA_01688* were amplified from NA1000 gDNA by PCR respectively with primers 2018/2019 and 2020/2021. The PCR were then respectively digested with *Bam* HI/*Eco* RI and *Eco* RI/*Hind* III; and ligated into the pNPTS138 vector cut with *Hind* III and *Bam* HI.

pNPTS138-Δ*kup* (pHR1144)
Upstream and downstream regions of *C. crescentus kup* (*CCNA_00130*) were amplified from NA1000 gDNA by PCR respectively with primers 2014/2015 and 2016/2017. The PCR were then respectively digested with *Bam* HI/*Eco* RI and *Eco* RI/*Hind* III; and ligated into the pNPTS138 vector cut with *Hind* III and *Bam* HI.

pNPTS138-$kdpE_{D56E}$ (pHR1258)
Upstream and dowstream regions of *C. crescentus kdpE* D56E were amplified from NA1000 gDNA by PCR respectively with primers 2592/2597 and 2596/2595. Both PCR products were then used in an overlap with primers 2592/2595 and ligated into the pNPTS138 vector cut with *Eco* RV.

pNPTS138-$kdpE_{D56A}$ (pHR1259)
Upstream and dowstream regions of *C. crescentus kdpE* D56A were amplified from NA1000 gDNA by PCR respectively with primers 2592/2594 and 2593/2595. Both PCR products were then used in an overlap with primers 2592/2595 and ligated into the pNPTS138 vector cut with *Eco* RV.

pNPTS138-$kdpD_{H670N}$ (pHR1297)
Upstream and dowstream regions of *C. crescentus kdpE* H670N were amplified from NA1000 gDNA by PCR respectively with primers 2588/2590 and 2589/2591. Both PCR products were then used in an overlap with primers 2588/2591 and ligated into the pNPTS138 vector cut with *Eco* RV.

pNPTS138-$kdpD_{F832L}$ (pHR1521)
Upstream and dowstream regions of *C. crescentus kdpE* F832L were amplified from NA1000 gDNA by PCR respectively with primers 3788/3789 and 3790/3791. Both PCR products were then used in an overlap with primers 3788/3791 and ligated into the pNPTS138 vector cut with *Eco* RV.

pNPTS138-$kup_{A87P}$ (pHR1522)
*C. crescentus kup* A87P was amplified from NA1000 Δ*kdpABCDE* suppressor gDNA by PCR respectively with primers 3270/3271 and ligated into the pNPTS138 vector cut with *Eco* RV.

pNPTS138-$kup_{G253S}$ (pHR1523)
*C. crescentus kup* G253S was amplified from NA1000 Δ*kdpABCDE* suppressor gDNA by PCR respectively with primers 3270/3271 and ligated into the pNPTS138 vector cut with *Eco* RV.

pNPTS138-$kup_{S456R}$ (pHR1524)

*C. crescentus kup* S456R was amplified from NA1000 Δ*kdpABCDE* suppressor gDNA by PCR respectively with primers 3270/3271 and ligated into the pNPTS138 vector cut with *Eco* RV.

pRKlac290-P*kdp*::*lacZ* (pHR1260)
P*kdpA* was amplified from NA1000gDNA by PCR with primers 2171/2172. The The PCR product was then digested with *Kpn* I and *Hind* III and ligated into the pRK290-P*ctrA* vector cut with the same restriction enzymes.

pMR10-P*kdp*::*kdpD* (pHR1296)
P*kdpA*::*kdpD* was amplified from NA1000 Δ*kdpABC* gDNA by PCR with primers 2171/2034 , digested with *Kpn* I/*Pst* I *and* ligated into the pMR10 vector cut with the same restriction enzymes.

pXC-5-P*xyl*::*kdpE* (pHR1525)
*kdpE* was amplified from NA1000 gDNA by PCR primers 2153/2154. The PCR was then digested with *Nde* I and *Sac* I, and ligated into the pHR557 (pXCHYC-5) vector cut with the same restriction enzymes.

pET-28a-*kdpE* (pHR1269)
*kdpE* was amplified from NA1000 gDNA by PCR primers 2153/2154. The PCR was then digested with *Nde* I and *Sac* I, and ligated into the pHR557 (pXCHYC-5) vector cut with the same restriction enzymes.

### Supplementary references.

Casadaban, M. J., & Cohen, S. N. (1980). Analysis of gene control signals by DNA fusion and cloning in *Escherichia coli*. *Journal of molecular biology*, *138*(2), 179-207.

Gober, J.W., and Shapiro, L. (1992). A developmentally regulated *Caulobacter* flagellar promoter is activated by 3' enhancer and IHF binding elements. Mol. Biol. Cell *3*, 913–926.

Skerker, J. M. & Shapiro, L. Identification and cell cycle control of a novel pilus system in Caulobacter crescentus. *EMBO J.* **19**, 3223–3234 (2000).

Thanbichler, M., Iniesta, A. A., & Shapiro, L. (2007). A comprehensive set of plasmids for vanillate-and xylose-inducible gene expression in *Caulobacter crescentus*. *Nucleic acids research*, *35*(20), e137-e137.
